# A weakly structured stem for human origins in Africa

**DOI:** 10.1101/2022.03.23.485528

**Authors:** Aaron P. Ragsdale, Timothy D. Weaver, Elizabeth G. Atkinson, Eileen Hoal, Marlo Möller, Brenna M. Henn, Simon Gravel

## Abstract

While it is now broadly accepted that *Homo sapiens* originated within Africa, considerable uncertainty surrounds specific models of divergence and migration across the continent. Progress is hampered by a paucity of fossil and genomic data, as well as variability in prior divergence time estimates. Here we use linkage disequilibrium and diversity-based statistics, optimized for rapid, complex demographic inference to discriminate among such models. We infer detailed demographic models for populations across Africa, including representatives from eastern and western groups, as well as 44 newly whole-genome sequenced individuals from the Nama (Khoe-San). Despite the complexity of African population history, contemporary population structure dates back to Marine Isotope Stage (MIS) 5. The earliest population divergence among contemporary populations occurs 120-135ka, between the Khoe-San and other groups. Prior to the divergence of contemporary African groups, we infer long-lasting structure between two or more weakly differentiated ancestral *Homo* populations connected by gene flow over hundreds of thousands of years (i.e. a weakly structured stem). We find that weakly structured stem models provide more likely explanations of polymorphism that had previously been attributed to contributions from archaic hominins in Africa. In contrast to models with archaic introgression, we predict that fossil remains from coexisting ancestral populations should be morphologically similar. Despite genetic similarity between these populations, an inferred 1–4% of genetic differentiation among contemporary human populations can be attributed to genetic drift between stem populations. We show that model misspecification explains variation in previous divergence time estimates and argue that studying a suite of models is key to robust inferences about deep history.

## Introduction

Archaeological sites from the Middle Stone Age (approx. 300-40ka) are widely distributed across Africa, and are particularly well represented in the northern, eastern and southern parts of the continent. Similarly, fossils such as those from the sites of Jebel Irhoud, Morocco ^1^, Herto, Ethiopia ^2^ and Klasies River, South Africa ^3^ demonstrate that derived *Homo sapiens* anatomical features were also present across the continent during this period. It has been difficult to reconcile these lines of evidence with evidence from genomics, which have suggested a predominantly tree-like model of recent population divergence from a single ancestral population. It is unclear whether fossil specimens and archaeological sites represent populations which contributed to our ancestors as population precedents, or were local “dead-ends” from which contemporary *Homo sapiens* do not descend. Recently, attempts to reconcile genetic and paleoanthropological data include proposals for a Pan-African origin of *Homo sapiens* by which populations in many regions of the continent contributed to the formation of *Homo sapiens* beginning at least 300ka ^4,5,6^.

Genetic models have been hampered in their contribution to this discussion because they primarily assume (or, at least, have been tested under) a tree-like model of isolation-with-migration. Alternative theoretical scenarios have been proposed, such as stepping stone models ^7^ or population coalescence and fragmentation ^6^, but these approaches are more challenging to interpret and fit to data. However, new population genetic tools now allow for inference involving tens to hundreds of genomes from multiple populations and greater complexity ^8,9,10^. Inspired by evidence for Neanderthal admixture with humans in Eurasia, several recent articles have shown that introducing an archaic hominin ghost population contributing to African populations in the period surrounding the Out-of-Africa migration event substantially improves the description of genetic data relative to single-origin models ^11,12,13,14,9,15,16^. This has driven speculation about the geographic range of this ghost population, possible links to specific fossils, and the possibility of finding ancient DNA evidence (e.g. ^13^). However, these prior articles share two weaknesses. First, they only contrast a single-origin model with an archaic hominin admixture model, leaving out other plausible models ^17^ (Figure 1). Second, they focus on a small subset of African diversity, either because of small sample sizes (2-5 genomes) or because they rely on 1000 Genomes data which only recruited populations of recent West African or Bantu-speaking ancestry (Figure 2C). While ancient DNA from Eurasia has helped us understand early human history outside of Africa, there is no comparably ancient DNA to elucidate early history in Africa ^18^.

**Figure 1:**
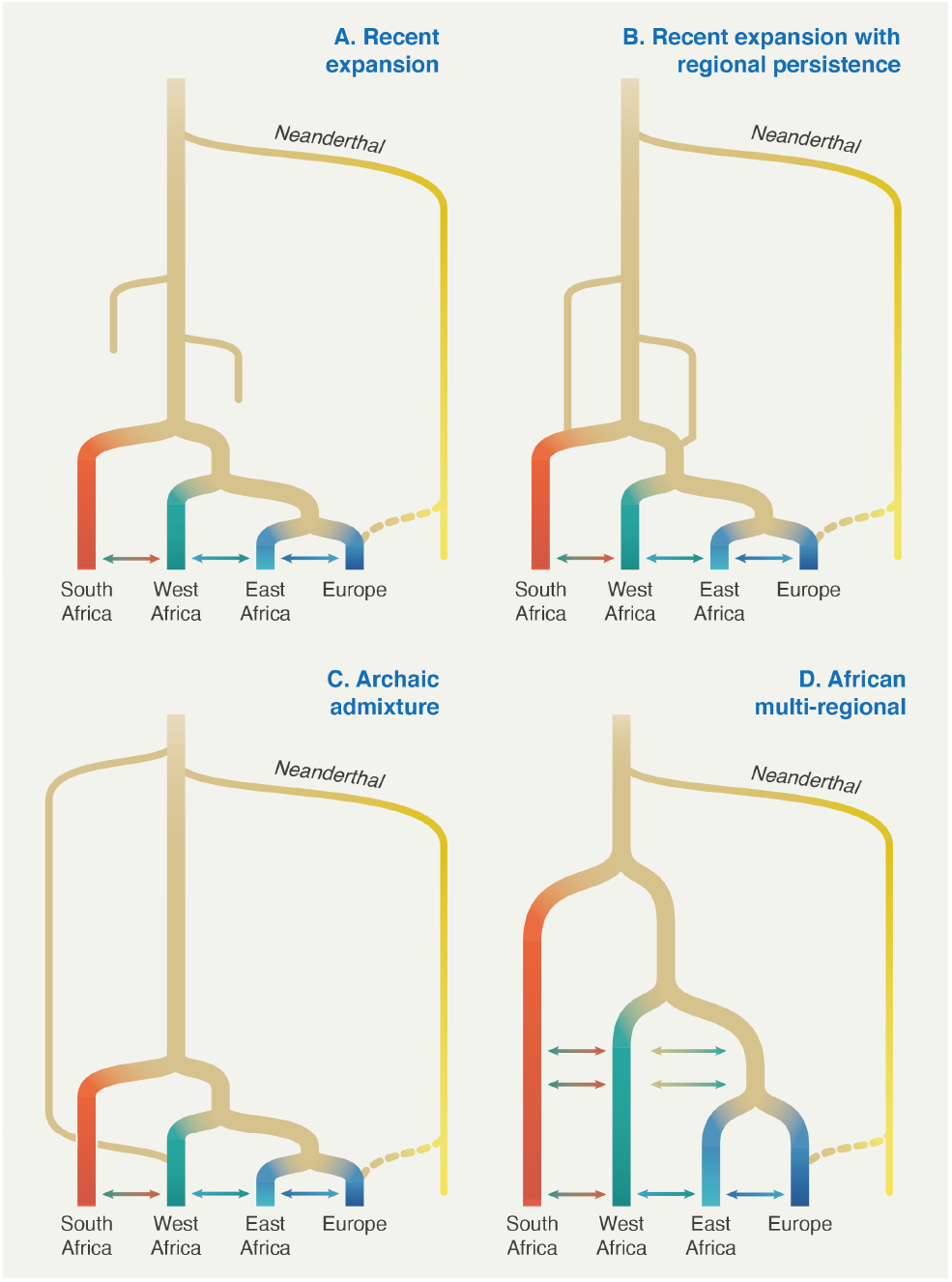
Proposed conceptual models of early human history in Africa. These models have been designed to translate models from the paleoanthropological literature into genetically testable demographic models ^17^. We used these conceptual models as starting points to build detailed parameterized demographic models (Supp. Information section 3) that were then fit to genetic data.

**Figure 2:**
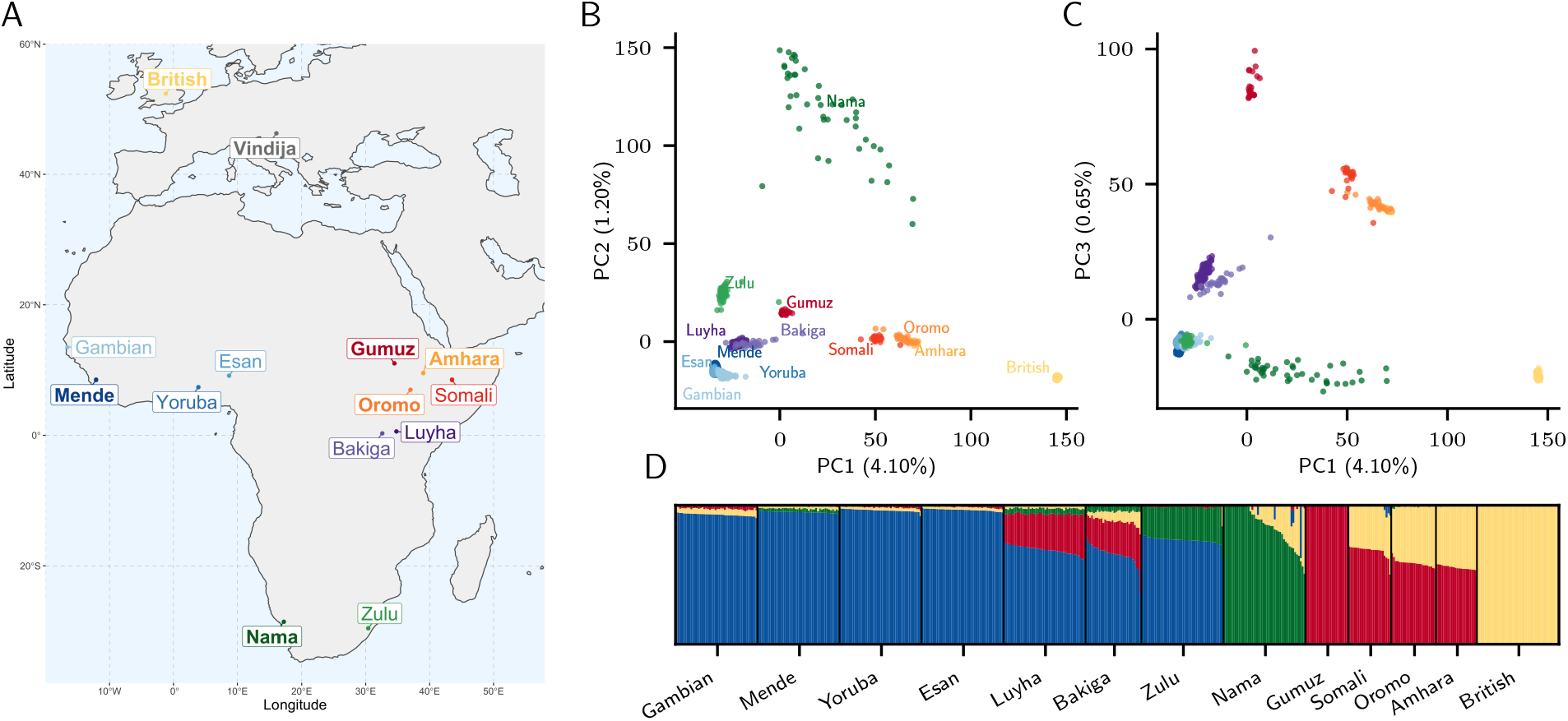
Genetic diversity across Africa. (A) Select populations from the 1000 Genomes and African Diversity Reference Panels illustrate diversity from western, eastern and southern Africa. We chose representatives from each region (bold labels) to build parameterized models, including the newly-sequenced Nama from South Africa, Mende from Sierre Leone, Gumuz, Oromo and Amhara from Ethiopia, and the British and Vindija Neanderthal individual. (B, C) PCA highlights the range of genetic divergence anchored by western Africans, Nama, Gumuz and the British. Percentages show variance explained by each principal component. (D) `ADMIXTURE` with *K* = 4 illustrates signatures of recent gene flow in Africa which reflect colonial-period migration into the Nama, Back-to-Africa gene flow among some Ethiopians, and Khoe-San admixture in the Zulu.

We therefore aim to discriminate among a broader set of demographic models by studying the genomes of contemporary populations. We take as our starting point four classes of models (single population expansion, single population expansion with regional persistence, archaic hominin admixture, and multi-regional evolution, Figure 1), using 290 genomes from southern, eastern, and western Africa as well as Eurasia. By including geographically and genetically diverse populations across Africa, we infer demographic models that explain more features of genetic diversity in more populations than previously reported. These analyses confirm the inadequacy of tree-like models and provide an opportunity to directly evaluate a wide range of alternative models.

## Results

We inferred detailed demographic histories using 4x-8x whole-genome sequencing data for four diverse African populations, comprising the Nama (Khoe-San from South Africa, newly presented here), Mende (from Sierra Leone, MSL from the Phase 3 1000 Genomes Project ^19^), Gumuz (recent descendants of a hunter-gatherer group from Ethiopia ^20,21^), and eastern African agriculturalists (Amhara and Oromo from Ethiopia ^20^). The Amhara and Oromo populations, despite speaking distinct Afro-Asiatic languages, are highly genetically similar ^22,21^ and thus the two groups were combined for a larger sample size (Figure 2). We also included the British (GBR) from the 1000 Genomes Project in our demographic models as a representative source of back-to-Africa gene flow and recent colonial admixture in South Africa. Finally, we used a high-coverage ancient Neanderthal genome from Vindija Cave, Croatia ^23^ to account for gene flow from Neanderthals into non-Africans and gauge the relative time depth of divergence, assuming Neanderthals diverged 550ka from a common stem. We computed one- and two-locus statistics whose expectation within and across populations can be computed efficiently and that are well suited for both low- and high-coverage genomes ^9,24^. Using a maximum-likelihood inference framework, we then fit to these statistics a family of parameterized demographic models that involve population splits, size changes, continuous and variable migration rates, and punctuated admixture events, to learn about the nature of population structure over the past million years.

### A Late Middle Stone Age common ancestry for contemporary humans

We began with a model of geographic expansion from a single ancestral, unstructured source followed by migration between populations, without allowing for contribution from an African archaic hominin lineage or population structure prior to the expansion (Figure 1A). As expected ^9^, this first model was a poor fit to the data qualitatively (Figure S4) and quantitatively (log-likelihood (*LL*) *≈ −*189, 400, Table S2). We next explored a suite of models in which population structure predates the differentiation of contemporary groups, including models allowing for ancestral reticulation (such as fragmentation-and-coalescence or meta-population models, Figure 1B), archaic hominin admixture (Figure 1C), and African multi-regionalism (Figure 1D).

Regardless of the model choice for early epochs, inference of human demographic history for the last 150ka was remarkably robust. The earliest divergence among contemporary human populations differentiates the southern African Nama from other African groups between 110–135ka, with low to moderate levels of subsequent gene flow (Table 1). In none of the high-likelihood models which we explored did the divergence between Nama and other populations exceed *∼*140ka. We conclude that geographic patterns of contemporary *Homo sapiens* population structure date back to the late Middle Stone Age in Africa, likely arising during MIS 5. Although we find evidence for earlier population structure in Africa (see below), contemporary populations cannot be easily mapped onto the more ancient ‘stem’ groups as only a small proportion of drift between contemporary populations can be attributed to drift between stems (Figures 4 and S10–S13).

**Table 1:** Migration and divergence parameters from best fit models. Labeled migration rates correspond to symmetric continuous migration bands shown in Figure 3. Both the continuous migration and merger models inferred a relatively deep split of human stem branches, though they were connected by ongoing migration that maintained their genetic similarity. Bold text indicates migration rates above 10^−4^. In both models, the branch ancestral to the Nama shares a common ancestral population with the other African groups *∼*120–135ka. Following this divergence, the population ancestral to other African groups branches into West and East African groups 60ka. *Migration rates and durations are shown between branches ancestral to 1) Nama and East Africans and their ancestors, and 2) Nama and Mende, respectively. “Divergence times” correspond to the most recent common ancestral population, and does not account for continuous migration or earlier reticulations.

| Likelihood | Label | Population Pair | Divergence Time (ka) | Migration rate per generation | Migration duration (kyr) |
| --- | --- | --- | --- | --- | --- |
| **Continuous Model** |  |  |  |  |  |
| $LL = -115,500$ | a | Stem 1, Stem 2 | 1,163 | 6.43e-5 | 1,028 |
|  | b | Stem 2, Nama | NA | 5.82e-5 | 130 |
|  | c, d | Stem 2, Other Africans* | NA | 3.10e-5, **1.64e-4** | 130, 55 |
|  | e, f | Nama, Other Africans* | 135 | 4.1e-5, 9.8e-6 | 135, 60 |
|  | g | Mende, East Africans | 60 | **2.14e-4** | 60 |
|  | h | East Africans, British | 50 | 4.17e-5 | 50 |
|  | i | Gumuz, Amhara/Oromo | 12 | **3.36e-4** | 12 |
| **Merger Model** |  |  |  |  |  |
| $LL = -102,600$ | a | Stem 1, Stem 2 | 1,442 | **1.16e-4** | 963 |
|  | – | Stem 1S, Stem 1E | 479 | 0 (fixed) | – |
|  | b | Stem 2 to Nama | 119 | **0.71** | pulse |
|  | c | Stem 2 to Stem 1E | 98 | **0.50** | pulse |
|  | d | Stem 2 to Mende | 25 | **0.18** | pulse |
|  | e, f | Nama, Other Africans* | 119 | 4.4e-5, 7.1e-6 | 119, 60 |
|  | g | Mende, East Africans* | 60 | **1.98e-4** | 60 |
|  | h | East Africans, British | 50 | 3.87e-5 | 50 |
|  | i | Gumuz, Amhara/Oromo | 12 | **3.59e-4** | 12 |

Given this consistency in inferred recent history and the numerical challenge of optimizing a large number of parameters, we fixed several parameters related to recent population history so as to focus on more ancient events. Fixed parameters included the time of divergence between western and eastern African populations, set to 60ka, just prior to the split of Eurasians and East Africans set to 50ka. We also fixed the amount of admixture from Neanderthals to Europeans directly following the out-of-Africa migration which was set to 1.5% at 45ka (Supp. Information). These constraints allowed us to integrate information from previous genetic and archaeological research to infer robust migration rates. For example, all models infer relatively high gene flow between eastern and western Africa (*m ≈* 2 *×* 10^−4^, the proportion of migrant lineages per generation). We further find that Back-to-Africa gene flow at the beginning of the Holocene primarily affected the ancestors of the Ethiopian agricultural populations, comprising over half of their genetic ancestry, estimated to be 64–65%. The past 5,000 years also saw major demographic changes, including strong population growth for western Africans as they specialized in yam and oil palm agriculture (estimated 3-fold growth). We observe significant gene flow from the Amhara and Oromo into the Nama, a signal which is likely a proxy for the movement of eastern African caprid and cattle pastoralists ^25,26^, here estimated to constitute a 25% ancestry contribution 2,000 ya. Colonial period admixture from Europeans into the Nama was estimated at 15%, similar to proportions suggested by `ADMIXTURE` (Figure 2).

### Deep but connected population structure within Africa

To account for population structure prior to 135ka, three of our four models allowed for two or more “stem” populations which could diverge either before or after the Neanderthal split. We considered models both with and without migration between these stem populations, and in both cases we tested two different types of gene exchange during the expansion phase, as illustrated in Figure S2: 1) One of the stem population expands (splits into contemporary populations), and the other stem population(s) has continuous symmetric migration with those populations; or 2) one or more of the stem populations expands, with instantaneous pulse (or “merger”) events from the other stem population, so that recent populations are formed by mergers of multiple ancestral populations. Depending on parameter values, this scenario encompasses archaic hominin introgression and fragmentation-and-coalescence models (such as Figure 1B and C). For many parameters, confidence intervals based on bootstrapping are relatively narrow (Tables S2–S6), reflecting an informative statistical approach. Model assumptions have, however, a larger impact on parameter estimates (and thus, real uncertainty). To convey model uncertainty, we highlight features of the two inferred models with high likelihoods. These are referred to as the “multiple-merger” and the “continuous-migration” models. Both allow for migration between stem branches, but differ primarily in the timing of the early divergence of stem populations and their relative *N*_*e*_ (Figure 3). The two models also differ in the mode of divergence during the Middle Stone Age.

**Figure 3:**
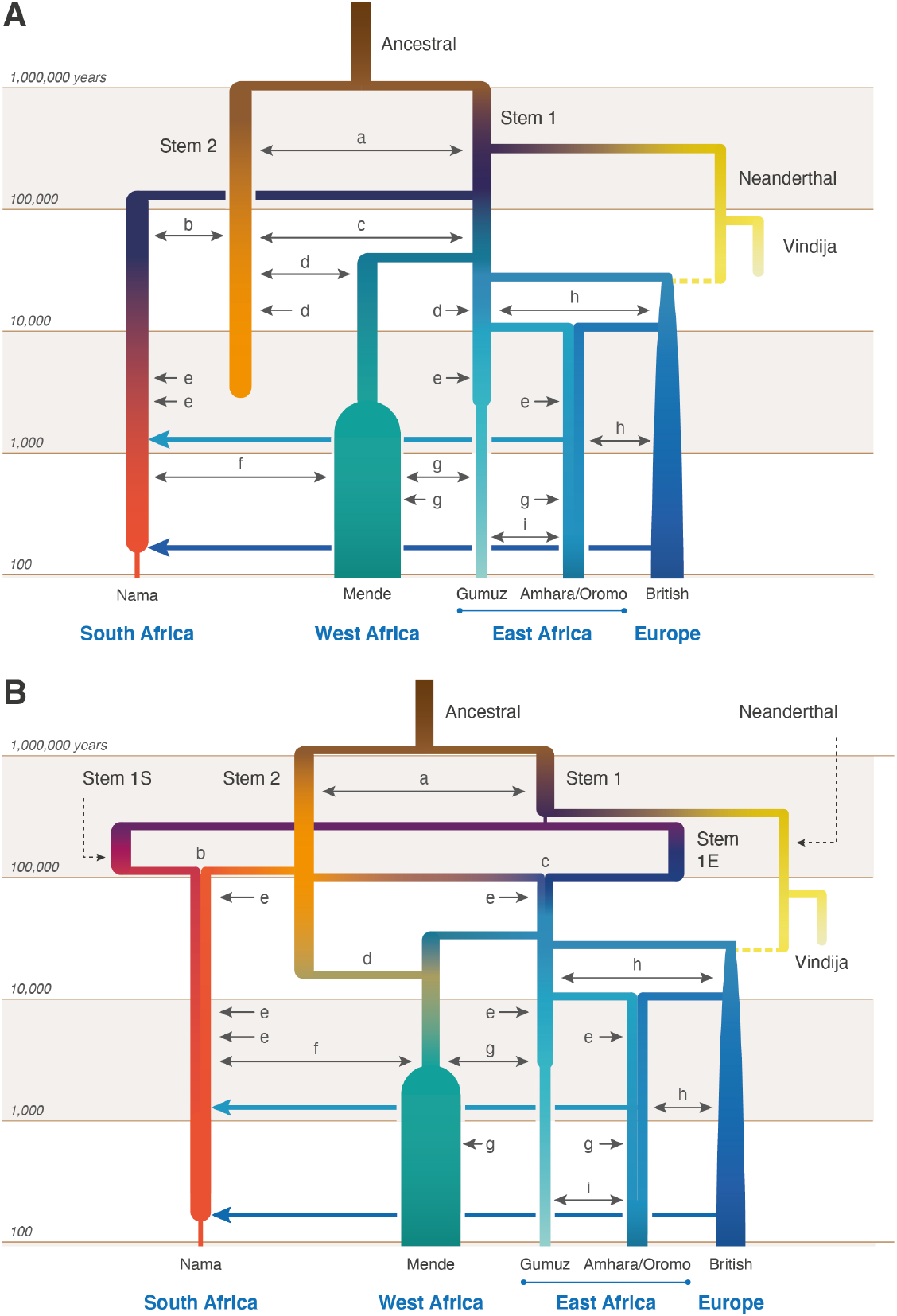
A weakly structured stem best describes two-locus statistics. In the two best-fitting parameterizations of early population structure, continuous migration (A) and multiple mergers (B), models that include ongoing migration between stem populations outperform those in which stem populations are isolated. Most recent populations are connected by continuous, reciprocal migration that are indicated by double-headed arrows (labels matched to migration rates and divergence times in Table 1). These migrations last for the duration of co-existence of contemporaneous populations with constant migration rates over those intervals. The merger-with-stem-migration model (B, with *LL* =*−* 102, 600) outperformed the continuous-migration model (A, with *LL* = *−*115, 500). Colors are used to distinguish overlapping branches and link to Figure 2.

Allowing for continuous migration between the stem populations substantially improved the fits relative to zero migration between stems (*LL≈ −* 102, 600 vs. *−*107, 700 in the merger model and *LL≈ −*115, 500 vs. *−*126, 600 in the continuous migration model). With continuous migration between stems, population structure extends back to 1.1–1.4Ma (Table 1). Migration between the stems in these models is moderate, with a fraction of migrant lineages each generation estimated as *m* = 6.4×10^−5^ 1.2×10^−4^. For comparison, this is similar to inferred migration rates between connected contemporary populations over the past 50ka (Table 1). This ongoing (or at least, periodic) gene flow qualitatively distinguishes these models from previously proposed archaic hominin admixture models (Figure 1C) as the early branches remain closely related, and each branch contributes large amounts to all contemporary populations (Figure 4). Because of this relatedness, only 1% to 4% of genetic differentiation among contemporary populations can be traced back to this early population structure (Supp. Information)

**Figure 4:**
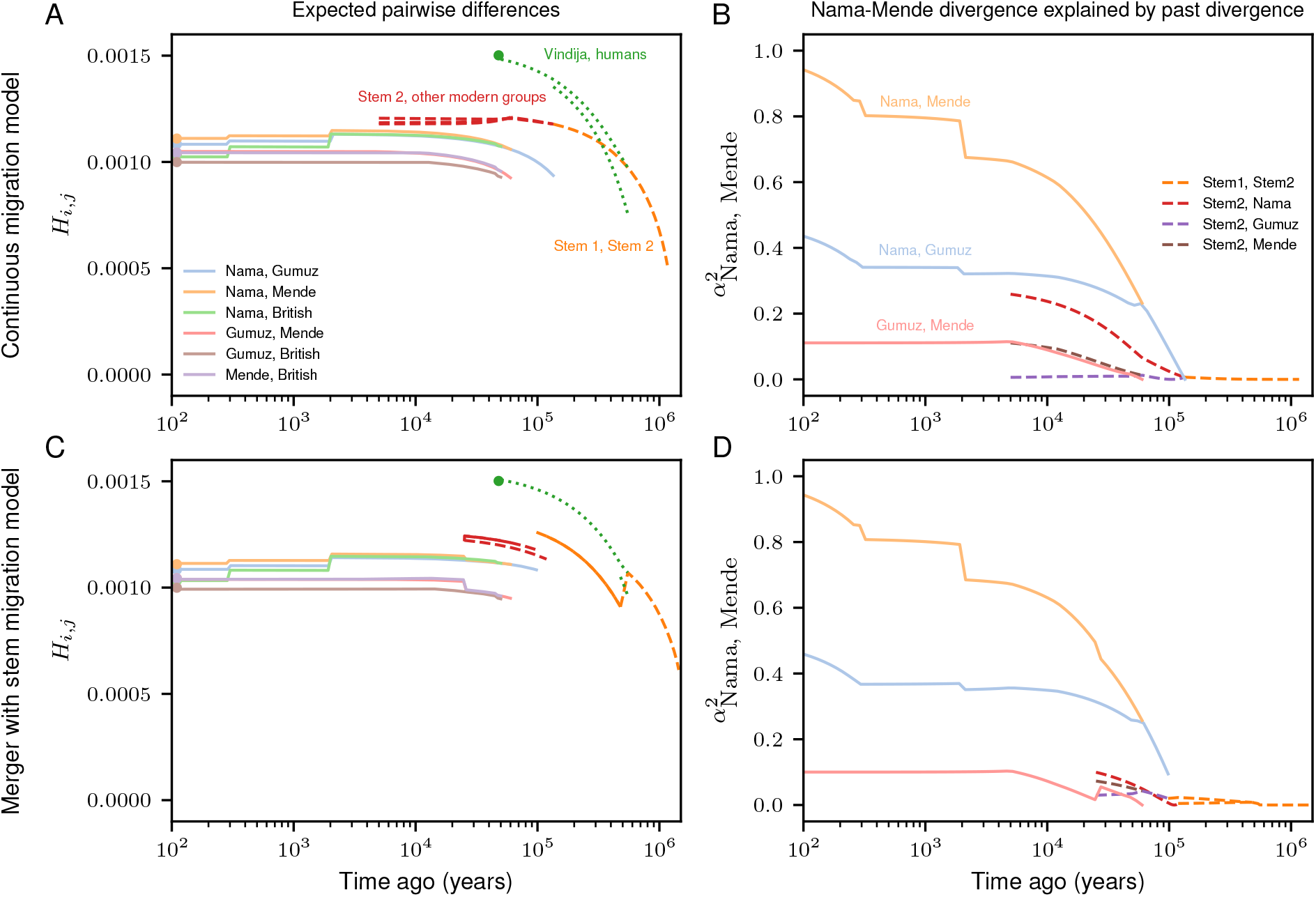
Structure among stems is weak and present-day structure is mostly recent. From the best fit models of our two parameterizations (A and B: continuous migration, C and D: merger with stem migration), we predicted pairwise differences *H*_*i,j*_ between individuals sampled from populations *i* and *j* existing at time *t* (A and C). (B and D) To understand how drift between stems explains contemporary structure, we also computed the proportion *α*^2^ of drift between pairs of sampled contemporary populations that aligns with drift between past populations (here Nama and Mende, see Section S5.2 for details and additional comparisons in Figures S10–S13). Both models infer deep population structure with modest contributions to contemporary genetic differentiation. Most present differentiation dates back to the last 100ka.

Under the continuous-migration model, one of the two stems diverges into lineages leading to contemporary populations in western, southern and eastern Africa, and the other (Stem 2) contributes variable ancestry to those populations. This migration from Stem 2 is highest with the Mende (*m* = 1.6*×* 10^−4^) compared to the Nama and East African populations (*m* = 5.8*×* 10^−5^ and 3.1*×* 10^−5^, respectively), with migration allowed to occur until 5ka. A sampled lineage from the Nama, Mende, and Gumuz have probabilities of being in Stem 2 at the time of Stem 1 expansion (135ka) of approximately 0.145, 0.2, and 0.13, respectively, though these probabilities change over time, precluding the notion of a fixed admixture proportion.

In contrast, under the multiple-merger model, stem populations merge with varying proportions to form the different contemporary groups. We observe a sharp bottleneck in Stem 1 down to *N*_*e*_ = 117 after the split of the Neanderthal branch. This represents the lower bound allowed in our optimization (i.e. 1% of the ancestral *N*_*e*_), although the size of this bottleneck is poorly constrained 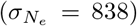. After a long period of exchange with Stem 2, Stem 1 then fractures into “Stem 1E” and “Stem 1S” 479ka. The timing of this divergence was also poorly constrained (*σ*_*T*_ = 166ka). These populations evolve independently until approximately 119ka when Stem 1S and Stem 2 combine to form the ancestors of the Nama, with proportions 29%, 71% respectively. Similarly, Stem 1E and Stem 2 combine in equal proportions (50% each) to form the ancestors of the western Africans and eastern Africans (and thus also all individuals who later disperse during the Out of Africa event). Finally, the Mende receive a large additional pulse of gene flow from Stem 2, replacing 18% of their population 25ka. The later Stem 2 contribution to the western African Mende resulted in better model fits (Δ*LL ≈*60, 000). This may indicate that an ancestral Stem 2 population occupied western or Central Africa, broadly speaking. The differing proportions in the Nama and eastern Africans may also indicate geographic separation of Stem 1S in southern Africa and Stem 1E in eastern Africa.

### Reconciling multiple lines of genetic evidence

Previous studies have found support for archaic hominin admixture in Africa using two-locus statistics ^13,9^, the conditional SFS (cSFS) ^16^, and reconstruction of gene genealogies ^10^. However, none of these studies considered a weakly structured stem. We validated our inferred models with additional independent approaches. We find that the observed cSFS (conditioned on the derived allele being carried in the Neanderthal sample) is very well-described by the merger model (Figures 5A-C and S14–S17), even though this statistic was not used in the fit. Our best-fit models outperform archaic hominin admixture models fit directly to the cSFS (for example, compare to Figure 1 in Durvasula and Sankararaman (2020) ^16^).

**Figure 5:**
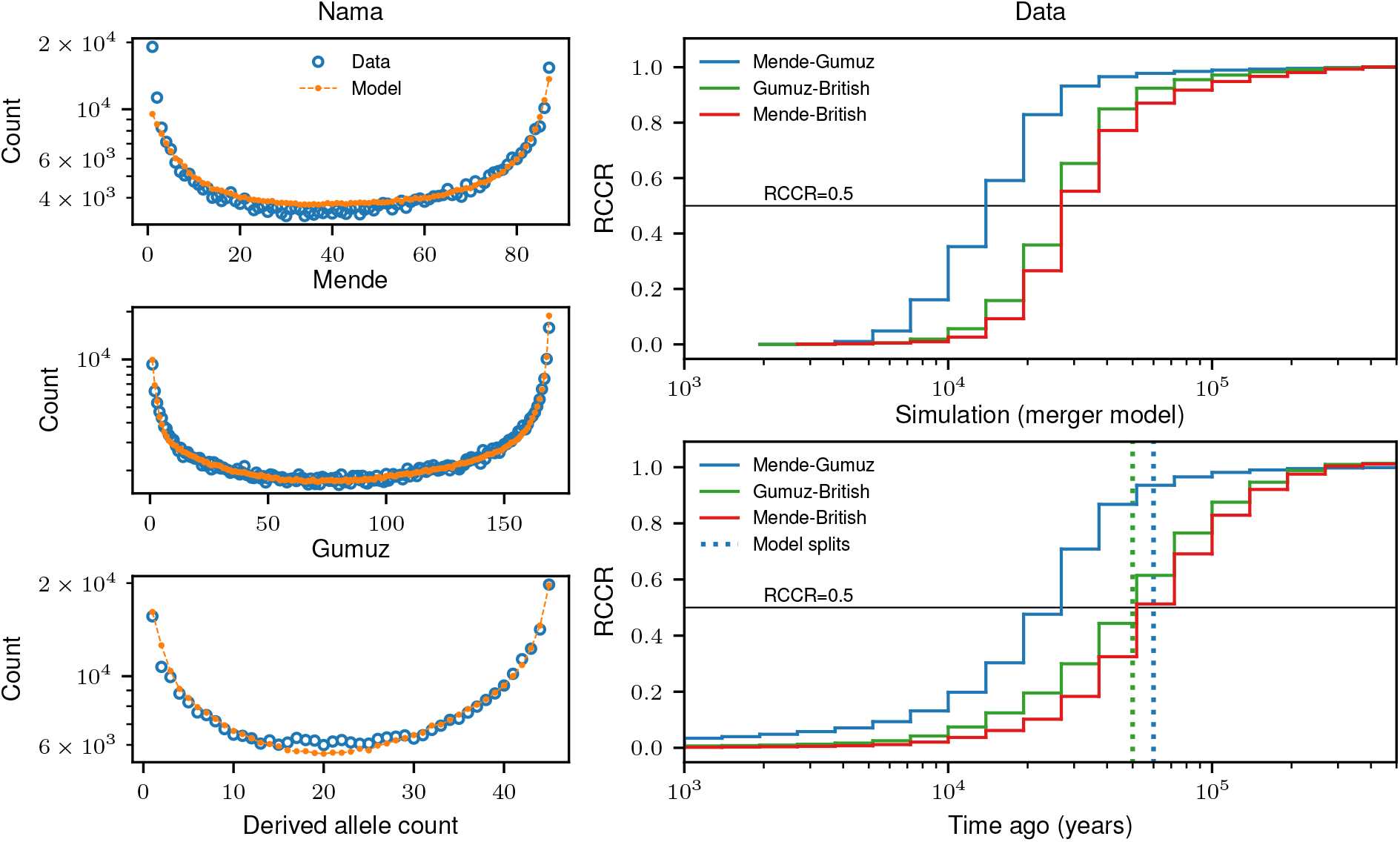
Model validation using independent statistics. (A–C) Using our best fit models to linkage disequilibrium and pairwise diversity statistics, we simulated expected conditional site-frequency-spectra (cSFS) and compared to the observed cSFS from the data. Our inferred models provide a good fit to the data, even though these were not used in our inference. Across the three populations, ancestral state misidentification was consistently inferred to be 1.5–1.7% for intergenic loci (Supp. Information). (D, E) We used Relate ^10^ to reconstruct genome-wide gene genealogies, which we used to estimate coalescence rate trajectories and and cross-coalescence rates between pairs of populations. While coalescence rate distributions are informative statistics about past evolutionary processes, interpretation can be hindered by migration and population structure, and translating relative cross-coalescence rate curves (RCCR) into population divergence times is especially prone to misinterpretation. For example, the Mende-Gumuz comparison shows a more recent increased RCCR than either population with the British, a pattern that is recapitulated under our best-fit model, even though the Mende-Gumuz split occurs prior to the Gumuz-British split.

We used `Relate` ^10^ to infer the distribution of coalescence rates over time in real data and data simulated from our inferred models. Many previous studies have found a reduction of coalescence rates between 1000ka and 100ka in humans, and thus inferred an increase in *N*_*e*_ during the same period ^27^. This increase in inferred *N*_*e*_ could be due to either an increase in population size or to ancestral population structure during the Middle Stone Age ^28^. All models, including the single-origin model, recapitulate an inferred ancestral increase in *N*_*e*_ between 100ka-1Ma (Figure S20). Whereas the single-origin model achieves this by an increase in *N*_*e*_ during that period, the best-fit model recapitulate this pattern without corresponding population size changes.

Relative cross-coalescence rates (rCCR) have recently been used to estimate divergence between a pair of populations, as measured by the rate of coalescence between two groups divided by the mean within population coalescence. Simulations of rCCR accuracy, however, focus on a ‘clean split’ between populations whereby groups diverge without subsequent gene flow. Published estimates of the earliest human divergences with rCCR, which range from 150ka-100ka ^29^, may be significantly biased when compared to more complex models with gene flow as inferred here. We find that midpoint estimates of rCCR are poor estimates for population divergence, often underestimating divergence time by 50% or greater (e.g., Mende vs. Gumuz *∼*15ka compared to a true divergence of 60ka), and recent migration can lead to the misordering of divergence events (Figure 5E). We suggest that rCCR analyses which do not fit multiple parameters including gene flow should be interpreted with caution.

## Discussion

Any attempt at building detailed models of human history is subject to model misspecification. This is true of earlier studies, which often assumed that data inconsistent with a single origin model must be explained by archaic hominin admixture. This is also true of this study. While it remains prohibitive to fully explore the space of plausible models of early human population structure, we sought to capture model uncertainty by exploring multiple parameterizations of early history. The best-fit models presented here include reticulation and migration between early human populations rather than archaic hominin admixture from long-isolated branches. We cannot rule out that more complex models involving additional stems, or hybrid models including both weak structure and archaic hominin admixture may better explain the data. Because parameters related to the split time, migration rates, and relative sizes of the early stems were variable across models, reflecting a degree of confounding among these parameters, we refrained from introducing additional branches associated with more parameters during that period. Rather than interpreting the two stems as representing well-defined and stable populations over hundreds of thousands of years, we interpret the weakly structured stem as consistent with a population coalescence and fragmentation model ^6^. Models including additional diversity within Africa, and early ancient DNA samples from Africa, could further distinguish the archaic hominin admixture model from the weakly-structured-stem model.

### The Middle Stone Age in Africa

By contrast, our inferred models paint a more consistent picture of the late Middle Stone Age as a critical period of change, assuming that estimates from the recombination clock accurately relate to geological chronologies (Supp. Information). During the Middle Stone Age, the multiple merger model indicates three major stem lineages in Africa, tentatively assigned to southern (Stem 1S), eastern (Stem 1E) and western/central Africa (Stem 2). While the length of isolation among the stems is variable across model fits, models with a period of divergence, isolation and then a merger event (i.e. a “reticulation” out-performed models with bifurcating divergence and continuous gene flow.

A population reticulation involves multiple stems contributing genetically to the formation of a group. One way in which this can happen is through the geographic expansion of one or both stems. For example, if during MIS 5, either Stem 1S (Figure 3B) from southern Africa moved northward thereby encountering Stem 2, or Stem 2 moved from central/western Africa southward into Stem 1S – then we could observe disproportionate ancestry contributions from different stems in contemporary groups. We observed two merger events. The first, between Stem 1S and Stem 2, results in the formation of an ancestral Khoe-San population 120ka. The second *≈* 100ka between Stem 1E and Stem 2, results in the formation of the ancestors of East/West Africans as well as later “Out of Africans”. The rapid rise in sea levels and increased precipitation during MIS 5e, following a glacial period of aridity across Africa ^30^, might have triggered migration inland away from the coasts, as has been suggested, e.g., for the Paleo-Agulhas plain ^31^.

Following these merger events, the stems subsequently fracture into subpopulations which then appear to persist over the past *∼*120ka. These subpopulations can be linked to contemporary groups despite subsequent gene flow across the continent; for example, a genetic lineage sampled in the Gumuz has a 0.44 probability of being inherited from the ancestral ‘eastern’ subpopulation (Stem 1E) 150ka versus 0.03 probability of being inherited from the ‘southern’ subpopulation (Stem 1S) and 0.53 probability of being inherited from Stem 2 (see Table S7 for additional comparisons). We also find that Stem 2 continued to contribute to western Africans during the Last Glacial Period, indicative that this gene flow likely occurred in western/central Africa (Table 1).

### Contrasting archaic hominin admixture and a weakly structured stem

Evidence for archaic hominin admixture in Eurasia has bolstered the plausibility of archaic hominin admixture having also occurred in Africa. For this reason, previous work has focused on archaic hominin admixture to explain patterns of polymorphism inconsistent with a single origin model. Here, we have shown that weakly-structured-stem models better capture these patterns. They also help explain an ecological riddle posed by the archaic hominin admixture model. Neanderthal populations were separated from early *Homo sapiens* by thousands of kilometers and continental geographic barriers. By contrast, an archaic hominin population in Africa would need to have stayed in relative reproductive isolation from the ancestral human lineage over hundreds of thousands of years despite closer geographic proximity and reproductive compatibility. The weakly-structured-stem model resolves this ecological riddle by accommodating continuous or recurrent contact between two or more groups present in Africa.

There is evidence for both deleterious and adaptive archaic-hominin-derived alleles in contemporary genomes in the form of a depletion of Neanderthal ancestry in regulatory regions ^32^ or an increased frequency of archaic-hominin-related haplotypes such as at *EPAS1* among Tibetans ^33^. Under previous African archaic hominin admixture models, the estimated 8–10% introgression rate is much higher than Neanderthal gene flow, and would have plausibly been fertile ground for dramatic selection for or against archaic-hominin-derived haplotypes ^34^. By contrast, adaptation under a weakly structured stem would have occurred continuously over much longer periods. Patterns of polymorphism that are inconsistent with the single-stem model predictions have been used to infer putative archaic admixed segments ^11,13,34,16^, negative selection against such segments ^34^, and pervasive positive selection ^35^. However, such approaches are subject to high false positives in the presence of population structure with migration ^32^, and their interpretation should be re-examined in light of a weakly-structured-stem model within Africa.

Multiple studies have shown a correspondence between phenotypic differentiation, usually assessed with measurements of the cranium, and genetic differentiation among human populations and between humans and Neanderthals ^36,37,38^ (see also Section 5.3). This correspondence allows predictions of our model to be related to the fossil record. The fossil record of Africa is sparse during the time period of the stems, but of the available fossils, some are very similar in morphology to contemporary humans (e.g., from Omo Kibish, Ethiopia ^39,40^), others are similar in some morphological features but not others (e.g., from Jebel Irhoud, Morocco ^1,41^), and others are very different in morphology (e.g., from Dinaledi, South Africa ^42,43^). If, as our model predicts, the genetic differences between the stems were comparable to those among contemporary human populations, the most morphologically divergent fossils are unlikely to represent branches that contributed appreciably to contemporary human ancestries.

## Supporting information

Fit models in Demes format

## Acknowledgements

We are grateful for the DNA contribution from each participant which enabled this study; in particular we wish to highlight the generous participation of the Richtersveld Nama community in South Africa and help from local research assistants Willem DeKlerk and Hendrik Kaimann. Additional assistance and community engagement was conducted by Justin Myrick, Chris Gignoux, Caitlen Uren and Cedric Werely. We thank the African Genome Diversity Project for data generation, including Tommy Carensten, Deepti Gurdasani, and Manj Sandhu. We thank Luke Anderson-Trocm’
se for assistance in creating the map in Figure 2. This research was supported by an NIH grant R35GM133531 (to BMH). The content is solely the responsibility of the authors and does not necessarily represent the official views of the National Institutes of Health.

## References

[1] Hublin, J.-J. et al. New fossils from Jebel Irhoud, Morocco and the pan-African origin of Homo sapiens. Nature 546, 289–292 (2017).

[2] White, T. D. et al. Pleistocene Homo sapiens from Middle Awash, Ethiopia. Nature 423, 742–747 (2003).

[3] Deacon, H. J. Two Late Pleistocene-Holocene Archaeological Depositories from the Southern Cape, South Africa. The South African Archaeological Bulletin 50, 121–131 (1995).

[4] Stringer, C. The origin and evolution of Homo sapiens. Philos. Trans. R. Soc. Lond. B Biol. Sci. 371 (2016).

[5] Scerri, E. M. L. et al. Did Our Species Evolve in Subdivided Populations across Africa, and Why Does It Matter? Trends Ecol. Evol. 33, 582–594 (2018).

[6] Scerri, E. M. L., Chikhi, L. & Thomas, M. G. Beyond multiregional and simple out-of-Africa models of human evolution. Nat Ecol Evol 3, 1370–1372 (2019).

[7] Arredondo, A. et al. Inferring number of populations and changes in connectivity under the n-island model. Heredity 126, 896–912 (2021).

[8] Kamm, J., Terhorst, J., Durbin, R. & Song, Y. S. Efficiently inferring the demographic history of many populations with allele count data. J. Am. Stat. Assoc. 115, 1472–1487 (2020).

[9] Ragsdale, A. P. & Gravel, S. Models of archaic admixture and recent history from two-locus statistics. PLoS Genet. 15, e1008204 (2019).

[10] Speidel, L., Forest, M., Shi, S. & Myers, S. R. A method for genome-wide genealogy estimation for thousands of samples. Nat. Genet. 51, 1321–1329 (2019).

[11] Plagnol, V. & Wall, J. D. Possible ancestral structure in human populations. PLoS Genet. 2, e105 (2006).

[12] Hammer, M. F., Woerner, A. E., Mendez, F. L., Watkins, J. C. & Wall, J. D. Genetic evidence for archaic admixture in Africa. Proc. Natl. Acad. Sci. U. S. A. 108, 15123–15128 (2011).

[13] Hsieh, P. et al. Model-based analyses of whole-genome data reveal a complex evolutionary history involving archaic introgression in Central African Pygmies. Genome Res. 26, 291–300 (2016).

[14] Hey, J. et al. Phylogeny Estimation by Integration over Isolation with Migration Models. Mol. Biol. Evol. 35, 2805–2818 (2018).

[15] Lorente-Galdos, B. et al. Whole-genome sequence analysis of a Pan African set of samples reveals archaic gene flow from an extinct basal population of modern humans into sub-Saharan populations. Genome Biol. 20, 77 (2019).

[16] Durvasula, A. & Sankararaman, S. Recovering signals of ghost archaic introgression in African populations. Sci Adv 6, eaax5097 (2020).

[17] Henn, B. M., Steele, T. E. & Weaver, T. D. Clarifying distinct models of modern human origins in Africa. Curr. Opin. Genet. Dev. 53, 148–156 (2018).

[18] Lipson, M. et al. Ancient DNA and deep population structure in sub-Saharan African foragers. Nature (2022).

[19] 1000 Genomes Project Consortium et al. A global reference for human genetic variation. Nature 526, 68–74 (2015).

[20] Gurdasani, D. et al. The African Genome Variation Project shapes medical genetics in Africa. Nature 517, 327–332 (2015).

[21] Gopalan, S. et al. Hunter-gatherer genomes reveal diverse demographic trajectories during the rise of farming in Eastern Africa. Curr. Biol. (2022).

[22] Pagani, L. et al. Tracing the route of modern humans out of Africa by using 225 human genome sequences from Ethiopians and Egyptians. Am. J. Hum. Genet. 96, 986–991 (2015).

[23] Prüfer, K. et al. A high-coverage Neandertal genome from Vindija Cave in Croatia. Science 358, 655–658 (2017).

[24] Ragsdale, A. P. & Gravel, S. Unbiased Estimation of Linkage Disequilibrium from Unphased Data. Mol. Biol. Evol. 37, 923–932 (2020).

[25] Henn, B. M. et al. Y-chromosomal evidence of a pastoralist migration through Tanzania to southern Africa. Proc. Natl. Acad. Sci. U. S. A. 105, 10693–10698 (2008).

[26] Breton, G. et al. Lactase persistence alleles reveal partial East African ancestry of southern African Khoe pastoralists. Curr. Biol. 24, 852–858 (2014).

[27] Li, H. & Durbin, R. Inference of human population history from individual whole-genome sequences. Nature 475, 493–496 (2011).

[28] Mazet, O., Rodríguez, W., Grusea, S., Boitard, S. & Chikhi, L. On the importance of being structured: instantaneous coalescence rates and human evolution—lessons for ancestral population size inference? Heredity 116, 362–371 (2015).

[29] Bergström, A., Stringer, C., Hajdinjak, M., Scerri, E. M. L. & Skoglund, P. Origins of modern human ancestry. Nature 590, 229–237 (2021).

[30] Blome, M. W., Cohen, A. S., Tryon, C. A., Brooks, A. S. & Russell, J. The environmental context for the origins of modern human diversity: A synthesis of regional variability in African climate 150,000–30,000 years ago. J. Hum. Evol. 62, 563–592 (2012).

[31] Marean, C. W. et al. Stone Age people in a changing South African Greater Cape Floristic Region. In Fynbos (Oxford University Press, 2014).

[32] Petr, M., Pääbo, S., Kelso, J. & Vernot, B. Limits of long-term selection against Neandertal introgression. Proc. Natl. Acad. Sci. U. S. A. 116, 1639–1644 (2019).

[33] Zhang, X. et al. The history and evolution of the Denisovan-EPAS1 haplotype in Tibetans. Proc. Natl. Acad. Sci. U. S. A. 118 (2021).

[34] Wall, J. D., Ratan, A., Stawiski, E. & GenomeAsia 100K Consortium. Identification of African-Specific Admixture between Modern and Archaic Humans. Am. J. Hum. Genet. 105, 1254–1261 (2019).

[35] Schrider, D. R. & Kern, A. D. Soft Sweeps Are the Dominant Mode of Adaptation in the Human Genome. Mol. Biol. Evol. 34, 1863–1877 (2017).

[36] Relethford, J. H. Craniometric variation among modern human populations. Am. J. Phys. Anthropol. 95, 53–62 (1994).

[37] Weaver, T. D., Roseman, C. C. & Stringer, C. B. Close correspondence between quantitative-and molecular-genetic divergence times for Neandertals and modern humans. Proc. Natl. Acad. Sci. U. S. A. 105, 4645–4649 (2008).

[38] von Cramon-Taubadel, N. Congruence of individual cranial bone morphology and neutral molecular affinity patterns in modern humans. Am. J. Phys. Anthropol. 140, 205–215 (2009).

[39] Day, M. H. Omo human skeletal remains. Nature 222, 1135–1138 (1969).

[40] Vidal, C. M. et al. Age of the oldest known Homo sapiens from eastern Africa. Nature 601, 579–583 (2022).

[41] Richter, D. et al. The age of the hominin fossils from Jebel Irhoud, Morocco, and the origins of the Middle Stone Age. Nature 546, 293–296 (2017).

[42] Berger, L. R. et al. Homo naledi, a new species of the genus Homo from the Dinaledi Chamber, South Africa. Elife 4 (2015).

[43] Dirks, P. H. et al. The age of Homo naledi and associated sediments in the Rising Star Cave, South Africa. Elife 6 (2017).

