## Supporting Information for "A weakly structured stem for human origins in Africa"

March 14, 2022

**References** **12**

**Supporting tables and figures** **16**

### 1 Data and sequencing

#### 1.1 Sequencing and variant calling

Low coverage (4-8x) Illumina short read data were generated for the Nama, Gumuz, Amhara and Oromo populations as part of the African Diversity Reference Panel (Sanger / Wellcome Trust) (Gurdasani *et al.*, 2015; Pagani *et al.*, 2015) and approved through a secondary data analysis agreement for this project. Briefly, raw reads were aligned to GRCH37 with BWAmem, duplicates marked with Picard MarkDuplicates, reads were realigned around indels with GATK RealignerTargetCreater/IndelRealigner followed by BQSR with dbSNP 137. Contamination checks were performed requiring that FREEMIX $< 0.05$; contamination checks resulted in the elimination of 22 Nama samples. We note that the high heterozygosity in these genomes both due to inherent genetic diversity and admixture may have violated base assumptions for this heterozygosity check. Genomes were then variant called with GATK3.2 Unified Genotyper (DePristo *et al.*, 2011) using joint calling across 2,478 individuals within the ADRP dataset with a minimum base quality of 17. Data were merged with 1000 Genomes Phase 3 (1000 Genomes Project Consortium *et al.*, 2015) using the union of sites identified with bcftools isec (-n+1) (Danecek *et al.*, 2021) then refined with Variant Recalibrator with a truth sensitivity threshold of 99.5%. HapMap III and dbSNP 138 served as known sites while 1000 Genomes Phase 1 Omni2.5 and Phase 1 genomic SNP served as the training set. After VR, no batch effects were observed along PC1 and PC2 for 1000 Genomes vs. ADRP. Phasing on the combined dataset was performed via SHAPEIT2 (Delaneau *et al.*, 2013) and utilized the duoHMM option for duos and trios. We present new whole-genome data for 82 Nama individuals available under accession number EGAD00001006198. Among these 82 Nama samples, we down-sampled the individuals to minimize close relatives and admixture, such that 44 Nama genomes were retained. Second- and third-degrees relatives were inferred from reconstructed pedigrees with Omni2.5 SNP array data. Individuals with $> 70\%$ estimated Khoe-San ancestry were retained for analysis, after partitioning ancestry into $k = 6$ clusters with `ADMIXTURE` (Alexander *et al.*, 2009) where alternative ancestries represent European, West African, Near Eastern, and East African gene flow.

#### 1.2 Nama sample collection and consent

DNA samples were collected from three Nama communities in the Richtersveld region of South Africa, which borders southern Namibia, in 2012. A community guide was present during each interview and facilitated consent in Afrikaans or Nama. Written consent was recorded per our IRB protocol with human subjects approval from Stanford University (Protocol #13829), Stellenbosch University (N11/07/210) and later maintained via SUNY Stony Brook (Protocol #727494). Saliva samples were extracted from Oragene [OGR-500] kits at Stellenbosch University. Results stemming from genetic analyses have been communicated in 2015, 2019 and 2021 via community presentations, a radio interview and to representatives of the Richtersveld National Park.

#### 1.3 Details about populations used in analyses

From the merged dataset of the African Diversity Reference Panel (ADRP) (Gurdasani *et al.*, 2015; Pagani *et al.*, 2015) and the 1000 Genomes Project Consortium *et al.* (2015) phase 3 data, we subsampled to the populations included in this study. From the 1000 Genomes data, these included all populations in the African super-population (ESN, GWD, LWK, MSL, and YRI) and four non-African populations (CEU, GBR, CHB, and PJL). From the ADRP, these included the Amhara, Bakiga, Gumuz, Nama, Oromo, Somali, Nama, and Zulu. This larger set of populations were used in the PCA and `Admixture` analyses. The `Relate` analysis used a slightly smaller set, excluding the Bakiga, Somali, and Zulu samples.

For the demographic modeling, we further subset to the regional populations, from West Africa: Mende from Sierra Leone (MSL); South Africa: Nama from South Africa (newly sequenced); East Africa: Gumuz (traditionally hunter-gatherer with low levels of Eurasian admixture), Oromo and Amhara (combined as traditionally agriculturalists with large proportion of back-to-Africa Eurasian ancestry); and Europe: British (GBR). These were combined with the high-coverage Vindija Neanderthal genome (Prüfer *et al.*, 2017), resulting in 290 total samples used in demographic inference.

##### 1.4 Filtering and subsetting data

All analyses presented in this work focus on biallelic single nucleotide polymorphisms within the 1000 Genomes Phase 3 strict mask. For the `moments-LD` analysis, we focused on intergenic regions because these appear less affected by natural selection compared to both synonymous and nonsynonymous variation (Ragsdale *et al.*, 2018), removing regions that fell outside of the 1000 Genomes strict callability mask. To enable comparison with Neanderthal DNA, we excluded regions for which the Vindija Neanderthal sample had less than 100 contiguous base pairs.

##### 1.5 PCA and ADMIXTURE analysis

We used principal component analysis (PCA) implemented in `scikit-learn` (Pedregosa *et al.*, 2011) and `ADMIXTURE` version 1.3.0 (Alexander and Lange, 2011) to summarize genetic diversity among a broader set of 1000 Genomes and ADRP populations (Figure 2 in the main text). Data filtering was identical for both analyses. We retained biallelic SNPs with no missing data and minor allele frequency greater that 5%. We then thinned the remaining sites to remove SNPs in high linkage disequilibrium with one another. We used the `locate_unlinked` function from `scikit-allel` version 1.2.1 (Miles *et al.*, 2021) to identify SNPs in low linkage disequilibrium, using a threshold of $r^2 = 0.1$, a window size of 100 SNPs, and a step size of 20 SNPs, removing the remaining SNPs. This process was iterated five times for each chromosome, resulting in 327,656 roughly unlinked SNPs used in the PCA and `ADMIXTURE` analyses.

### 2 Computing statistics

#### 2.1 LD and diversity statistics used in model fits

We used multi-population linkage disequilibrium (LD) and pairwise diversity statistics to fit demographic models to data. These statistics, introduced and described in detail in Ragsdale and Gravel (2019), are the multi-population analog of the classic LD statistics first described and computed by Hill and Robertson (1968); Ohta and Kimura (1971).

Given two biallelic loci, with alleles $A$ and $a$ and the left locus and alleles $B$ and $b$ at the right locus, the standard covariance measure of LD is $D = p_{AB}p_{ab} - p_{Ab}p_{aB}$, where $p_{AB}$ is the frequency of $AB$ haplotypes in a population (and thus the probability of drawing an $AB$ haplotype in a random sample of that population). Hill and Robertson (1968) solve for the expectation of $D^2$ using a system of equations that includes $\mathbb{E}[Dz] = \mathbb{E}[D(1-2p_A)(1-2p_B)]$ and $\mathbb{E}[\pi_2] = \mathbb{E}[p_A(1-p_A)p_B(1-p_B)]$, where $p_A$ and $p_B$ are the frequencies of $A$ and $B$ at the left and right loci, respectively. This system also requires the expected pairwise diversity (or expected heterozygosity, assuming random mating), denoted $\mathbb{E}[H] = \mathbb{E}[2p_A(1-p_A)] = \mathbb{E}[2p_B(1-p_B)]$, assuming equal mutation rates at the two loci.

Ragsdale and Gravel (2019) showed how to compute the analogous multi-population set of LD statistics, and we refer readers there for details on their definitions, computation, and interpretations. In short, we obtain expectations of $D^2$ in each population, the cross-population product $D_iD_j$ (where $i$ and $j$ index populations), as well as those additional statistics $Dz$ and $\pi_2$ taken over different combinations of population indexing. That is $Dz_{i,j,k} = D_i(1-2p_{A,j})(1-2p_{B,k})$ and $\pi_{2;i,j,k,l} = p_{A,i}(1-p_{A,j})p_{B,k}(1-2p_{B,l})$. We consider statistics normalized by $\pi_2$ in an arbitrary reference population (throughout, we use the Mende $\pi_2$), which removes any dependence on the mutation rate. Thus, statistics take the form $\sigma^2_{d;i,j} = \frac{\mathbb{E}[D^2_{i,j}]}{\mathbb{E}[\pi_2]}$, $\sigma_{Dz;i,j,k} = \frac{\mathbb{E}[Dz_{i,j,k}]}{\mathbb{E}[\pi_2]}$, and so on. Multi-population pairwise diversity statistics $\mathbb{E}[H_i] = \mathbb{E}[2p_i(1-p_i)]$ and $\mathbb{E}[H_{i,j}] = \mathbb{E}[p_i(1-p_j) + p_j(1-p_i)]$ were also normalized by $H$ in the Mende, so that all pairwise diversity measures are relative to the reference population.

$\sigma_{Dz}$ has been shown to be sensitive to deep population structure and archaic hominin admixture (Ragsdale and Gravel, 2019), and this statistic is closely related to $S^*$ statistics used to scan for introgressed haplotypes (Plagnol and Wall, 2006). Pairwise diversity statistics $H_{i,j}$ have also been widely used in demographic inference involving ancient DNA and many statistics, as $f_2$, $f_3$ and $f_4$ statistics can be expressed as linear combinations of $H_{i,j}$. $f$-statistics form the basis of admixture graph analysis (Lipson,

2020). Therefore, the set of statistics used here encompass multiple features of genetic data that have been used to infer models of archaic hominin admixture and population structure involving many populations.

#### 2.2 Computing LD and diversity statistics

We compared single- and two-locus statistics in the data to predictions based on detailed demographic models. Model predictions were obtained using recursions described in Ragsdale and Gravel (2019) and implemented in the software `moments` (`https://bitbucket.org/simongravel/moments/src/main/`). The model computes expected patterns of single-nucleotide pairwise diversity and linkage disequilibrium as a function of recombination distance between variants within and across populations, under the assumption of neutrality.

For numerical convenience, observed genetic variants were binned by recombination distances. We assessed the robustness of the statistics to errors in the recombination maps by considering two different recombination maps, the OMNI YRI and HapMapII (1000 Genomes Project Consortium *et al.*, 2015; International HapMap Consortium *et al.*, 2007). Statistics were largely unchanged by using a different map (Figure S3), although both maps were inferred using array data, which is sparser than whole-genome sequencing data. In subsequent analyses, we used data based on recombination rates from the OMNI YRI map.

We removed bins of recombination distance less than a recombination distance of $r = 5 \times 10^{-6}$ (at a rough estimate of 1 cM/Mb, this corresponds to a minimum distance of 500 bp on average) to avoid previously reported biases at short distances due to processes like multinucleotide mutations (Harris and Nielsen, 2014; Ragsdale and Gravel, 2019). To avoid uncertainty in phasing, we used unphased genotypes to compute LD statistics, as proposed in Ragsdale and Gravel (2020).

#### 2.3 Estimating two-locus statistics with small sample sizes

The approach from Ragsdale and Gravel (2020) provides unbiased estimates of the LD statistics considered here, which can be directly compared to expectations for those same statistics computed from the multi-population Hill-Robertson system. Estimated statistics for populations with smaller sample sizes have greater uncertainty, which is accounted for by computing variances and covariances via bootstrap over genomic regions, assuming that there is no correlation between distant genomic regions (i.e., no long-range LD).

Some statistics, such as $\mathbb{E}[D_i D_j]$, require at least two diploid samples from a population to compute. Since we used a single Neanderthal sample, such statistics for the Neanderthal population were not used in the fit. By contrast, there are statistics that only require a single sample in a given population to estimate. These include cross-population nucleotide diversity measures, as well as some LD statistics involving more than one population. For example, statistics of the form $D_{\text{human}}(1-2p_{\text{human}})(1-2q_{\text{neanderthal}})$ require a single Neanderthal sample and are informative of the Neanderthal demography. These statistics were included in the fit, but statistics requiring more than one Neanderthal sample to estimate were removed.

#### 2.4 Computing conditional SFS

The conditional site frequency spectrum (or cSFS) is the distribution of allele frequencies restricted to loci that satisfy a given condition. Specifically, we consider the distribution of allele frequencies in present-day populations conditioned on the Vindija Neanderthal carrying the derived allele relative to the inferred ancestral allele. Ancestral alleles alleles were determined from a 6 primate alignment (1000 Genomes Project Consortium *et al.*, 2015). This cSFS is expected to be close to uniform under neutrality and a simple split model (with no subsequent migration) between the ancestors of humans and Neanderthals (Chen *et al.*, 2007). By contrast, a U-shaped distribution has been taken as evidence for introgression from an archaic hominin population that split from humans at least as far back in time as the human-Neanderthal split (Yang *et al.*, 2012; Durvasula and Sankararaman, 2020). Sites with no calls in the Vindija Neanderthal were excluded from this analysis.

Because we were concerned that cSFS analyses may be affected by incorrect inference of the ancestral allele (Hernandez *et al.*, 2007), we computed the cSFS for all mutations, and for transitions and transversions separately (Figures S14–S17). Comparisons of these observed cSFS with model predictions are discussed in the model prediction section below.

### 3 Model specification and fitting

For the early history, we tested model parameterizations that cover many of the proposed scenarios of population structure, size changes, and/or archaic hominin admixture. The simplest model, in terms of number of parameters, was a single-origin expansion of humans, with no structure in the stem and no archaic hominin admixture aside from the Neanderthal admixture in Eurasian populations following the out-of-Africa migration. This model allowed for a population size change in the stem of humans between the ancestral split of the human-Neanderthal lineages and the more recent split of branches leading to southern and western/eastern African populations.

To include population structure in early human history, we considered multiple parameterizations of models that allowed more than a single stem population. In general, stem populations were allowed to vary in their sizes, split times, and migration rates, with parameterizations flexible enough to encompass proposed scenarios of either archaic hominin admixture or population structure, both connected by gene flow or with isolation between stems.

In one parameterization of early structure, which we refer to as a "continuous migration" model, a secondary stem population (stem 2) split from the primary stem (stem 1) that leads to contemporary humans. Stem 1 contributed to present-day populations via a series of population splits similar to the single-origin model, while stem 2 contributed through continuous symmetric migrations with contemporaneous populations. The symmetrical migration rates could differ across population pairs and over different epochs. This continuous migration was allowed until stem 2 disappeared, which occurred as recently as 5ka. We tested models that both allowed or disallowed migration between stems, i.e., before stem 1 split into S/E/W African populations.

In another parameterization of early structure, a secondary stem population (stem 2) contributed ancestry to present-day populations via a series of instantaneous admixture events (i.e., pulse admixture or "merger" events) to lineages leading to sampled present-day African populations. Merger events were allowed to occur in one or more of the Nama, Mende, and Gumuz branches, as well as the branch of East and West Africans prior to their split. Those admixture events were allowed to occur at any time along those branches, and with any proportions, and stem 2 was allowed to split from the primary stem at any time before subsequent divergences (and either before or after the split of the Neanderthal branch). We tested models that allowed migration between the early stems, before subsequent splits and admixture events, as well as models that were restricted to isolation between stem branches. Depending on the specific parameters, such models encompass commonly-considered scenarios for admixture from a ghost archaic hominin population (e.g., if a long-isolated lineage more recently contributes a minority of ancestry to one or more populations), as well as relatively simple fragmentation-coalescence scenarios.

Based on the geographical locations of present-day populations, we tentatively labeled ancestral branches using a parsimony in migration, referring to South, East, and West African branches (S/E/W AFR, Figures 1, 2). We do not know the geographical location of these ancestral populations (nor even if they correspond to populations with a well-defined geographic range), and these labels should be considered as tentative. However, we found it useful to name branches in reference to where in Africa their descendants are found. Parameters were converted to physical units (years, effective population sizes, etc) assuming a human generation time of 29 years.

#### 3.1 General strategy for building models and introducing complexity

With up to six sampled populations in final demographic models that we fit, there are many parameters to learn. Even in the simplest model involving all populations (such as the tree-like single-origin model), there are a few dozen parameters defining split times, migration rates, admixture timings and proportions, and population sizes and size changes. Thus, parameter space for a given model topology is large. In addition, the space of possible model topologies is itself large – as the number of populations increases, the number of possible topologies also increases, as there are more possibilities for the order of divergence and admixture events.

In order to narrow the set of possible models to plausible scenarios and to avoid overfitting, we took an approach that combined the incremental addition of complexity, starting with fewer populations before combining all populations, as well as fixing parameters that have been previously estimated or that fit consistently across all model scenarios. By initially considering sets of two or three populations, we were able to narrow down the relative orders of divergences between African and Eurasian populations. Assuming simple isolation-with-migration models, the Nama appeared to be the earliest diverging population of those we considered, with West African (Mende) and East African (Gumuz) populations diverging more recently, followed by the split of the Eurasian branch from the East African branch.

We performed an initial round of optimization including all six populations with a family of models including single- and multiple-stem scenarios as described in the previous section. We identified parameters that reached consistent values across all models. These included the timings of recent or non-central divergences and admixture events:

- Neanderthal/human split: 550ka
- East/West African population split: 60ka
- East African/European split: 50ka
- Neanderthal introgression to Eurasian branch: 45ka
- Eurasian back-to-African migration: 12ka

When testing multiple variations of the more complex models, we kept these values fixed (see Table S1 for a complete list of all fixed parameters). This reduced the potential for overfitting the more complex models, while reducing the computational cost of optimization.

Our models also included recent events to account for known migrations, admixtures, and growths and declines in effective population sizes. Many of these parameters were fixed based on previous historical, genetic, or anthropological research, namely

- Mende population expansion and Gumuz population decline: 5ka (Gopalan *et al.*, 2022)
- East African pastoralist to South African admixture: 2ka (Uren *et al.*, 2016)
- European to South African admixture: 10 generations ago
- South African population decline following colonial admixture: 9 generations ago

allowed to vary in the fits. The total number of parameters that were ultimately inferred were 21-31, depending on the complexity of the model.

#### 3.2 Optimization using `moments`

`moments-LD` uses a composite likelihood approach to simultaneously fit relative pairwise diversity and LD statistics over a range of recombination distances. Likelihoods were computed independently for pairwise diversity and each recombination distance bin using a multivariate Guassian likelihood function as described in Section 2.2. These were multiplied across bins and the single set of heterozygosity statistics following the approach detailed in Ragsdale and Gravel (2019). For each model tested, we ran multiple rounds of optimization, alternating between the *optimize_log_fmin*, *optimize_log_powell*, and *optimize_lbfgsb* methods to explore parameter space and hone in on the best fit parameters. Initial guesses for parameters were chosen from demographically plausible starting points and then perturbed to explore space, using gradient descent (on the log of the parameters). The best fits from these initial rounds of optimization were then chosen as the starting points for optimization using the Powell and/or the L-BFGS-B methods (as implemented in the SciPy optimization package (Virtanen *et al.*, 2020)). This process was repeated with alternating optimization methods until the best fit parameter set converged consistently.

#### 3.3 Uncertainty and confidence intervals using Godambe methods

To compute likelihoods, we needed an estimate of the joint distribution of summary statistics. We estimated uncertainty due to the finite amount of genetic material used in inference using bootstrap over 500 segments along the genome with roughly equal lengths of retained sequences within each segment. First, for each distance bin, we used these bootstrap samples to obtain a variance-covariances matrix across all statistics. This variance-covariance matrix was used to obtain a model likelihood for each recombination distance bin and single-locus nucleotide diversity, as a multivariate Gaussian likelihood. The full model likelihood was taken as the product of likelihoods over each bin. In other words, we optimized a composite likelihood where observations in different bins were taken to be independent.

For the best fit parameters, we computed confidence intervals using the same set of bootstrapped samples and Godambe Information approach (Coffman *et al.*, 2016), which corrects composite likelihood estimates of confidence intervals to account for nonindependence in the data, including linkage between loci and nonindependence of recombination bins.

Tables S2–S6 give the best fit values for all parameters in the four models considered here, along with standard errors using computed using the Godambe method. Figures S4–S7 show the corresponding fits to many of the LD statistics used in the inference. The $\sigma_{Dz}$ statistics are fit poorly by the single-origin model, while the models with multiple stems all provide reasonable fits to the data. The merger-with-stem-migration model performed the best of these four models, although the fits were nearly visually indistiguishable from the fits from the merger-with-stem-migration model.

### 4 Gene genealogy reconstruction

We used `Relate` version 1.0.16 (Speidel *et al.*, 2019) to reconstruct genome-wide gene genealogies using a combined set of 1000 Genomes and African Diversity Reference Panel datasets, retaining all AFR-labeled populations and GBR, CEU, PJL, and CHB from the 1000 Genomes panel and the Nama, Gumuz, Oromo, and Amhara from ADRP. We used all autosomes and applied the 1000 Genomes Phase 3 strict mask, we used an ancestral sequence determined by a 6-primate alignment (human_ancestor_GRCh37_e59), and we used the HapMap II combined recombination map, all in GRCh37 coordinates. We assumed a generation time of 29 years, a mutation rate of 1.25e-8, and followed the standard pipeline described in the `Relate` documentation.

From the reconstructed gene genealogies, we computed coalescence rates within and between populations using `Relate`'s function `RelateCoalescentRate`. This allows for an estimate of the instantaneous inverse coalescence rate (IICR) for samples drawn within each population (the inverse of which is often interpreted as the effective population size history), and the relative cross coalescence rates between pairs of populations (which are commonly used to estimate divergence times).

Following Speidel *et al.* (2019) we also identified "deep branches" within gene trees. Such a "deep branch" is a branch within a marginal tree (meaning, the tree at a given locus) that has its upper end (i.e., the node of its coalescence with another branch) extending to more than 1 million years in age, and we partitioned deep branches based on their lower node ages (where two or more branches coalesce into this single branch) into bins between 0 and 1Ma. Such branches can be categorized by their association with Neanderthals by comparing mutations that fall upon such a branch to the allele found in a Neanderthal genome sequence. For this, we used the published high-coverage Vindija Neanderthal (Prüfer *et al.*, 2017). Again following the analysis in Speidel *et al.* (2019), if one or more mutations on a deep branch (carrying at least two derived mutations) are shared with the Neanderthal sequence, the branch is inferred to have passed through the Neanderthal lineage.

A deep branch is assigned to a contemporary population if at least one sample from that population has ancestry that passes through that branch. Speidel *et al.* (2019) show that deep branches with lower ends more recent than the Neanderthal introgression event are enriched for Neanderthal-matching alleles in Eurasian populations, while a large majority of deep branches in 1000 Genomes West African populations do not match either the Neanderthal or Denisovan samples. This observation was taken as further evidence for deep population structure or archaic hominin admixture in West African populations from an unidentified hominin unrelated to the Neanderthal/Denisovan complex.

### 5 Predictions from inferred demographic models

#### 5.1 Divergence between coexisting populations over time

We considered a few different measures of differentiation between populations in four inferred models, starting with $F_{ST}$. For each of the four inferred demographic models presented here, we computed predicted $F_{ST}$ for pairs of contemporaneous populations over time.

$F_{ST}$ represents the proportion of genetic variance in a multi-population sample that can be attributed to drift between populations in that sample (e.g., Bhatia *et al.* (2013)). $F_{ST}$ depends on population divergence times, gene flow, and population sizes. For present-day sampled populations, predicted $F_{ST}$ can be compared to data (Fig. S8), while past estimates represent model predictions about the differentiation between past populations. We computed $F_{ST}$ as a ratio of expected (or average) variances, using observed pairwise diversity within and between populations as in Peter (2016):

$$F_{ST}(i,j) = \frac{\mathbb{E}[2H_{i,j} - H_i - H_j]}{\mathbb{E}[2H_{i,j} + H_i + H_j]}, \tag{S1}$$

where $H_{ij}$ is the average number of differences between one haploid copy of the genome from population $i$ and one from population $j$, and $H_i$ and $H_j$ are the average number of differences for two haploid copies from the same population.

In all inferred models, $F_{ST}$ between Neanderthals and the ancestors of contemporary humans were much larger than between any pair of contemporary human populations or their ancestors (stems). $F_{ST}$ between human populations over the last $100ka$ remained relatively low, at $\sim 0.1$ or less. In the continuous-migration model, the two stem populations also were predicted to have low $F_{ST}$ within this range, as migration rates were inferred to be high enough to maintain genetic similarity. In both the merger-without-stem-migration and the merger-with-stem-migration, one of the stem populations was inferred to have a short, severe bottleneck, which led to increased $F_{ST}$ between stem populations during that time. In the merger-with-stem-migration, $F_{ST}$ reached a maximum value of 0.3 between stems, intermediate between human-Neanderthal divergence (0.45) and the maximum $F_{ST}$ observed between contemporary human populations ($\lessapprox 0.1$.)

Whereas the high $F_{ST}$ value between Neanderthal and humans reflects both elevated genetic differences between the genomes of humans and Neanderthals and reduced diversity within the Neanderthal population (Fig. S8), the increased $F_{ST}$ between the stems is primarily due to an inferred bottleneck reducing stem diversity, while genetic differentiation between individuals from the two stems is within the range of genetic differentiation between (and even within) human populations. In other words, our best fit model describes two stems with one stem showing reduced diversity, but individuals across stems showing similar differentiation to that between two contemporary human population.

#### 5.2 Shared differentiation between pairs of contemporary and pairs of ancient populations

To evaluate the degree to which divergence among contemporary human populations is inherited from divergence between early stem population, we computed $f_4$ statistics of the form

$$f_4(\text{stem}_1, \text{stem}_2; \text{present}_1, \text{present}_2) = \mathbb{E}[(p_{\text{stem}_1} - p_{\text{stem}_2})(p_{\text{present}_1} - p_{\text{present}_2})] \tag{S2}$$

This statistic measures co-linearity of genetic drift between contemporary populations $v = (p_{\text{present}_1} - p_{\text{present}_2})$ and stem populations $w = (p_{\text{stem}_1} - p_{\text{stem}_2})$ (see Figures S10–S13 and Lipson (2020) for an overview of $f_4$ statistics).

To contrast these statistics to divergence across contemporary populations,

$$f_2(\text{present}_1, \text{present}_2) = \mathbb{E}[(p_{\text{present}_1} - p_{\text{present}_2})^2], \tag{S3}$$

we interpret the $f$-statistics as scalar products between vectors $v$ and $w$, i.e., $f_2(\text{present}_1, \text{present}_2) = v \cdot v$ and $f_4(\text{stem}_1, \text{stem}_2; \text{present}_1, \text{present}_2) = v \cdot w$. We can then write $v = v_\perp + v_\parallel$ as the sum of a component $v_\perp$ perpendicular and a component $v_\parallel$ parallel to $w = (p_{\text{stem}_1} - p_{\text{stem}_2})$. We can then decompose

$f_2(\text{present}_1, \text{present}_2) = v_\perp^2 + v_\parallel^2$, with

$$v_\parallel^2 = \frac{(v \cdot w)^2}{w \cdot w} = \frac{f_4^2(\text{stem}_1, \text{stem}_2; \text{present}_1, \text{present}_2)}{f_2(\text{stem}_1, \text{stem}_2)} \tag{S4}$$

Finally, the proportion of present-day structure $v^2 = f_2(\text{present}_1, \text{present}_2)$ that is accounted for by the component parallel to ancient population structure is

$$\alpha^2 = \frac{v_\parallel^2}{v^2} = \frac{f_4^2(\text{stem}_1, \text{stem}_2; \text{present}_1, \text{present}_2)}{f_2(\text{stem}_1, \text{stem}_2) f_2(\text{present}_1, \text{present}_2)}. \tag{S5}$$

$\alpha^2$ can be interpreted as the proportion of the $f_2$ divergence across contemporary populations that is aligned with the differences in allele frequencies across ancient populations. However, drift occurring after the time of sampling of ancient populations is not correlated to drift occurring before that time. For this reason, we also interpret $\alpha^2$ as the proportion of the $f_2$ divergence across contemporary populations that is *explained by* the differences in allele frequencies across ancient populations. We report this predicted statistic for each of the four inferred demographic models in Figures S10–S13. In our best fit models (continuous migration and merger with stem migration), differentiation between Stem 1 and Stem 2 (before the split of Stem 1 into Stem 1E and Stem 1S in the merger model) contributes a maximum of 4% to differentiation between the Nama and Gumuz, and closer to 1% to differentiation between the Nama and Mende or the Mende and Gumuz.

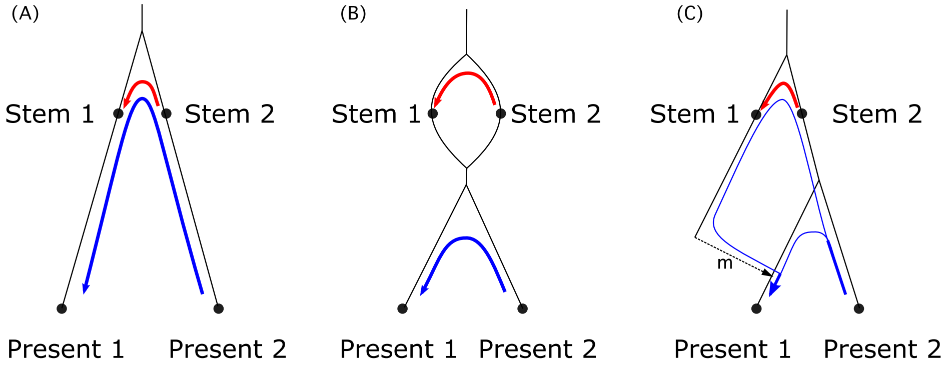

Figure S1: **Present-day structure attributable to ancient population structure.** The statistic $f_4(\text{stem}_1, \text{stem}_2; \text{present}_1, \text{present}_2)$ captures how much of present-day population structure can be attributed to population structure among the stems. The blue line represents genetic drift along ancestral branches between contemporary populations, while the red line represents drift between stem populations. $f_4$ measures the overlap between the blue and red line, which corresponds to shared genetic drift between the two pairs. In scenario A, contemporary populations are descended from distinct stems, and $f_4(\text{stem}_1, \text{stem}_2; \text{present}_1, \text{present}_2)$ equals the drift between stems, $f_2(\text{stem}_1, \text{stem}_2)$. In scenario B, despite deep population structure, there is no shared drift and $f_4 = 0$. That is, present-day population structure is unrelated to structure among stems. In more realistic scenario C, where $\text{present}_1$ receives a proportion $m$ of its ancestry from the dotted arrow, the overlap is only found along the corresponding path, and $f_4$ depends on both ancestry proportion $m$ and drift between stems, $f_4(\text{stem}_1, \text{stem}_2; \text{present}_1, \text{present}_2) = m f_2(\text{stem}_1, \text{stem}_2)$

#### 5.3 Relating model predictions to morphological variation

Here we show how the predictions of our model can be related to morphological variation in the fossil record. Let $\Delta^2$ be the expected squared difference in a morphological measurement between fossil 1 and fossil 2,

$$\Delta^2 = \mathbb{E}\left[\left(\bar{X}_1 - \bar{X}_2\right)^2\right] + \mathbb{E}\left[V_1\right] + \mathbb{E}\left[V_2\right], \tag{S6}$$

where $\bar{X}_1$ and $V_1$ and $\bar{X}_2$ and $V_2$ are measurement means and variances for the populations from which fossil 1 and fossil 2 are sampled.

If we assume that the trait being measured is evolving neutrally, which seems reasonable as an approximation based on studies of morphological (cranial) variation in humans and Neanderthals (Relethford, 1994; Weaver *et al.*, 2008; von Cramon-Taubadel, 2009), then

$$\mathbb{E}\left[\left(\bar{X}_1 - \bar{X}_2\right)^2\right] = 4\sigma_m^2\tau_B - 2\sigma_m^2\tau_{W_1} - 2\sigma_m^2\tau_{W_2}, \tag{S7}$$

$$\mathbb{E}\left[V_1\right] = \frac{1}{h^2}\sigma_m^2\tau_{W_1}, \tag{S8}$$

and

$$\mathbb{E}\left[V_2\right] = \frac{1}{h^2}\sigma_m^2\tau_{W_2}, \tag{S9}$$

where $\sigma_m^2$ is the additive genetic variance introduced by mutation (per zygote per generation), $h^2$ is the (narrow sense) heritability, $\tau_{W_1}$ and $\tau_{W_2}$ are the average coalescence time of pairs of alleles from the same population, and $\tau_B$ is the average coalescence time of pairs of alleles from different populations (Whitlock, 1999; Weaver, 2016). Substituting and simplifying gives

$$\Delta^2 = \sigma_m^2\left(4\tau_B + \frac{1-2h^2}{h^2}\left(\tau_{W_1} + \tau_{W_2}\right)\right), \tag{S10}$$

which relates $\Delta^2$ to between and within population coalescence times.

As a further approximation, if we assume that the heritability of the measurement is close to one half, then

$$\Delta^2 \approx 4\sigma_m^2\tau_B, \tag{S11}$$

which shows that under these assumptions $\Delta^2$ is proportional to $\tau_B$.

Nucleotide diversity, $\pi$, for pairs of individuals sampled from different populations is also proportional to $\tau_B$. Therefore, ratios of $\Delta^2$ can be predicted from ratios of $\pi$ for the same comparisons. For example, a ratio for which $\pi$ for Stem 1 and Stem 2 is the numerator and $\pi$ for a pair of contemporary human populations is the denominator can be used to predict $\Delta^2$ for two fossils sampled from each of the two stems based on $\Delta^2$ for contemporary humans. In this way, the predictions of our model can be related to morphological variation in the fossil record.

### 6 Validations using simulations from inferred demographic models

Here we focus on four parameterizations of early human history: (1) a "single origin" with recent expansion, (2) primary and second stem populations with "continuous migration", (3) two stem populations that do not exchange migrants before merging to form regional African populations, referred to as a "merger without stem migration", and (4) two stem populations that are allowed to exchange migrants before merging to form regional African populations, referred to as a "merger with stem migration". Given the best fit parameters for each model (Tables S2–S6), we used `msprime` version 1.0.4 (Kelleher *et al.*, 2016; Baumdicker *et al.*, 2022) to simulate sampled individual genomes to match the data used in inference (see simulation details below). We used these simulations to compare our model predictions to predictions based on the conditional SFS (cSFS) and on relate curves (see below).

These comparisons serve as independent validations of our inferred demographic models by assessing how well those models fit aspects of the data that have been used in other studies as evidence of population structure or archaic hominin admixture from an unidentified hominin species within Africa.

#### 6.1 Simulation details

The four inferred demographic models (single origin, continuous migration, merger without stem migration, and merger with stem migration) were converted to `demes` format (Gower, Ragsdale, *et al.*, in prep) and used as the input demography for `msprime` simulations. We sampled the same number of individuals from each population as used in the inference (Nama: 44, Mende: 85, Gumuz: 23, combined Oromo/Amhara: 46, British: 91, and Vindija Neanderthal: 1). We assumed a generation time of 29 years, a per-base mutation rate of $1.25 \times 10^{-8}$, a per-base recombination rate of $1 \times 10^{-8}$ (corresponding to 1cM/Mb), and we simulated 100 replicates with sequence lengths of 50Mb per replicate. We computed the conditional site frequency spectrum (cSFS) conditioned on observing the derived variant in the Neanderthal sample, and reconstructed gene genealogies using `Relate` to obtain coalescence rate trajectories and distributions of deep branches passing through the Neanderthal branch. Each analysis followed the same procedure as described above in the analysis of the real data.

#### 6.2 cSFS prediction under inferred models

We compared the observed cSFS to model predictions for the Nama, Mende, and Gumuz populations. The cSFS from data is sensitive to ancestral allele misidentification, but our simulations assume perfect knowledge of the ancestral allele. To investigate the role of misidentification, we compared predicted and observed cSFS for transitions and transversions separately as well as jointly.

For each comparison, we fit a misidentification parameter $p_{misid}$ as well as a scaled mutation rate $\theta$. The misidentification parameter is the fraction $p_{misid}$ of loci that have the ancestral state misidentified. Accounting for $p_{misid}$ mixes the predicted joint population SFS with a small amount of the ancestral-allele-reversed SFS (i.e. for a two-population joint SFS, $SFS_{misid}(i,j) = (1-p_{misid}) * SFS(i,j) + p_{misid} * SFS(n-i, m-j)$), where $n$ and $m$ are the sample sizes from the two populations. The cSFS for the first population (conditioning on the second) is found by projecting the sample size to $(n, 1)$ and taking $cSFS = SFS(:, 1)$. $\theta$ and $p_{misid}$ were chosen to maximize a likelihood assuming that each observed entry in the cSFS is drawn from a Poisson distribution (Sawyer and Hartl, 1992).

### 7 Supplementary results

#### 7.1 Alternative statistics for four models

##### 7.1.1 Conditional SFS

The single-origin model fit the cSFS relatively poorly across the three populations compared to the models that allow structure between stem populations (Fig. S14). The model that best fit the cSFS was the merger-with-stem-migration model, which also provided the best fit to pairwise diversity and LD statistics in the initial optimization. For the Mende and Gumuz, the merger-with-stem-migration model fits the data very well, while it underestimates rare variants in the Nama cSFS (Fig. S17). Models learned from the two-locus statistics provided a better qualitative description of the cSFS than models previously fit directly to the cSFS (see, e.g., figure 1 in Durvasula and Sankararaman (2020)). We found that the inferred ancestral misidentification rates were concordant across populations. The inferred rates of ancestral misidentification were about twice as high for transitions relative to transversions as expected, qualitatively, from the higher transition mutation rate (Hernandez *et al.*, 2007).

##### 7.1.2 Relate curves from inferred models

We used `Relate` (Speidel *et al.*, 2019) to reconstruct gene genealogies from the simulations under our four inferred demographic models. From each of these, following the same approach as for the analysis of the actual data, we computed coalescence rates within and between sampled populations. From the coalescence rates, we estimated the inverse instantaneous coalescence rate (IICR, often interpreted as an estimator of $N_e$) for each population, as well as relative cross coalescence rates between pairs of populations.

The IICR is only a reliable estimator of $N_e$ in the absence of population structure. However, since IIRC are readily estimated and visually interesting, it is commonly attempted even when the assumption is not met. Even if the assumption is not met, the inferred IIRC can be interpreted as a summary statistic for which model predictions can be compared to data. It can therefore be informative about aspects of the data that our models fail to predict. We also note that the coalescence rates inferred from data used a larger set of populations from the combined 1000 Genomes/AGRP dataset, while our model simulations only included a subset of 5 populations. This may lead to differences in resolution of the inferences.

The data shows the characteristic "hump" of increased IICR ($N_e$) 100-300k for all populations, followed by a decrease in size among all populations that is sharpest in the Eurasian populations and also pronounced in the East African populations that have high proportions of Eurasian ancestry (Amhara and Oromo) (Fig. S18). The model-inferred $N_e$ curves also show this general trend for the more recent reduction in $N_e$ (Fig. S20). In the the more distant past, $N_e$ fluctuates in a manner that is broadly similar across models with observed "humps" of increased $N_e$. Despite the large differences in parameterizations and interpretations of the four inferred demographic models, each of them show reconstructed $N_e$ trajectories that are qualitatively similar, highlighting the difficulty in learning complex demography in the deep past from reconstructed coalescence rate trajectories.

We also compared the relative cross-coalescence rates (RCCR) between pairs of populations in the four inferred demographic models (Figures S23–S26) to those in the data (Fig. S22). In the recent past (<10ka), each of the models show increased RCCR compared to the data. However, in the medium to more distant past, the RCCR provide a good match to the data in each of the four models, including both their timing and ordering of increased cross coalescences. The failure to match both the IIRC and RCCR in the recent past in all models suggests that additional parameters would be needed to account for recent increases in population sizes. As is the case with other methods that reconstruct size history trajectories from inferred coalescence rates (e.g. PSMC, MSMC), `Relate` has difficulty inferring coalescences that occur in the recent past.

###### 7.1.3 Distribution of deep branch affinities to Neanderthal sequence

Following Speidel *et al.* (2019), we also characterized Neanderthal-matching deep branches (see Section 4) in our simulated data under the four inferred demographic models. Comparing deep-branch distributions from the data (Fig. S28) to the models (Fig. S29), we find consistent trends between each of the four models and the data in the ordering of proportions of lineages from each population that are labeled as Neanderthal-matching deep branches. This proportion is slightly over-estimated in each of the African populations compared to the data, but is closest in the merger-with-stem-migration model (which also fit the LD statistics the best of the four highlighted models). Given the qualitative discrepancies between the data IICR and the model IICR, the differences in Neanderthal branch affinities is unsurprising.

##### 7.2 Mutation versus recombination rates

Prior studies primarily relied on the mutation clock in order to date events such as population divergence, and uncertainty about the mutation rate translates in uncertainty in timing estimates (e.g., Moorjani *et al.* (2016a)). Recent whole genome sequencing studies from non-pathological pedigrees have typically estimated mutation rates to be $1.2-1.3\times10^{-8}$ per site per generation (Sasani *et al.*, 2019; Tian *et al.*, 2019). A rate of $1.3\times10^{-8}$ also approximates the rate predicted by a mutational model from gnomAD (Karczewski *et al.*, 2020) in the ~650Mb of intergenic regions selected for demographic inference in this study. `moments-LD` differs from other approaches by relying on a recombination clock, which has been argued to be more robust to estimation than a mutational clock (Moorjani *et al.*, 2016b).

From our recombination-rate-calibrated demographic models, we can estimate the mutation rates needed to fit patterns of pairwise diversity. With our best-fit model (merger-with-stem-migration, Table S6), a mutation rate of $1.33\times10^{-8}$ recovers the observed diversity within and between populations, closely matching the average mutation rate estimated from the gnomAD mutation model. The other inferred demographic models (Tables S2–S5) would require mutation rates of $1.75-1.90\times10^{-8}$ to match pairwise diversity.

##### 7.3 Re-inference of IM models from simulated data

Recent studies have focused on characterizing the timing of the earliest divergences among human populations, and such analyses typically fit simple isolation-with-migration (IM) or "clean split" models to observed allele frequencies or inferred coalescence times among and between populations (see Weaver *et al.* (2008); Bergström *et al.* (2021) for recent reviews). Simple IM models typically fit a constant $N_e$ within the two diverged population, the split time, and a symmetric continuous migration rate (which could be fixed to zero, equivalent to the "clean split" model). We explored the effect of fitting such simplified models of history to data from more complex models by simulating the site frequency spectrum (SFS) under each of our four models for pairs of populations and reinferring an IM and clean split model for each pair. The simulations sampled 10 diploid individuals from each population, and we used `msprime` (Baumdicker *et al.*, 2022) to simulated 500 Mb of total sequence length (the sum over 500 replicates each 1 Mb in length) with constant recombination rate $r = 10^{-8}$ and mutation rate $u = 2 \times 10^{-8}$.

As expected, the clean split models consistently provide divergence times $T$ that are more recent than the IM models. Because $T$ and subsequent migration can be inversely confounded in SFS inference, fixing migration rates to zero results in more recent inferred $T$ (Fig. S30). Even though each of our four models inferred divergences of all human populations ∼120ka, ancient structure and size changes in the simulations biases re-inferences and gives inflated estimates of $T$. Such an effect has been shown to occur when the ancestral population size increases prior to the population split (Momigliano *et al.*, 2021), which is the case in our inferred single-origin model. Re-inferred $T$ between the Nama and other populations were >200ka under the IM model, and ∼50–130ka under the clean split model.

### References

1000 Genomes Project Consortium, A. Auton, L. D. Brooks, R. M. Durbin, E. P. Garrison, *et al.*, 2015 A global reference for human genetic variation. Nature **526**: 68–74.

Alexander, D. H., and K. Lange, 2011 Enhancements to the ADMIXTURE algorithm for individual ancestry estimation. BMC Bioinformatics **12**: 246.

Alexander, D. H., J. Novembre, and K. Lange, 2009 Fast model-based estimation of ancestry in unrelated individuals. Genome Res. **19**: 1655–1664.

Baumdicker, F., G. Bisschop, D. Goldstein, G. Gower, A. P. Ragsdale, *et al.*, 2022 Efficient ancestry and mutation simulation with msprime 1.0. Genetics **220**.

Bergström, A., C. Stringer, M. Hajdinjak, E. M. L. Scerri, and P. Skoglund, 2021 Origins of modern human ancestry. Nature **590**: 229–237.

Bhatia, G., N. Patterson, S. Sankararaman, and A. L. Price, 2013 Estimating and interpreting FST: the impact of rare variants. Genome Res. **23**: 1514–1521.

Chen, H., R. E. Green, S. Pääbo, and M. Slatkin, 2007 The joint allele-frequency spectrum in closely related species. Genetics **177**: 387–398.

Coffman, A. J., P. H. Hsieh, S. Gravel, and R. N. Gutenkunst, 2016 Computationally Efficient Composite Likelihood Statistics for Demographic Inference. Mol. Biol. Evol. **33**: 591–593.

Danecek, P., J. K. Bonfield, J. Liddle, J. Marshall, V. Ohan, *et al.*, 2021 Twelve years of SAMtools and BCFtools. Gigascience **10**.

Delaneau, O., J.-F. Zagury, and J. Marchini, 2013 Improved whole-chromosome phasing for disease and population genetic studies. Nat. Methods **10**: 5–6.

DePristo, M. A., E. Banks, R. Poplin, K. V. Garimella, J. R. Maguire, *et al.*, 2011 A framework for variation discovery and genotyping using next-generation DNA sequencing data. Nat. Genet. **43**: 491–498.

Durvasula, A., and S. Sankararaman, 2020 Recovering signals of ghost archaic introgression in African populations. Sci Adv **6**: eaax5097.

Gopalan, S., R. E. W. Berl, J. W. Myrick, Z. H. Garfield, A. W. Reynolds, *et al.*, 2022 Hunter-gatherer genomes reveal diverse demographic trajectories during the rise of farming in Eastern Africa. Curr. Biol. .

Gurdasani, D., T. Carstensen, F. Tekola-Ayele, L. Pagani, I. Tachmazidou, *et al.*, 2015 The African Genome Variation Project shapes medical genetics in Africa. Nature **517**: 327–332.

Harris, K., and R. Nielsen, 2014 Error-prone polymerase activity causes multinucleotide mutations in humans. Genome Res. **24**: 1445–1454.

Hernandez, R. D., S. H. Williamson, and C. D. Bustamante, 2007 Context dependence, ancestral misidentification, and spurious signatures of natural selection. Mol. Biol. Evol. **24**: 1792–1800.

Hill, W. G., and A. Robertson, 1968 Linkage disequilibrium in finite populations. Theor. Appl. Genet. **38**: 226–231.

International HapMap Consortium, K. A. Frazer, D. G. Ballinger, D. R. Cox, D. A. Hinds, *et al.*, 2007 A second generation human haplotype map of over 3.1 million SNPs. Nature **449**: 851–861.

Karczewski, K. J., L. C. Francioli, G. Tiao, B. B. Cummings, J. Alföldi, *et al.*, 2020 The mutational constraint spectrum quantified from variation in 141,456 humans. Nature **581**: 434–443.

Kelleher, J., A. M. Etheridge, and G. McVean, 2016 Efficient Coalescent Simulation and Genealogical Analysis for Large Sample Sizes. PLoS Comput. Biol. **12**: e1004842.

Lipson, M., 2020 Applying f4 -statistics and admixture graphs: Theory and examples. Mol. Ecol. Resour. **20**: 1658–1667.

Miles, A., M. R., P. Ralph, N. Harding, R. Pisupati, *et al.*, 2021 cggh/scikit-allel: v1.3.3.

Momigliano, P., A.-B. Florin, and J. Merilä, 2021 Biases in Demographic Modeling Affect Our Understanding of Recent Divergence. Mol. Biol. Evol. **38**: 2967–2985.

Moorjani, P., Z. Gao, and M. Przeworski, 2016a Human Germline Mutation and the Erratic Evolutionary Clock. PLoS Biol. **14**: e2000744.

Moorjani, P., S. Sankararaman, Q. Fu, M. Przeworski, N. Patterson, *et al.*, 2016b A genetic method for dating ancient genomes provides a direct estimate of human generation interval in the last 45,000 years. Proc. Natl. Acad. Sci. U. S. A. **113**: 5652–5657.

Ohta, T., and M. Kimura, 1971 Linkage disequilibrium between two segregating nucleotide sites under the steady flux of mutations in a finite population. Genetics **68**: 571–580.

Pagani, L., S. Schiffels, D. Gurdasani, P. Danecek, A. Scally, *et al.*, 2015 Tracing the route of modern humans out of Africa by using 225 human genome sequences from Ethiopians and Egyptians. Am. J. Hum. Genet. **96**: 986–991.

Pedregosa, F., G. Varoquaux, A. Gramfort, V. Michel, B. Thirion, *et al.*, 2011 Scikit-learn: Machine learning in Python. the Journal of machine Learning research **12**: 2825–2830.

Peter, B. M., 2016 Admixture, Population Structure, and F-Statistics. Genetics **202**: 1485–1501.

Plagnol, V., and J. D. Wall, 2006 Possible ancestral structure in human populations. PLoS Genet. **2**: e105.

Prüfer, K., C. de Filippo, S. Grote, F. Mafessoni, P. Korlević, *et al.*, 2017 A high-coverage Neandertal genome from Vindija Cave in Croatia. Science **358**: 655–658.

Ragsdale, A. P., and S. Gravel, 2019 Models of archaic admixture and recent history from two-locus statistics. PLoS Genet. **15**: e1008204.

Ragsdale, A. P., and S. Gravel, 2020 Unbiased Estimation of Linkage Disequilibrium from Unphased Data. Mol. Biol. Evol. **37**: 923–932.

Ragsdale, A. P., C. Moreau, and S. Gravel, 2018 Genomic inference using diffusion models and the allele frequency spectrum. Curr. Opin. Genet. Dev. **53**: 140–147.

Relethford, J. H., 1994 Craniometric variation among modern human populations. Am. J. Phys. Anthropol. **95**: 53–62.

Sasani, T. A., B. S. Pedersen, Z. Gao, L. Baird, M. Przeworski, *et al.*, 2019 Large, three-generation human families reveal post-zygotic mosaicism and variability in germline mutation accumulation. Elife **8**.

Sawyer, S. A., and D. L. Hartl, 1992 Population genetics of polymorphism and divergence. Genetics **132**: 1161–1176.

Speidel, L., M. Forest, S. Shi, and S. R. Myers, 2019 A method for genome-wide genealogy estimation for thousands of samples. Nat. Genet. **51**: 1321–1329.

Tian, X., B. L. Browning, and S. R. Browning, 2019 Estimating the Genome-wide Mutation Rate with Three-Way Identity by Descent. Am. J. Hum. Genet. **105**: 883–893.

Uren, C., M. Kim, A. R. Martin, D. Bobo, C. R. Gignoux, *et al.*, 2016 Fine-Scale Human Population Structure in Southern Africa Reflects Ecogeographic Boundaries. Genetics **204**: 303–314.

Virtanen, P., R. Gommers, T. E. Oliphant, M. Haberland, T. Reddy, *et al.*, 2020 SciPy 1.0: fundamental algorithms for scientific computing in Python. Nat. Methods **17**: 261–272.

von Cramon-Taubadel, N., 2009 Congruence of individual cranial bone morphology and neutral molecular affinity patterns in modern humans. Am. J. Phys. Anthropol. **140**: 205–215.

Weaver, T. D., 2016 Estimators for QST and coalescence times. Ecol. Evol. **6**: 7783–7786.

Weaver, T. D., C. C. Roseman, and C. B. Stringer, 2008 Close correspondence between quantitative- and molecular-genetic divergence times for Neandertals and modern humans. Proc. Natl. Acad. Sci. U. S. A. **105**: 4645–4649.

Whitlock, M. C., 1999 Neutral additive genetic variance in a metapopulation. Genet. Res. **74**: 215–221.

Yang, M. A., A.-S. Malaspinas, E. Y. Durand, and M. Slatkin, 2012 Ancient structure in Africa unlikely to explain Neanderthal and non-African genetic similarity. Mol. Biol. Evol. **29**: 2987–2995.

#### Supporting tables and figures

Table S1: **Fixed parameters in inferred demographic models.** Many of the fixed parameters are specific to the Neanderthal branch, are constrained by known history, or were consistently fit across multiple model parameterizations. We assumed a generation time of 29 years throughout. See Section 3.1 for details.

| Parameter | Description | Value |
|---|---|---|
| $T_{MH-Neand}$ | The time of the split between branches leading to Neanderthals and humans | 550ka |
| $T_{NI-Vin}$ | The time of the split between branches leading to the Vindija sample and the Neanderthal population that mixed with expanding humans | 80ka |
| $T_{Vin}$ | The sample time of the Vindija Neanderthal individual | 50ka |
| $T_{NI\to EUR}$ | The timing of admixture from Neanderthals to humans | 45ka |
| $f_{NI\to EUR}$ | The proportion of admixture from Neanderthals to humans | 0.015 |
| $T_{E-W}$ | The split time of East and West African populations | 60ka |
| $T_{GBR}$ | The split time of Eurasian and East African populations | 50ka |
| $T_{GBR\to EA}$ | The time of the back-to-Africa admixture event, giving rise to the Amhara/Oromo branch | 12ka |
| $T_{expan}$ | The expansion time of the Mende and contraction time of the Gumuz | 5ka |
| $T_{EA\to Nama}$ | The admixture time of East African pastoralists to the Nama | 67 gen. |
| $T_{GBR\to Nama}$ | The admixture time of Europeans to the Nama | 10 gen. |
| $T_{bottle}$ | The bottleneck time of the Nama population | 9 gen. |

Table S2: **Best-fit parameters from the single-origin model.** This model includes a single stem population that splits into contemporary human groups. Fixed parameters are specifed in Table S1. Inferred values are scaled to physical units assuming a generation time of 29 years. This model gave a log-likelihood of -189,434.

| Parameter | Description | Value | Std. err. |
|---|---|---|---|
| $N_e$ | Ancestral effective population size | 10198 | 403 |
| $N_{MH}$ | Size of human lineage between Neanderthal and Nama splits | 21111 | 529 |
| $N_{Nama_0}$ | Initial Nama size | 10224 | 370 |
| $N_{Nama_F}$ | Final Nama size | 222 | 9 |
| $N_{MSL_0}$ | Initial Mende size | 17211 | 769 |
| $N_{MSL_F}$ | Final Mende size | 16822 | 606 |
| $N_{EA}$ | Size of East African branch | 7139 | 273 |
| $N_{Gumuz_F}$ | Final Gumuz size | 3831 | 131 |
| $N_{EP}$ | East African agriculturist size | 13033 | 491 |
| $N_{GBR_0}$ | Initial British size | 846 | 33 |
| $N_{GBR_F}$ | Final British size | 12121 | 507 |
| $N_{Neand}$ | Neanderthal size | 1867 | 105 |
| $T_{Nama}$ | Nama split time (years) | 110400 | 2525 |
| $m_{Nama-MSL}$ | Nama–Mende symmetric migration rate | $2.82 \times 10^{-5}$ | $0.158 \times 10^{-5}$ |
| $m_{Nama-EA}$ | Nama–East Africa symmetric migration rate | $4.94 \times 10^{-5}$ | $0.197 \times 10^{-5}$ |
| $m_{MSL-EA}$ | Mende–East Africa migration rate | $18.76 \times 10^{-5}$ | $0.764 \times 10^{-5}$ |
| $m_{EA-GBR}$ | East Africa–Europe migration rate | $4.42 \times 10^{-5}$ | $0.239 \times 10^{-5}$ |
| $m_{EA-EA}$ | Intra-East Africa migration rate | $41.28 \times 10^{-5}$ | $1.33 \times 10^{-5}$ |
| $f_{GBR \to EP}$ | Ancestry proportion of East African agriculturalists from GBR 12ka ($1 - f$ from Gumuz) | 0.658 | 0.0039 |
| $f_{EP \to Nama}$ | Ancestry proportion from EA pastoralists to Nama 2ka | 0.279 | 0.0039 |
| $f_{GBR \to Nama}$ | Ancestry proportion from Europeans to Nama 10 generations ago | 0.150 | 0.0019 |

Table S3: **Best-fit parameters from the Continuous-Migration model without stem migration.** Inferred values are scaled to physical units assuming a generation time of 29 years. This model gave a log-likelihood of -126,644. This model allows for continuous migration between Stem 2 and contemporary populations, but disallows migration between Stem 2 and Stem 1, which later splits into branches leading to South, East, and West African populations.

| Parameter | Description | Value | Std. err. |
|---|---|---|---|
| $N_e$ | Ancestral effective population size | 10543 | 460 |
| $N_{stem1}$ | Size of stem 1 lineage between Neanderthal and Nama splits | 15297 | 1227 |
| $N_{stem2}$ | Size of stem 2 lineage | 9156 | 573 |
| $N_{Nama_0}$ | Initial Nama size | 11329 | 423 |
| $N_{Nama_F}$ | Final Nama size | 219 | 11 |
| $N_{MSL_0}$ | Initial Mende size | 11187 | 644 |
| $N_{MSL_F}$ | Final Mende size | 24056 | 945 |
| $N_{EA}$ | Size of East African branch | 7185 | 293 |
| $N_{Gumuz_F}$ | Final Gumuz size | 3746 | 184 |
| $N_{EP}$ | East African agriculturist size | 13187 | 645 |
| $N_{GBR_0}$ | Initial British size | 937 | 41 |
| $N_{GBR_F}$ | Final British size | 11478 | 420 |
| $N_{Neand}$ | Neanderthal size | 2285 | 113 |
| $T_{stems}$ | Stem split time (years) | 459720 | 21704 |
| $T_{Nama}$ | Nama split time (years) | 139725 | 3250 |
| $m_{Nama-MSL}$ | Nama–Mende symmetric migration rate | $1.93 \times 10^{-5}$ | $0.360 \times 10^{-5}$ |
| $m_{Nama-EA}$ | Nama–East Africa symmetric migration rate | $4.43 \times 10^{-5}$ | $0.309 \times 10^{-5}$ |
| $m_{MSL-EA}$ | Mende–East Africa migration rate | $20.8 \times 10^{-5}$ | $0.97 \times 10^{-5}$ |
| $m_{EA-GBR}$ | East Africa–Europe migration rate | $4.46 \times 10^{-5}$ | $0.171 \times 10^{-5}$ |
| $m_{EA-EA}$ | Intra-East Africa migration rate | $35.4 \times 10^{-5}$ | $1.85 \times 10^{-5}$ |
| $f_{GBR \to EP}$ | Ancestry proportion of East African agriculturalists from GBR 12ka ($1 - f$ from Gumuz) | 0.649 | 0.0038 |
| $f_{EP \to Nama}$ | Ancestry proportion from EA pastoralists to Nama 2ka | 0.268 | 0.0048 |
| $f_{GBR \to Nama}$ | Ancestry proportion from Europeans to Nama 10 generations ago | 0.153 | 0.0022 |
| $m_{stems}$ | Stem 1–stem 2 migration rate | 0 | fixed |
| $m_{stem2-Nama}$ | Stem 2–Nama migration rate | $4.33 \times 10^{-5}$ | $0.891 \times 10^{-5}$ |
| $m_{stem2-MSL}$ | Stem 2–Mende migration rate | $11.3 \times 10^{-5}$ | $1.83 \times 10^{-5}$ |
| $m_{stem2-EA}$ | Stem 2–East Africa migration rate | $2.77 \times 10^{-5}$ | $0.699 \times 10^{-5}$ |

Table S4: **Best-fit parameters from the Continuous-Migration model.** Inferred values are scaled to physical units assuming a generation time of 29 years. This model gave a log-likelihood of -115,500.

| Parameter | Description | Value | Std. err. |
|---|---|---|---|
| $N_e$ | Ancestral effective population size | 7270 | 1777 |
| $N_{stem1}$ | Size of stem 1 lineage between Neanderthal and Nama splits | 8256 | 1612 |
| $N_{stem2}$ | Size of stem 2 lineage | 13547 | 2488 |
| $N_{Nama_0}$ | Initial Nama size | 11939 | 2989 |
| $N_{Nama_F}$ | Final Nama size | 221 | 54 |
| $N_{MSL_0}$ | Initial Mende size | 9738 | 2479 |
| $N_{MSL_F}$ | Final Mende size | 28150 | 6628 |
| $N_{EA}$ | Size of East African branch | 7489 | 1841 |
| $N_{Gumuz_F}$ | Final Gumuz size | 3728 | 915 |
| $N_{EP}$ | East African agriculturist size | 13072 | 3246 |
| $N_{GBR_0}$ | Initial British size | 959 | 231 |
| $N_{GBR_F}$ | Final British size | 11822 | 2889 |
| $N_{Neand}$ | Neanderthal size | 2670 | 591 |
| $T_{stems}$ | Stem split time (years) | 1163072 | 390803 |
| $T_{Nama}$ | Nama split time (years) | 134745 | 17775 |
| $m_{Nama-MSL}$ | Nama–Mende symmetric migration rate | $0.98 \times 10^{-5}$ | $0.366 \times 10^{-5}$ |
| $m_{Nama-EA}$ | Nama–East Africa symmetric migration rate | $4.08 \times 10^{-5}$ | $1.02 \times 10^{-5}$ |
| $m_{MSL-EA}$ | Mende–East Africa migration rate | $21.4 \times 10^{-5}$ | $5.32 \times 10^{-5}$ |
| $m_{EA-GBR}$ | East Africa–Europe migration rate | $4.17 \times 10^{-5}$ | $1.02 \times 10^{-5}$ |
| $m_{EA-EA}$ | Intra-East Africa migration rate | $33.6 \times 10^{-5}$ | $8.35 \times 10^{-5}$ |
| $f_{GBR \to EP}$ | Ancestry proportion of East African agriculturalists from GBR 12ka ($1 - f$ from Gumuz) | 0.642 | 0.0037 |
| $f_{EP \to Nama}$ | Ancestry proportion from EA pastoralists to Nama 2ka | 0.255 | 0.0043 |
| $f_{GBR \to Nama}$ | Ancestry proportion from Europeans to Nama 10 generations ago | 0.156 | 0.0021 |
| $m_{stems}$ | Stem 1–stem 2 migration rate | $6.43 \times 10^{-5}$ | $1.05 \times 10^{-5}$ |
| $m_{stem2-Nama}$ | Stem 2–Nama migration rate | $5.82 \times 10^{-5}$ | $1.60 \times 10^{-5}$ |
| $m_{stem2-MSL}$ | Stem 2–Mende migration rate | $16.4 \times 10^{-5}$ | $4.19 \times 10^{-5}$ |
| $m_{stem2-EA}$ | Stem 2–East Africa migration rate | $3.10 \times 10^{-5}$ | $0.901 \times 10^{-5}$ |

Table S5: **Best-fit parameters from the Merger-Without-Stem-Migration model.** Inferred values are scaled to physical units assuming a generation time of 29 years. This model gave a log-likelihood of -107,652.

| Parameter | Description | Value | Std. err. |
|---|---|---|---|
| $N_e$ | Ancestral effective population size | 11258 | 326 |
| $N_{stem1}$ | Size of stem 1 lineage between stem 1–stem 2 split and stem 1E–stem 1S split | 113 | 76 |
| $N_{stem2}$ | Size of stem 2 lineage | 23984 | 1149 |
| $N_{Nama_0}$ | Initial Nama and stem 1S size | 13134 | 384 |
| $N_{Nama_F}$ | Final Nama size | 225 | 7.3 |
| $N_{MSL_0}$ | Initial Mende size | 11856 | 322 |
| $N_{MSL_F}$ | Final Mende size | 25558 | 987 |
| $N_{EA}$ | Size of East African and stem 1E branch | 9136 | 246 |
| $N_{Gumuz_F}$ | Final Gumuz size | 3385 | 102 |
| $N_{EP}$ | East African agriculturist size | 13650 | 408 |
| $N_{GBR_0}$ | Initial British size | 931 | 29 |
| $N_{GBR_F}$ | Final British size | 12064 | 334 |
| $N_{Neand}$ | Neanderthal size | 1935 | 91 |
| $T_{stems}$ | Stem split time (years) | 420881 | 27380 |
| $T_{stem1}$ | Stem 1 split time into stem 1E and stem 1S (years) | 367434 | 19952 |
| $m_{Nama-MSL}$ | Nama–Mende symmetric migration rate | $0.361 \times 10^{-5}$ | $0.113 \times 10^{-5}$ |
| $m_{Nama-EA}$ | Nama–East Africa symmetric migration rate | $4.00 \times 10^{-5}$ | $0.130 \times 10^{-5}$ |
| $m_{MSL-EA}$ | Mende–East Africa migration rate | $19.5 \times 10^{-5}$ | $0.548 \times 10^{-5}$ |
| $m_{EA-GBR}$ | East Africa–Europe migration rate | $3.77 \times 10^{-5}$ | $0.152 \times 10^{-5}$ |
| $m_{EA-EA}$ | Intra-East Africa migration rate | $37.1 \times 10^{-5}$ | $1.26 \times 10^{-5}$ |
| $f_{GBR \to EP}$ | Ancestry proportion of East African agriculturalists from GBR 12ka ($1 - f$ from Gumuz) | 0.647 | 0.0037 |
| $f_{EP \to Nama}$ | Ancestry proportion from EA pastoralists to Nama 2ka | 0.257 | 0.0042 |
| $f_{GBR \to Nama}$ | Ancestry proportion from Europeans to Nama 10 generations ago | 0.156 | 0.0021 |
| $m_{stems}$ | Stem 1–stem 2 migration rate | 0 | fixed |
| $T_{Nama}$ | Time of Nama merger event | 117392 | 8253 |
| $f_{stem2 \to Nama}$ | Proportion of stem 2 ancsestry making up initial Nama lineage ($1 - f$ from stem 1S) | 0.707 | 0.0086 |
| $T_{EA}$ | Time of East Africa merger event | 94892 | 3648 |
| $f_{stem2 \to EA}$ | Proportion of stem 2 ancestry making up initial East Africa lineage ($1 - f$ from stem 1E) | 0.481 | 0.0074 |
| $T_{MSL}$ | Time of secondary admixture from stem 2 to Mende | 23922 | 570 |
| $f_{stem2 \to MSL}$ | Proportion of ancestry from secondary stem 2 admixture to Mende | 0.168 | 0.0036 |

Table S6: **Best-fit parameters from the Merger-With-Stem-Migration model.** Inferred values are scaled to physical units assuming a generation time of 29 years. This model gave a log-likelihood of -102,633.

| Parameter | Description | Value | Std. err. |
|---|---|---|---|
| $N_e$ | Ancestral effective population size | 11479 | 1369 |
| $N_{stem1}$ | Size of stem 1 lineage between Neanderthal split and stem 1E–stem 1S split | 117 | 838 |
| $N_{stem2}$ | Size of stem 2 lineage | 24393 | 6668 |
| $N_{Nama_0}$ | Initial Nama size | 13211 | 1514 |
| $N_{Nama_F}$ | Final Nama size | 223 | 31 |
| $N_{MSL_0}$ | Initial Mende size | 11444 | 1165 |
| $N_{MSL_F}$ | Final Mende size | 27417 | 4332 |
| $N_{EA}$ | Size of East African Branch | 9077 | 1628 |
| $N_{Gumuz_F}$ | Final Gumuz size | 3402 | 337 |
| $N_{EP}$ | East African agriculturist size | 13506 | 1684 |
| $N_{GBR_0}$ | Initial British size | 953 | 122 |
| $N_{GBR_F}$ | Final British size | 12406 | 1678 |
| $N_{Neand}$ | Neanderthal size | 2416 | 235 |
| $T_{stems}$ | Stem split time (years) | 1442022 | 426449 |
| $T_{stem1}$ | Stem 1S–stem 1E split time (years) | 479401 | 166339 |
| $m_{Nama-MSL}$ | Nama–Mende symmetric migration rate | $0.712 \times 10^{-5}$ | $0.401 \times 10^{-5}$ |
| $m_{Nama-EA}$ | Nama–East Africa symmetric migration rate | $4.35 \times 10^{-5}$ | $0.912 \times 10^{-5}$ |
| $m_{MSL-EA}$ | Mende–East Africa migration rate | $19.8 \times 10^{-5}$ | $2.57 \times 10^{-5}$ |
| $m_{EA-GBR}$ | East Africa–Europe migration rate | $3.87 \times 10^{-5}$ | $0.550 \times 10^{-5}$ |
| $m_{EA-EA}$ | Intra-East Africa migration rate | $35.9 \times 10^{-5}$ | $5.36 \times 10^{-5}$ |
| $f_{GBR \rightarrow EP}$ | Ancestry proportion of East African agriculturalists from GBR 12ka ($1 - f$ from Gumuz) | 0.640 | 0.0075 |
| $f_{EP \rightarrow Nama}$ | Ancestry proportion from EA pastoralists to Nama 2ka | 0.257 | 0.0049 |
| $f_{GBR \rightarrow Nama}$ | Ancestry proportion from Europeans to Nama 10 generations ago | 0.157 | 0.0031 |
| $m_{stems}$ | Stem 1–stem 2 migration rate | $11.6 \times 10^{-5}$ | $8.74 \times 10^{-5}$ |
| $T_{Nama}$ | Time of Nama merger event | 118547 | 28170 |
| $f_{stem2 \rightarrow Nama}$ | Proportion of stem 2 ancsestry making up initial Nama lineage ($1 - f$ from stem 1S) | 0.714 | 0.067 |
| $T_{EA}$ | Time of East Africa merger event | 98083 | 8865 |
| $f_{stem2 \rightarrow EA}$ | Proportion of stem 2 ancestry making up initial East Africa lineage ($1 - f$ from stem 1E) | 0.495 | 0.059 |
| $T_{MSL}$ | Time of secondary admixture from stem 2 to Mende | 25119 | 641 |
| $f_{stem2 \rightarrow MSL}$ | Proportion of ancestry from secondary stem 2 admixture to Mende | 0.181 | 0.0085 |

Table S7: **Probability that a lineage sampled 500ya is found in a given branch.** Using our best-fit model (merger with stem migration), we calculated the probability that a lineage sampled in a given population (Nama, Mende, or Gumuz) is found in each branch at 10ka, 55ka, and 500ka. Lineages are sampled 500 years ago to avoid the effects of very recent admixture, such as recent European admixture in the Nama. Rows may not sum to one, as not every branch is listed here (such as Neanderthal).

| | Nama branch | Mende branch | Gumuz branch | Oromo/Amhara branch | British branch | Stem 1 | Stem 2 |
|---|---|---|---|---|---|---|---|
| Found 10ka | | | | | | | |
| Nama | 0.724 | 0.015 | 0.032 | 0.226 | 0.003 | | |
| Mende | 0.003 | 0.881 | 0.058 | 0.058 | 0.001 | | |
| Gumuz | 0.013 | 0.058 | 0.819 | 0.098 | 0.012 | | |
| Oromo/Amhara | 0.013 | 0.058 | 0.098 | 0.819 | 0.012 | | |
| Found 55ka | | | | | | | |
| Nama | 0.677 | 0.052 | 0.264 | | | | 0.005 |
| Mende | 0.021 | 0.562 | 0.272 | | | | 0.143 |
| Gumuz | 0.060 | 0.211 | 0.704 | | | | 0.023 |
| Oromo/Amhara | 0.041 | 0.142 | 0.793 | | | | 0.017 |
| Found 500ka | | | | | | | |
| Nama | | | | | | 0.379 | 0.618 |
| Mende | | | | | | 0.430 | 0.569 |
| Gumuz | | | | | | 0.473 | 0.525 |
| Oromo/Amhara | | | | | | 0.475 | 0.517 |

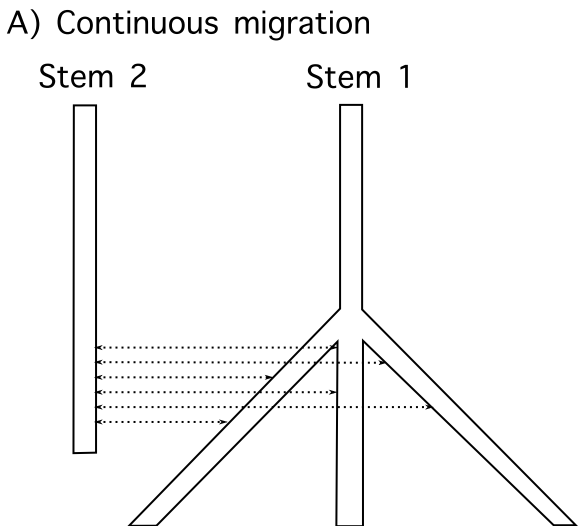

Figure S2: **Two scenarios for population rearrangement.** In our demographic models, we considered A) continuous migration models, in which one of the stem population expands (splits into contemporary populations), and the other stem population(s) has continuous symmetric migration with those populations; and B) one or more of the stem populations expands, with instantaneous pulse (or "merger") events from the other stem population, so that recent populations are formed by mergers of multiple ancestral populations.

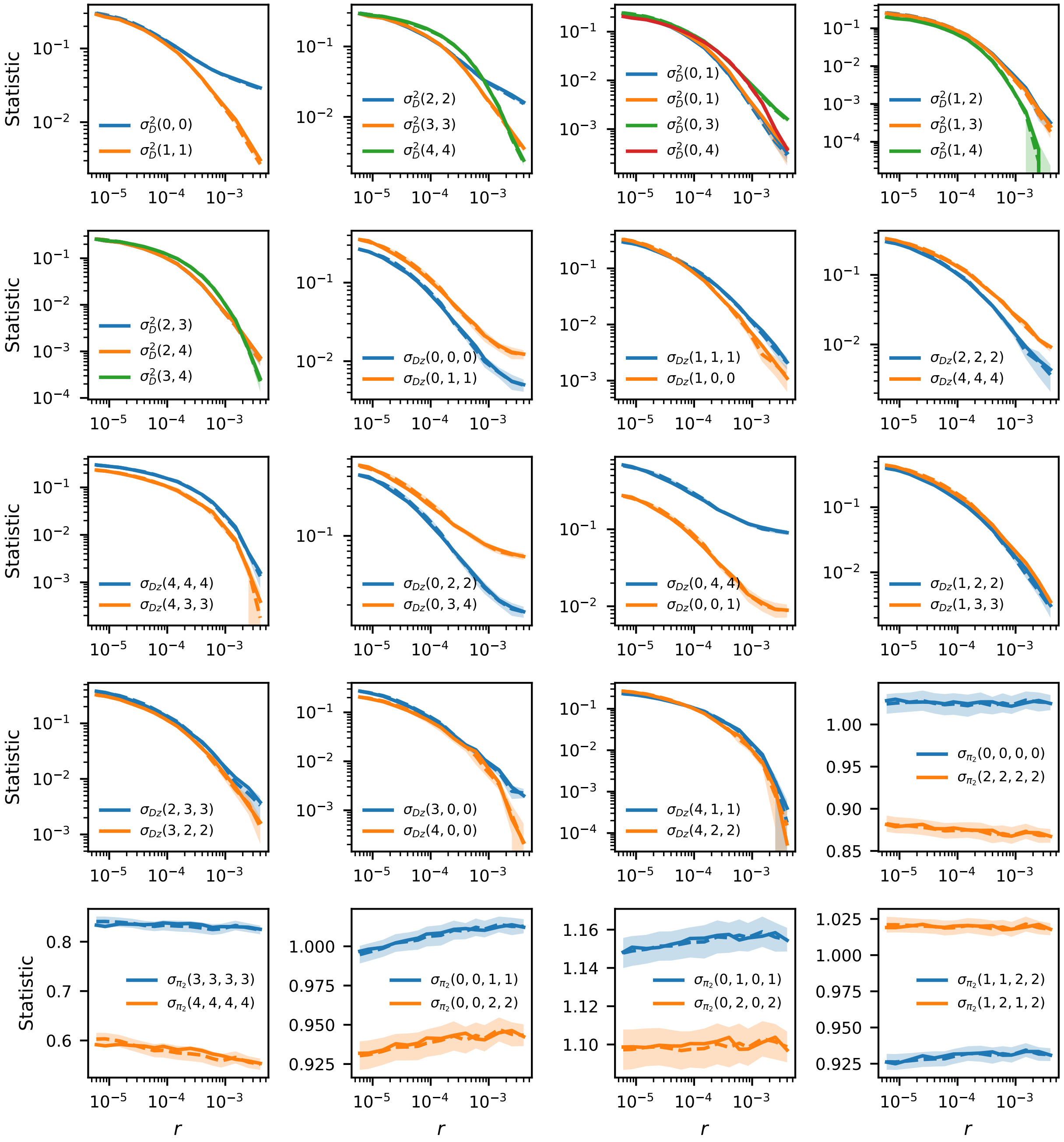

Figure S3: **Comparison of LD statistics parsed using two recombination maps.** The recombination map determines genetic distances between pairs of SNPs, so using different maps could result in systematic biases in parsed statistics if the maps differ significantly. Here, the solid lines are statistics parsed using the HapMapII recombination map, and dashed lines are using the OMNI-YRI map. Shading represents 95% confidence intervals from the OMNI map. These two maps results in LD statistics that are indistinguishable from one another. Notations for each statistic are described in section 2.1 and indexed by populations (0: Nama, 1: Mende, 2: Gumuz, 3: Amhara/Oromo, 4: British).

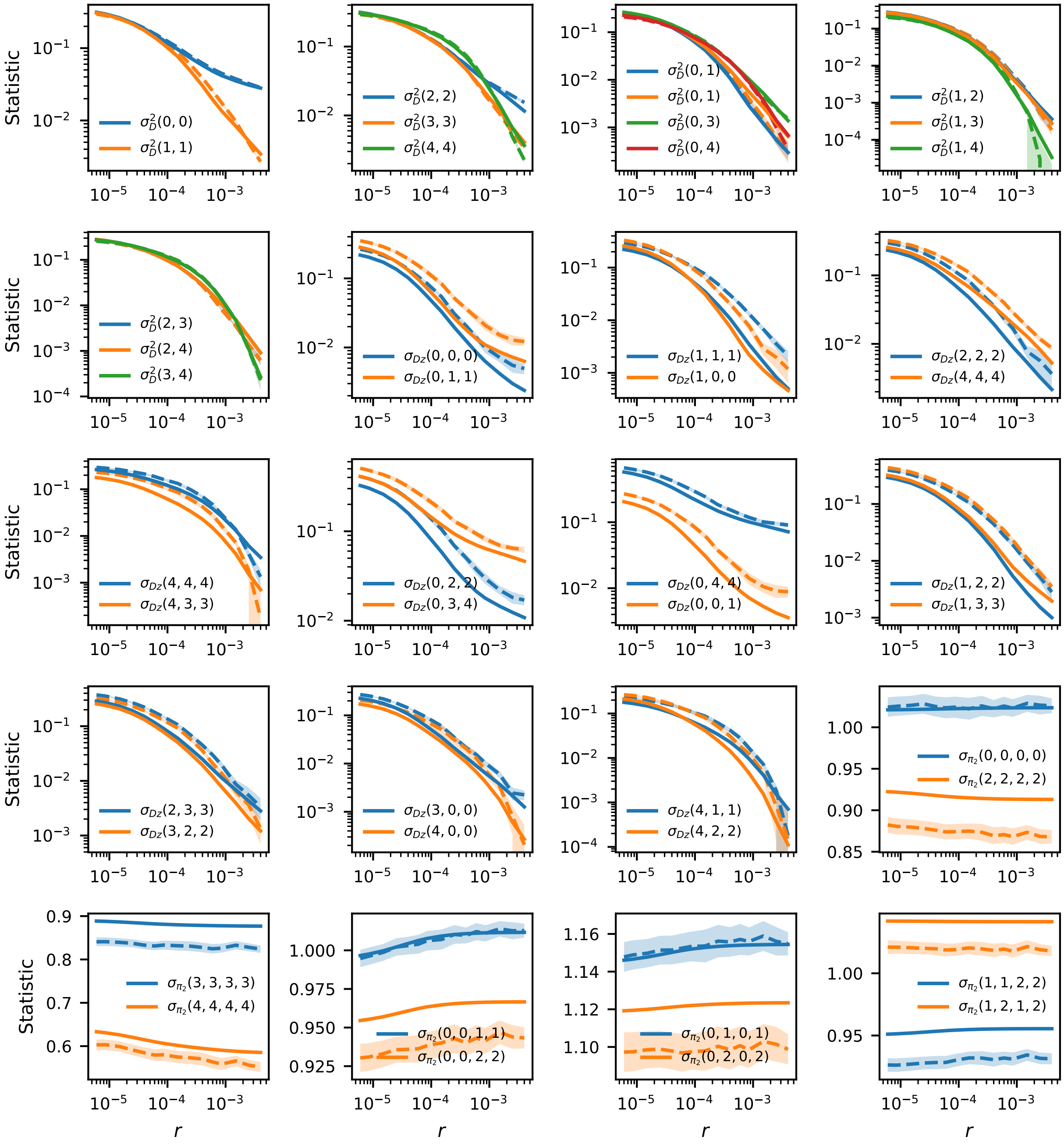

Figure S4: **Single-origin model fit to LD statistics.** Predicted vs observed two-locus statistics as a function of genetic distance for the single-origin model. Each panel represents a different set of two-locus statistics. Solid lines represent estimates using the single-origin model. Dashed lines represent observed statistics. Notation and indexing of statistics are described in the Fig. S3 caption.

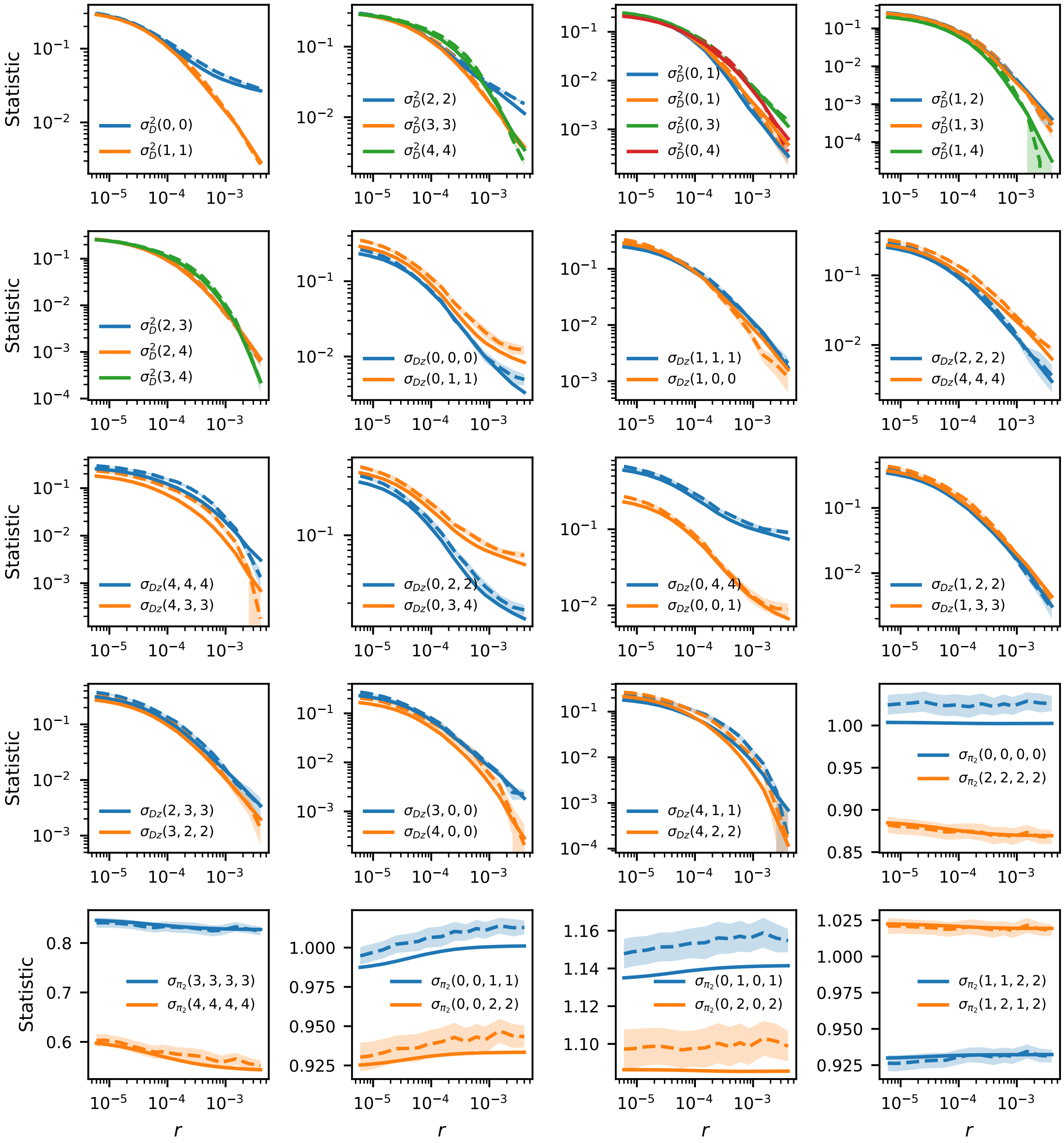

Figure S5: **Continuous-migration model fit to LD statistics.** Predicted vs observed two-locus statistics as a function of genetic distance for the continuous-migration model with migration between stems. Each panel represents a different set of two-locus statistics. Solid lines represent estimates using the continuous-migration model. Dashed lines represent observed statistics. Notation and indexing of statistics are described in the Fig. S3 caption.

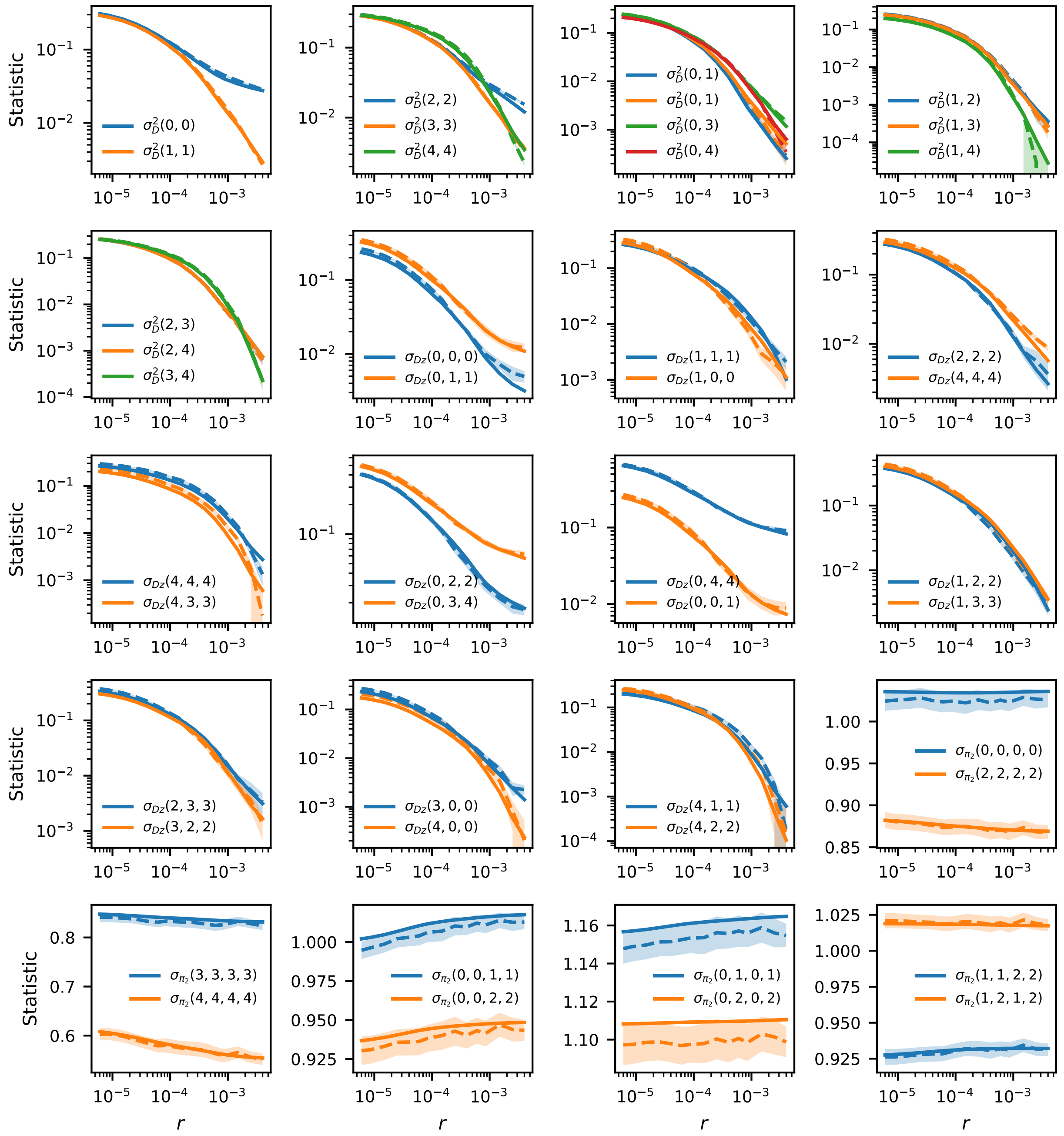

Figure S6: **Merger-without-stem-migration model fit to LD statistics.** Predicted vs observed two-locus statistics as a function of genetic distance for the merger-without-stem-migration model. Each panel represents a different set of two-locus statistics. Solid lines represent estimates using the merger-without-stem-migration model. Dashed lines represent observed statistics. Notation and indexing of statistics are described in the Fig. S3 caption.

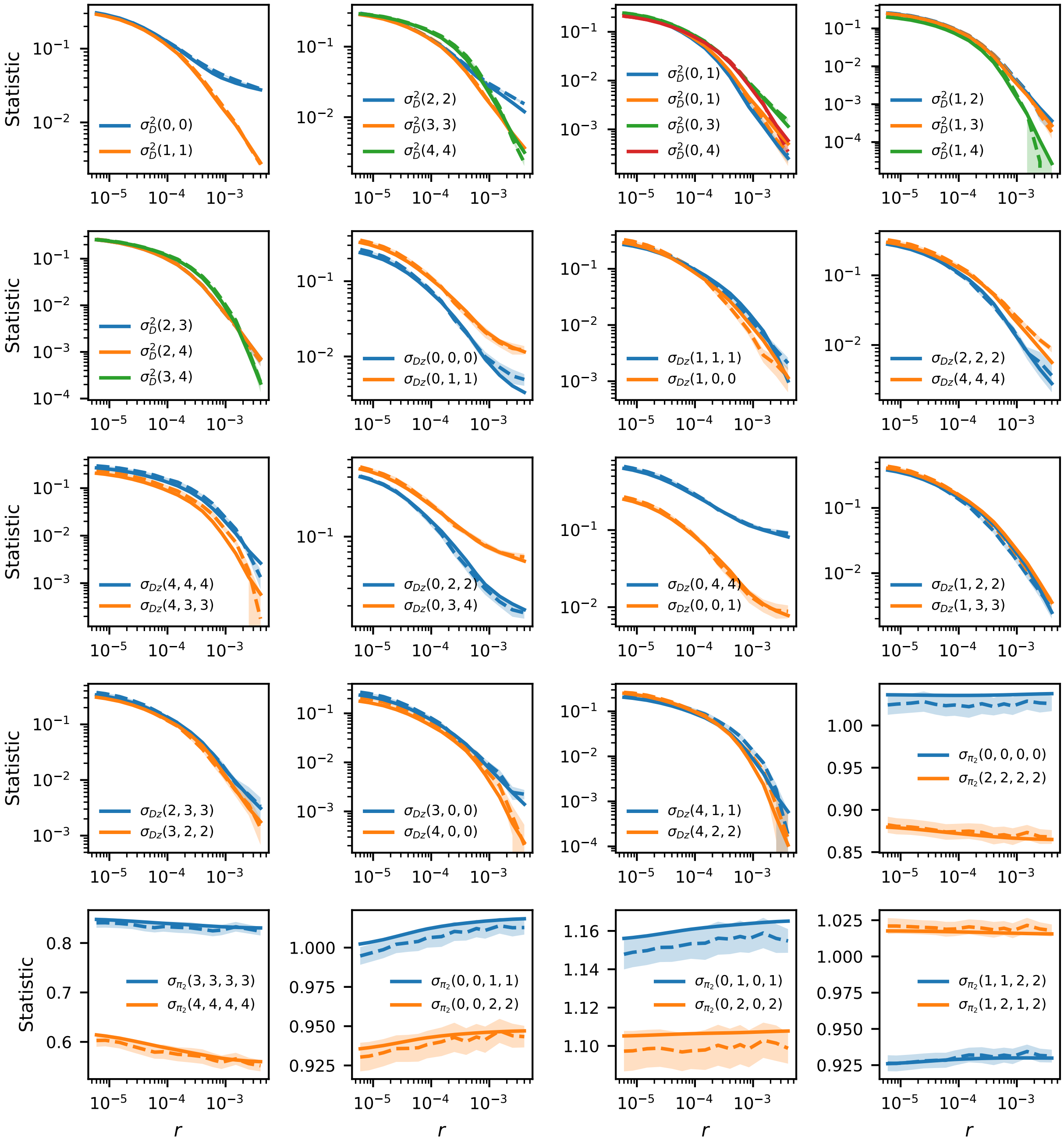

Figure S7: **Merger-with-stem-migration model fit to LD statistics.** Predicted vs observed two-locus statistics as a function of genetic distance for the merger-with-stem-migration model. Each panel represents a different set of two-locus statistics. Solid lines represent estimates using the merger-with-stem-migration model. Dashed lines represent observed statistics. Notation and indexing of statistics are described in the Fig. S3 caption.

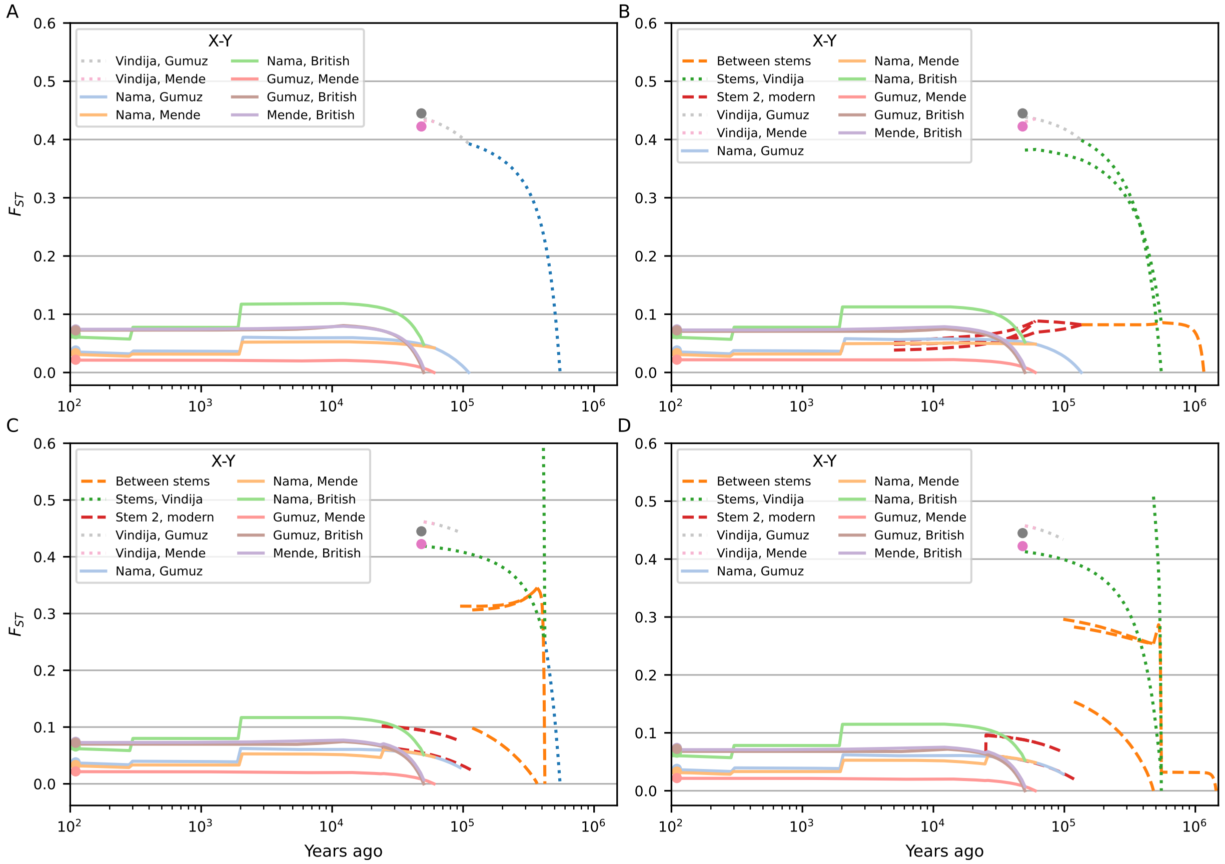

Figure S8: **Predicted $F_{ST}$ between contemporaneous populations over time.** All models (A: single origin, B: continuous migration, C: merger without stem migration, D: merger with stem migration) match observed $F_{ST}$ between sampled populations. The continuous-migration model predicts that human stem populations remain genetically similar, while the merger with and without stem migration models predict a period of increased $F_{ST}$ between ancestors of contemporary humans. This is largely due to the very small inferred $N_e$ in one of the stem branches, which leads to a rapid increase in differentiation due to drift.

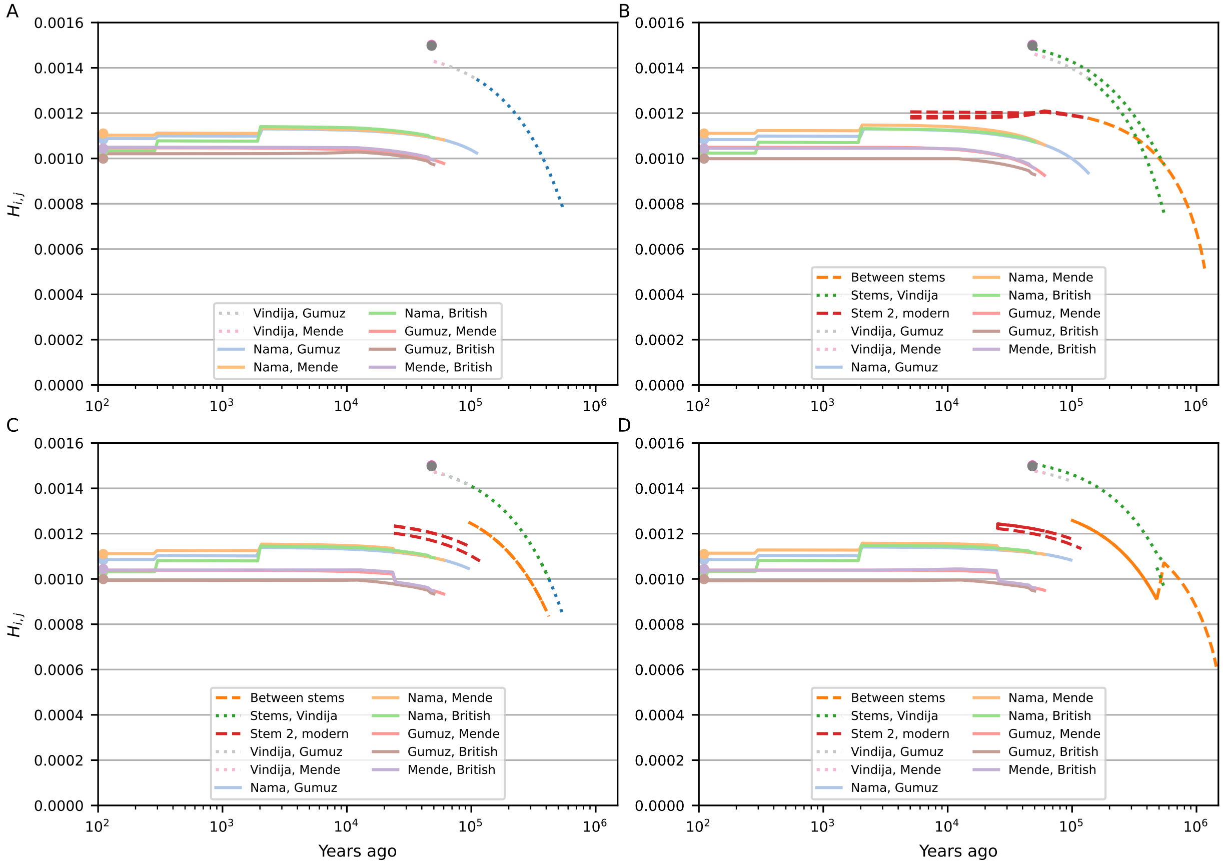

Figure S9: **Predicted pairwise differentiation between contemporaneous populations.** $H_{i,j}$ is the predicted pairwise differentiation between two populations per base-pair, using a mutation rate estimated from matching predicted to observed pairwise diversity from the intergenic data used in the fits (see Sec. 7.2). The single-origin model (A) shows larger deviations between observed and expected differentiation than the models with early human structure (B: continuous migration, C: merger without stem migration, D: merger with stem migration). In the merger models (C and D), while $F_{ST}$ is large between early stems due to the sharp bottleneck in one of those branches, the pairwise differences are comparable to the continuous-migration model. Stem 1E and Stem 1S have low differentiation, as the bottleneck in Stem 1 (which they both split from) sharply reduces diversity by the time of the split. This has the effect of large $F_{ST}$ but small $H_{i,j}$.

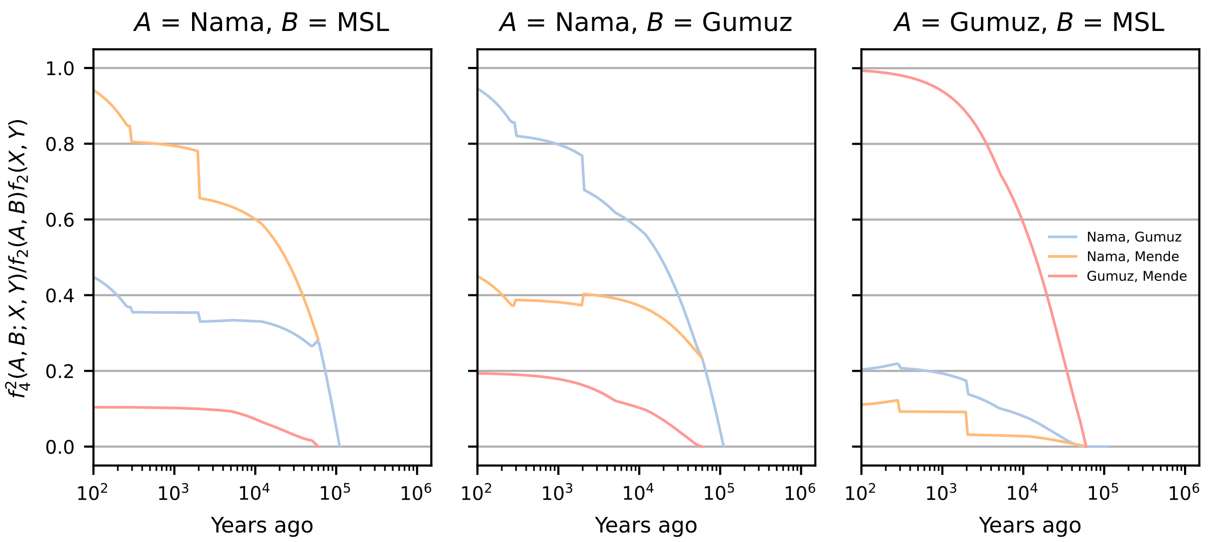

Figure S10: **Predicted $f_4$ between pairs of contemporary human populations and ancient branches from the single-origin model.** $f_4^2(A, B; X, Y)/f_2(A, B)f_2(X, Y)$ is interpreted as the proportion of the amount of drift between sampled populations $A$ and $B$ that aligns with drift between populations $X$ and $Y$, sampled in the past (see Section 5.2). In the single-origin model, nearly all differentiation between the Nama, Mende, and Gumuz populations arises after the divergence events, so normalized $f_4$ decays to zero by the time of those divergences.

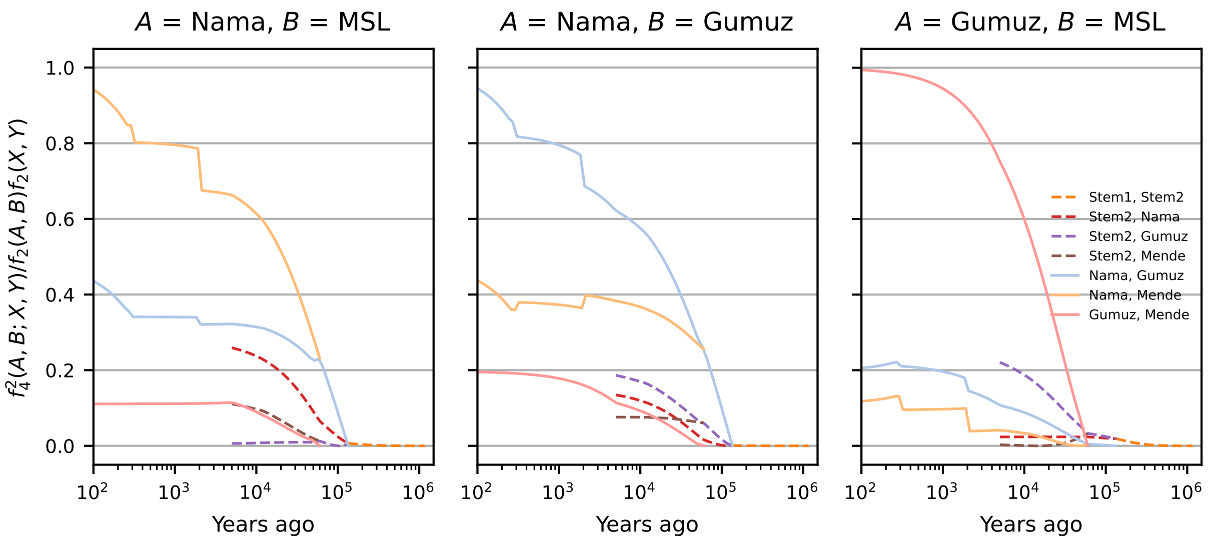

Figure S11: **Predicted $f_4$ between pairs of contemporary human populations and ancient branches from the continuous-migration model.** Despite population structure inferred to have extended up to 1Ma into the past, differentiation between contemporary African populations only traces back to differentiation between ancestral populations ∼100-200ka, indicating that while stem populations were structured (Fig. S8), differentiation due to that ancient structure contributes little to differentiation between contemporary populations.

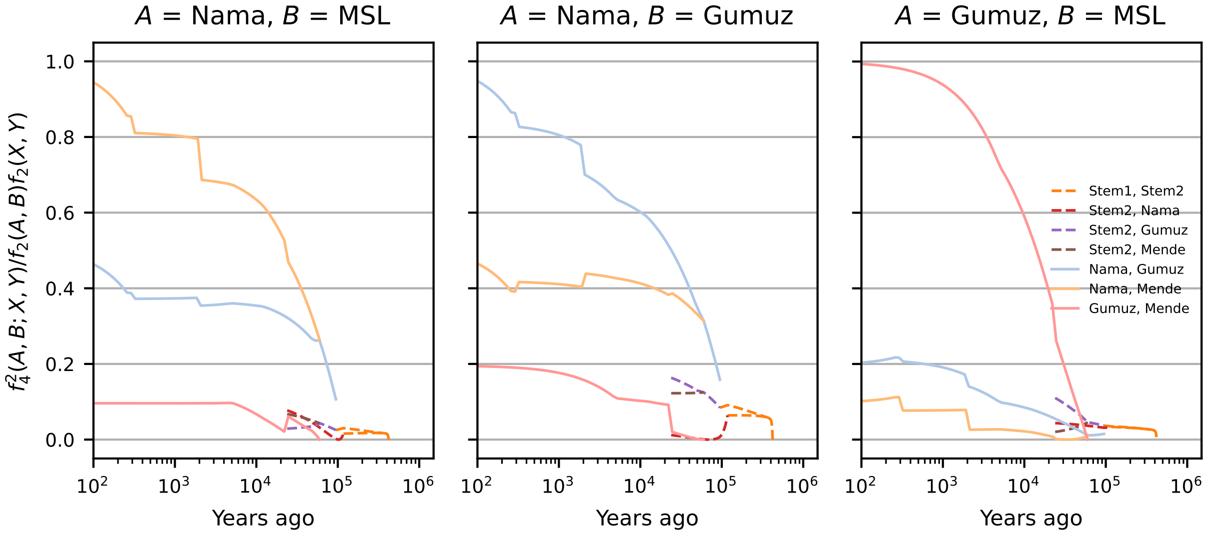

Figure S12: **Predicted $f_4$ between pairs of contemporary human populations and ancient branches from the merger-without-stem-migration model.** Unlike the continuous-migration model, population structure in the stems following the divergence of Neanderthal and human ancestors results in differentiation among contemporary populations that aligns with drift between the stems. In this model, migration between stem populations is disallowed.

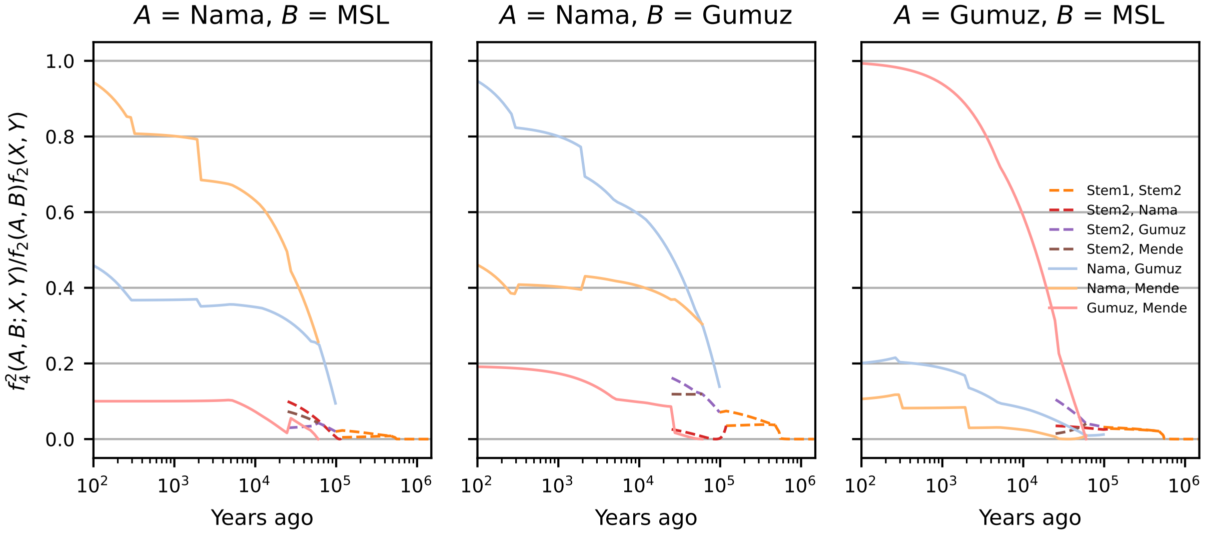

Figure S13: **Predicted $f_4$ between pairs of contemporary populations and ancient branches from the merger-with-stem-migration model.** Differentiation between present-day populations aligns with up to ≈ 5% of the drift between stems, although some population pairs show even less shared drift.

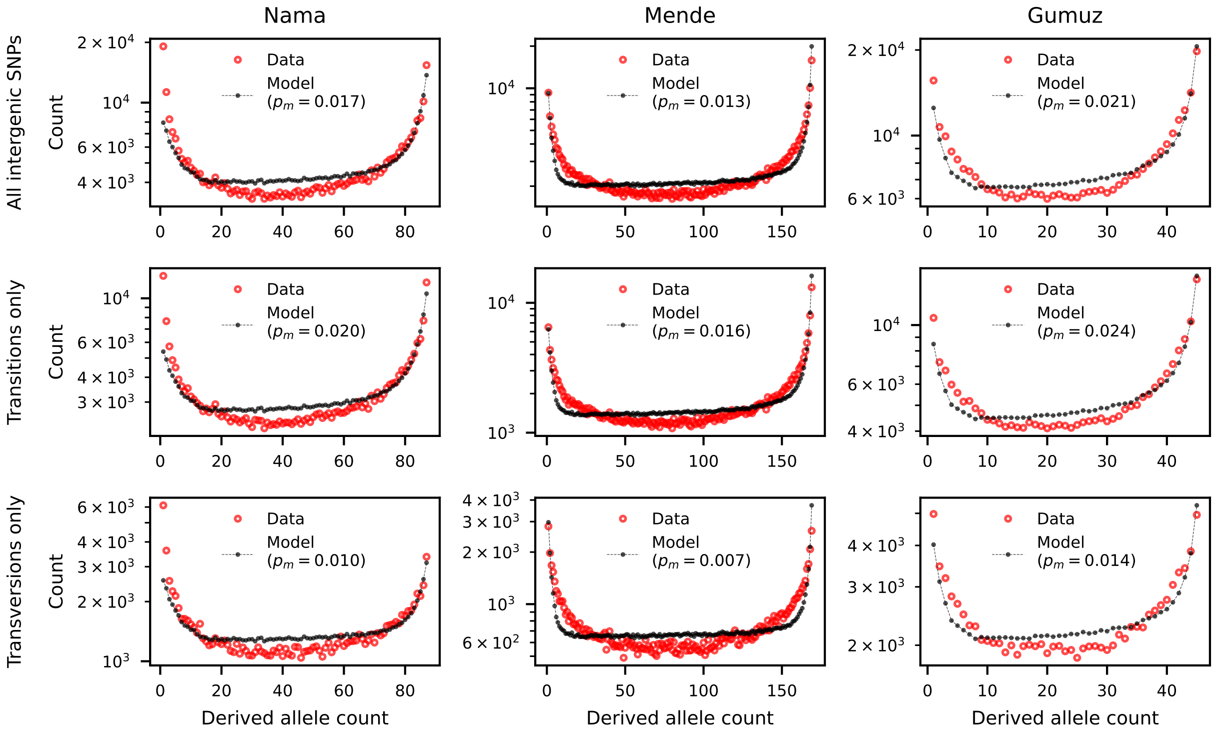

Figure S14: **Conditional SFS compared to single-origin model predictions.** The single-origin model provides a poor fit to the Neanderthal-conditional SFS in the Mende, Gumuz, and Nama (see Section 7.1.1). Ancestral state misidentification (labeled as $p_m$ in the figure legends) was inferred to be between 1 and 2%, and misidentification rates were consistent across model comparisons. Misidentification rates of transition-type mutations were roughly double that of transversion-type mutations, consistent with the higher mutation rates of transition mutations vs. transversions.

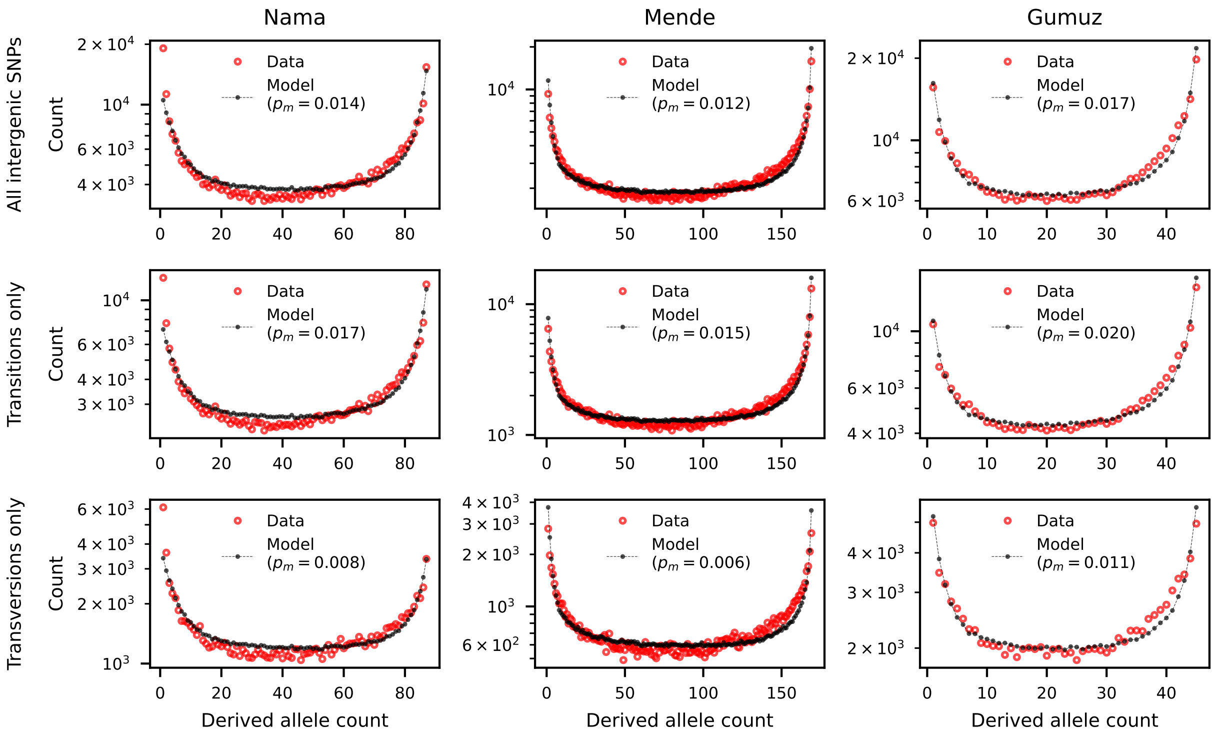

Figure S15: **Conditional SFS compared to continuous-migration model predictions.** The continuous-migration model provides a better fit to the Neanderthal-conditional SFS than either the single-origin model or the merger-without-stem-migration model, although systematic deviations are still noticeable in each population. Ancestral state misidentification ($p_m$), as estimated with all models, is roughly doubled for transition-type mutations compared to transversions.

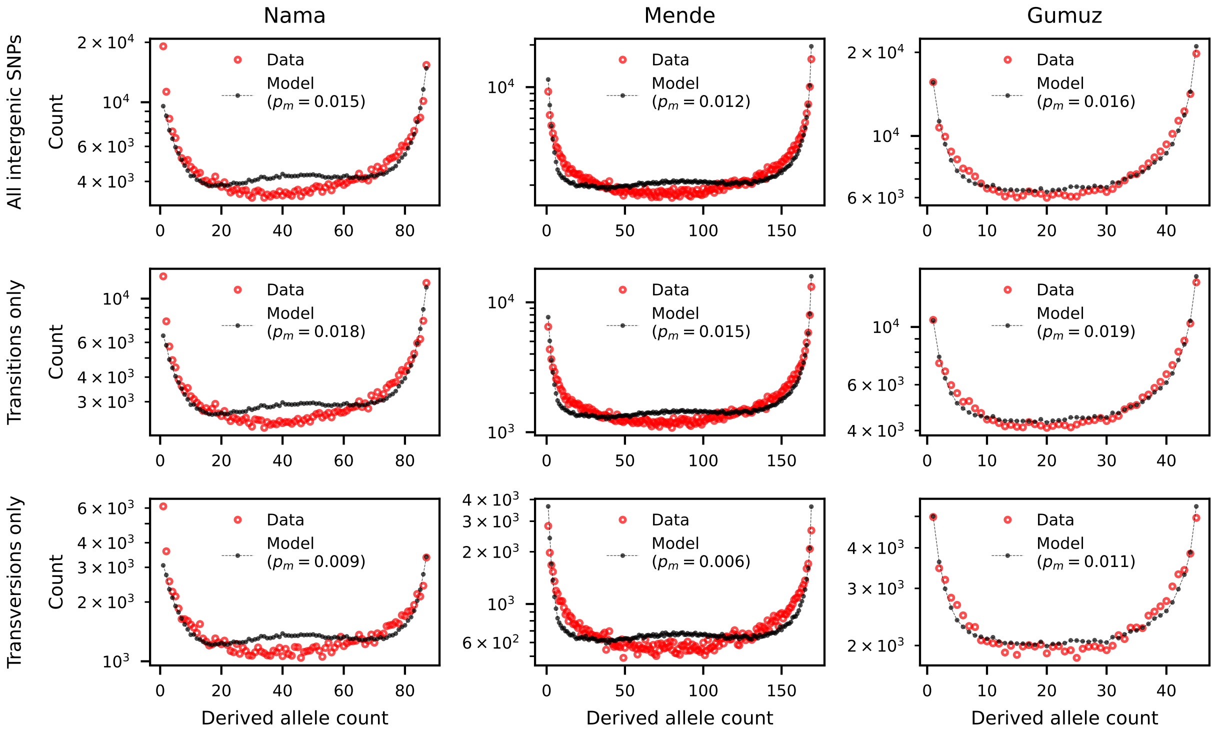

Figure S16: **Conditional SFS compared to merger-without-stem-migration model predictions.** The excess of intermediate-frequency variants in the Nama and Mende cSFS results in a poorer fit to the cSFS compared to the continuous-migration and merger-with-stem-migration models. Thus the model without stem migrations provides a poorer fit to both two-locus statistics and the cSFS. Ancestral state misidentification for transition-type mutations ($p_m$), as estimated with all models, is roughly double that of transversions.

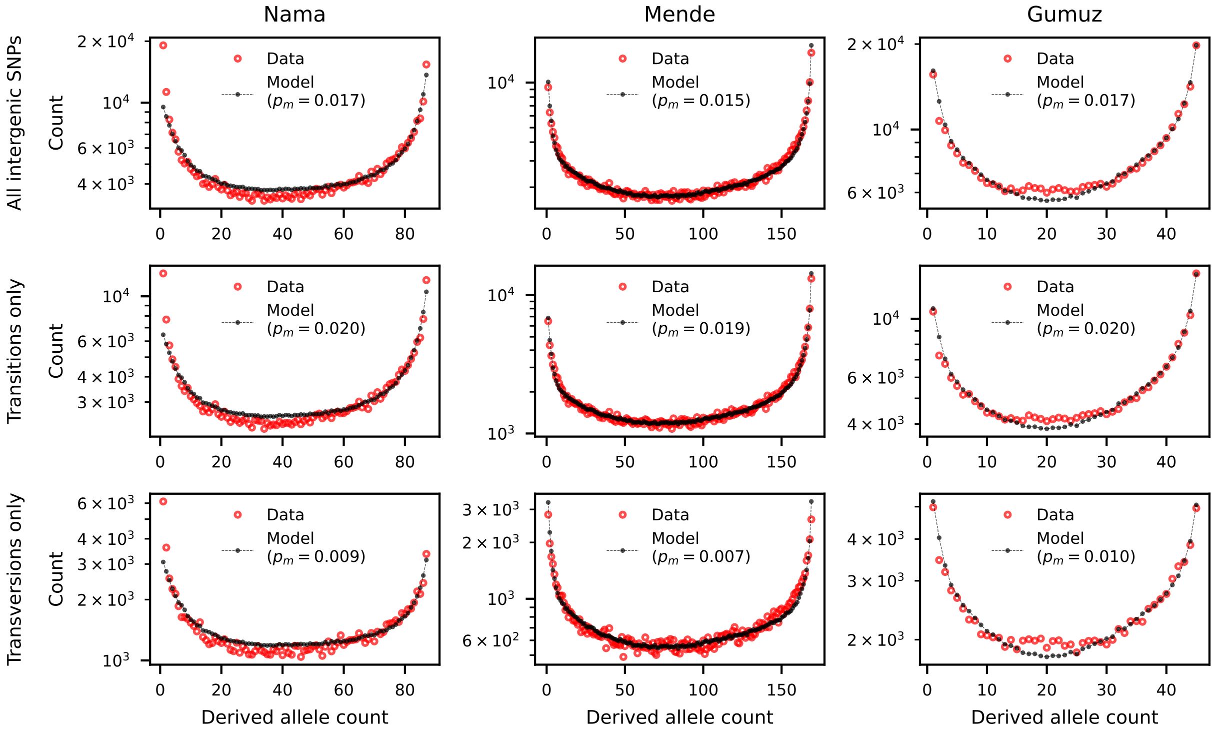

Figure S17: **Conditional SFS compared to merger-with-stem-migration model predictions.** The merger-with-stem-migration model, in addition to providing the best fit to heterozygosity and LD statistics, provided the best fit to the cSFS in the Nama, Gumuz, and Mende populations. Rates of ancestral state misidentification were consistent across each comparison, with the ancestral state of transversion mutations estimated to be $0.7-1.0\%$, and that of transition mutations estimated to be $1.9-2.0\%$. These misidentification rates are consistent with the known difference in mutation rates between transitions and transversions, as well as other SFS-based inferences in humans using the Thousand Genomes data and the 6-primate alignment (see Methods).

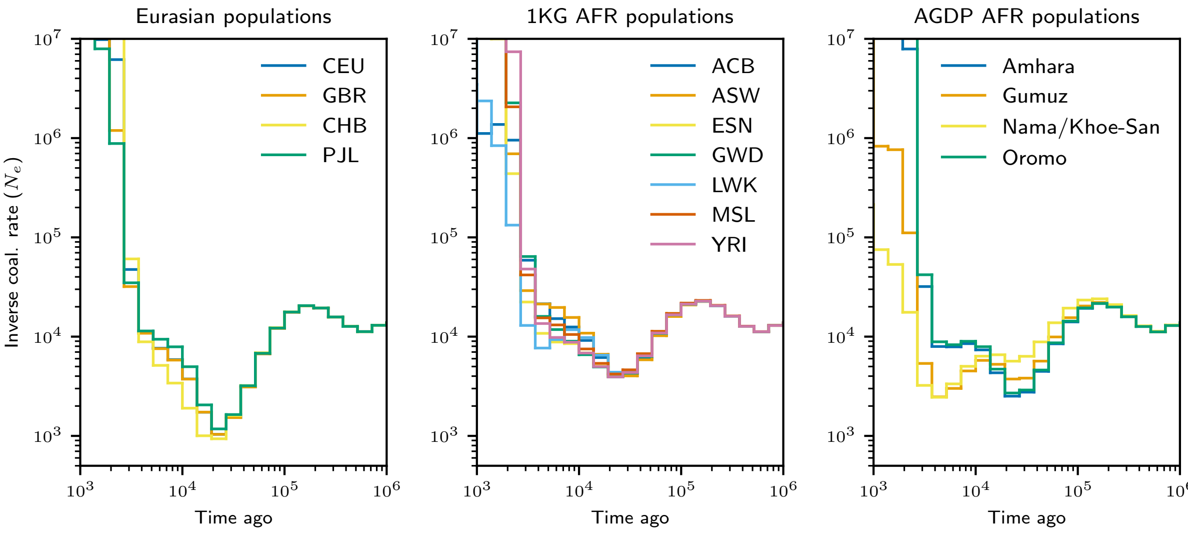

Figure S18: **Inverse instantaneous coalescence rates inferred from the data.** Using combined data from the Thousand Genomes Project and the AGRP, we reconstructed gene genealogies using Relate and calculated the inverse instantaneous coalescence rate for samples within each population. (A) Eurasian populations (CEU: European American, GBR: British, CHB: Han Chinese, PJL: Punjabi), (B) Thousand Genomes African populations (ACB: Afro-Caribbean, ASW: African American, ESN: Esan, GWD: Gambian, LWK: Luhya, MSL: Mende, YRI: Yoruba), (C) AGRP populations.

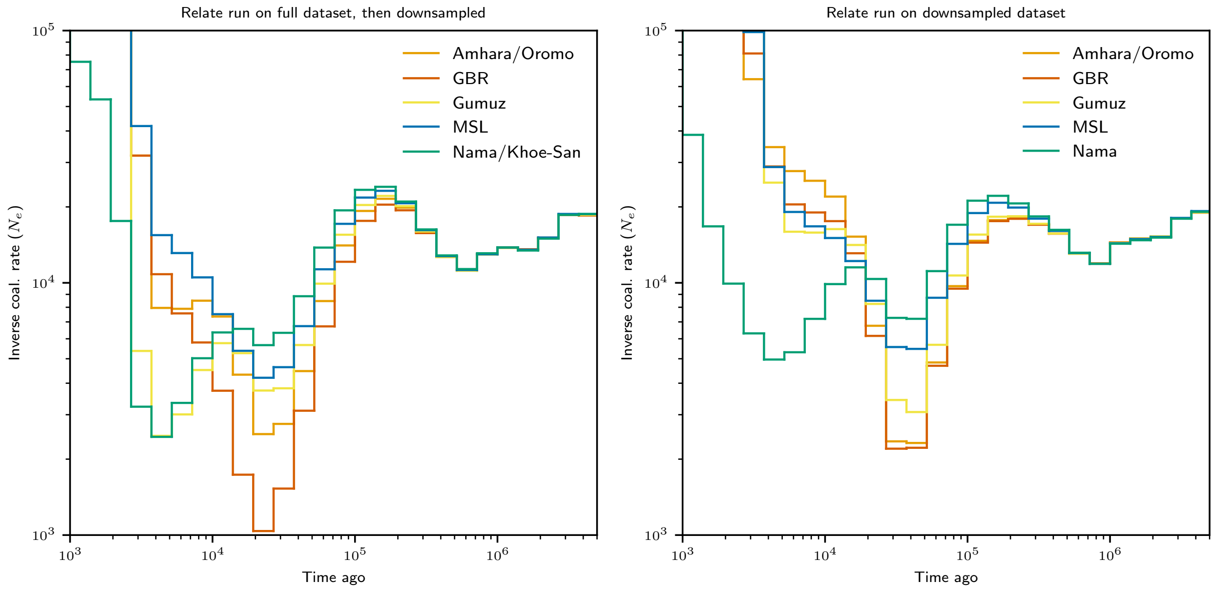

Figure S19: **Inverse instantaneous coalescence rates inferred from the data, subset to focal populations in the model inferences.** Fig. S18 shows IICR curves from reconstructed genealogies inferred using a larger dataset (including nearly 1,200 individuals) than the subset of populations we primarily focused on here. To assess the robustness of Relate-inferred genealogies and for a direct comparison to our inferred demographic models, we ran Relate on the 290 genomes from the Nama, Gumuz, Mende, Amhara/Oromo, and British. While the reconstructed IICR curves are qualitatively similar to those from the full dataset for these populations in the distant past, there are noticeable discrepancies over the recent history in the last tens of thousands of years. Left: Relate run on full dataset, showing focal populations. Right: Relate run directly on the subset dataset of focal populations.

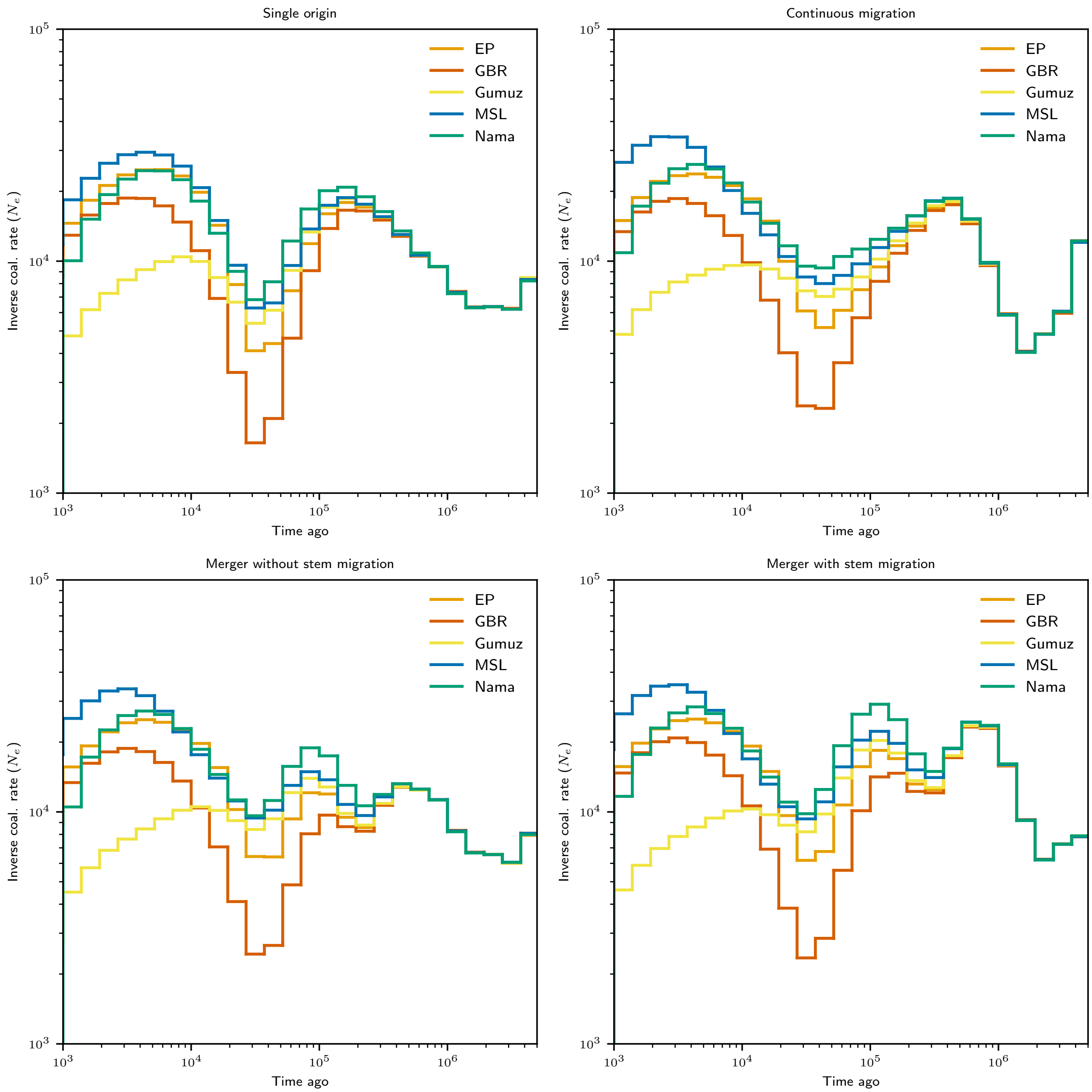

Figure S20: **Inverse instantaneous coalescence rates reinferred from simulated data.** We simulated data under our four highlighted models, sampling the same number of individuals per population as the original dataset, and then ran Relate on each simulated dataset to reconstruct IICR curves for each. Each model has qualitatively different histories, particularly for the distant past with varying numbers and timings of oscillations. However, due to the discrepancies in reconstructed genealogies using even the same data as input (Figs. S19 and S22), it is difficult to draw firm conclusions from quantitative comparisons between reconstructed genealogies.

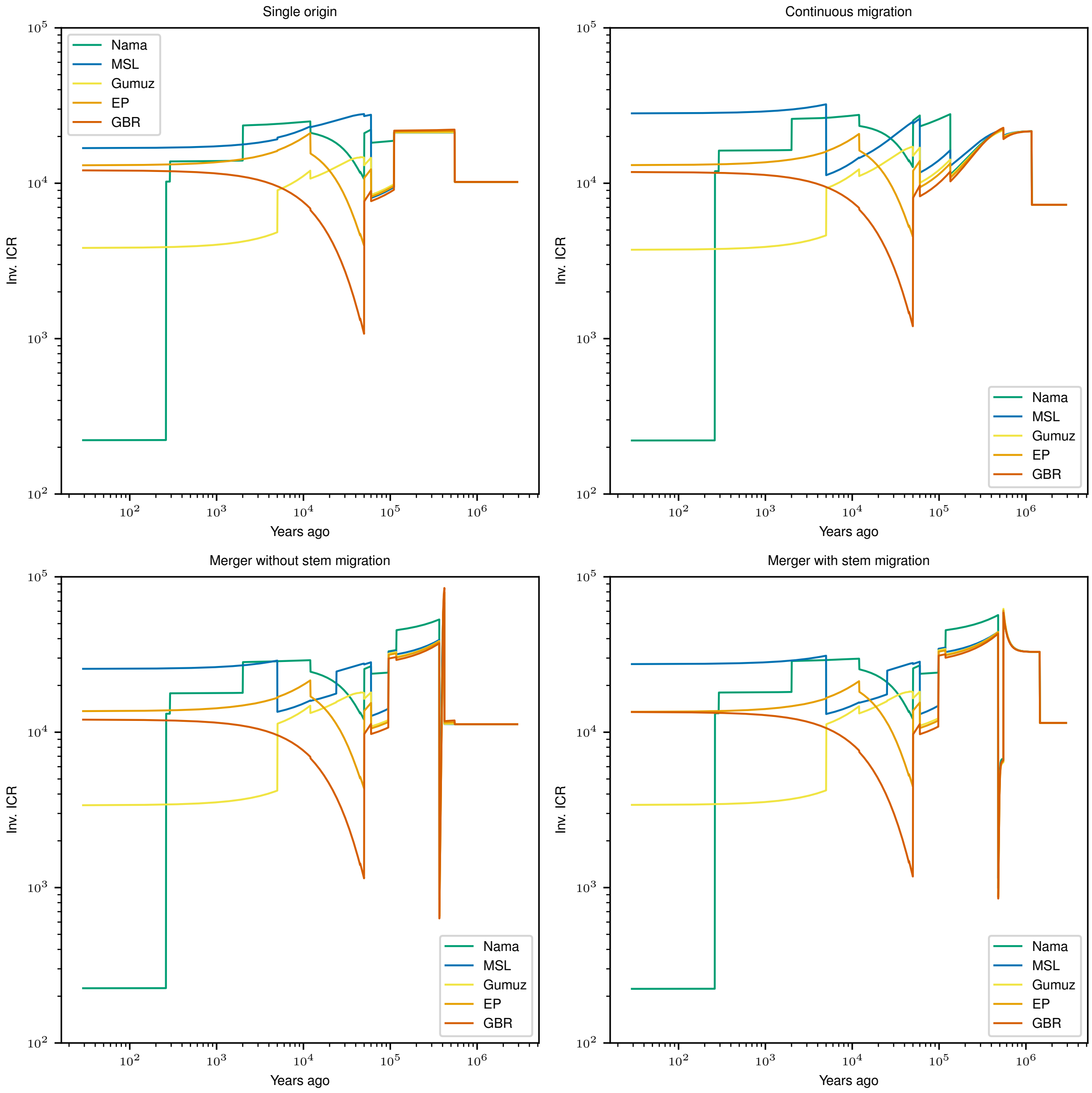

Figure S21: **Expected inverse instantaneous coalescence rates from models.** The inferred demographic models specify population sizes, split times, and migration rates and events, making it possible to compute the exact expected IICR curve for each population. Due to discrete events (instantaneous size changes, for example), the models can produce non-smooth IICR curves, although each predicts a period of increased "effective size" in the past $\sim 200 - 500$ka.

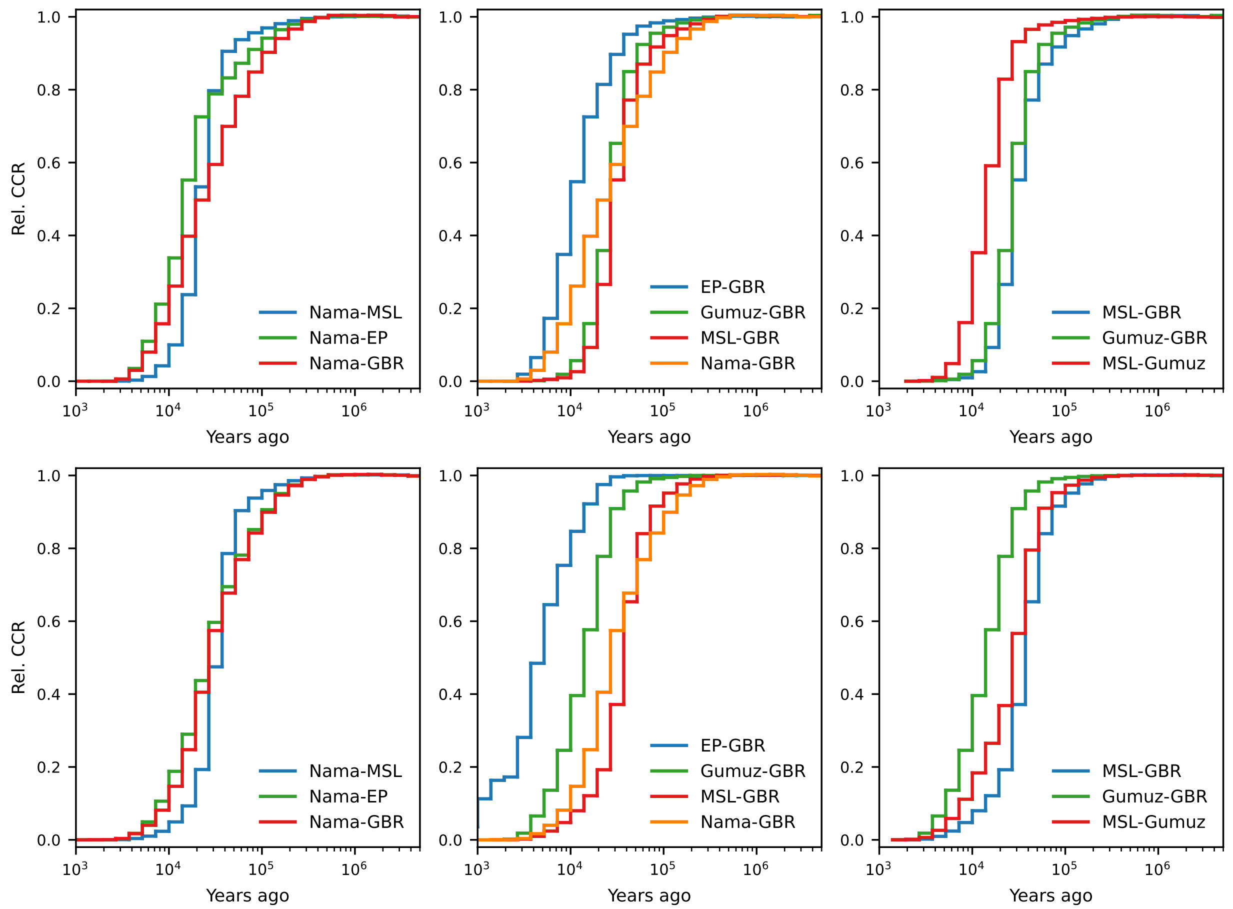

Figure S22: **Relative cross-coalescence rates inferred from the data.** Using the same reconstructed genealogies from Relate as displayed in Fig. S19, we computed relative cross-coalescence (RCCR) curves for pairs of populations using genealogies from the either the full dataset ($\sim 1,200$ individuals, top) and the subset of focal populations (290 individuals, bottom). Despite the overlap in samples, the two sets of inferred genealogies provided inconsistent RCCR, with both the timing and ordering of increased RCCR varying between the two datasets. Genealogies reconstructed from the full dataset provide a better match to each of inferred demographic models (Figs. S23–S26).

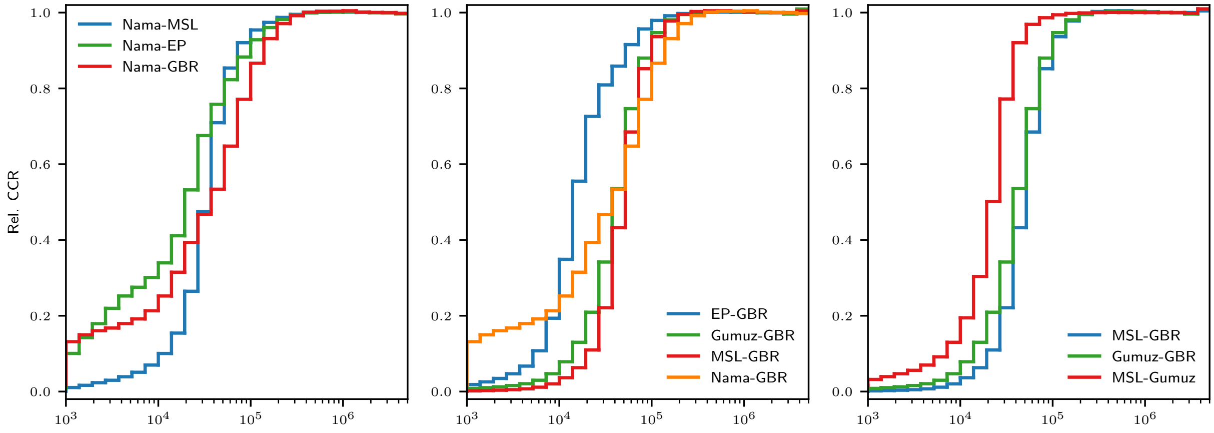

Figure S23: **Relative cross-coalescence rates reinferred from simulated data under the single-origin model.** RCCR curves from genealogies reconstructed from simulated data using Relate match those inferred from the full dataset (S22, top) for distant time points. There are large discrepancies in the very recent history, where Relate and other genome-wide coalescence-based methods are known to be underpowered.

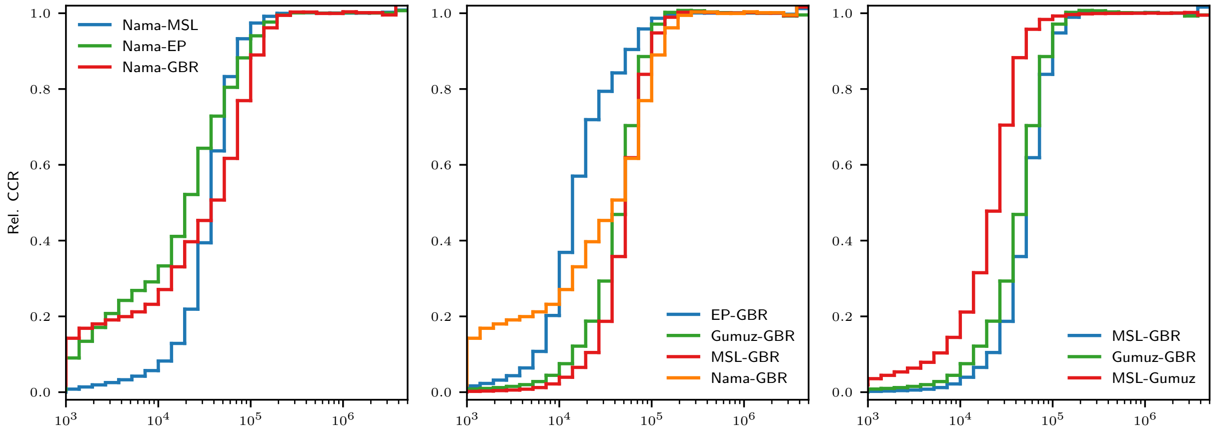

Figure S24: **Relative cross-coalescence rates reinferred from simulated data under the continuous-migration model.** RCCR curves from genealogies reconstructed from simulated data using Relate match those inferred from the full dataset (S22, top) for distant time points. There are large discrepancies in the very recent history, where Relate and other genome-wide coalescence-based methods are known to be underpowered.

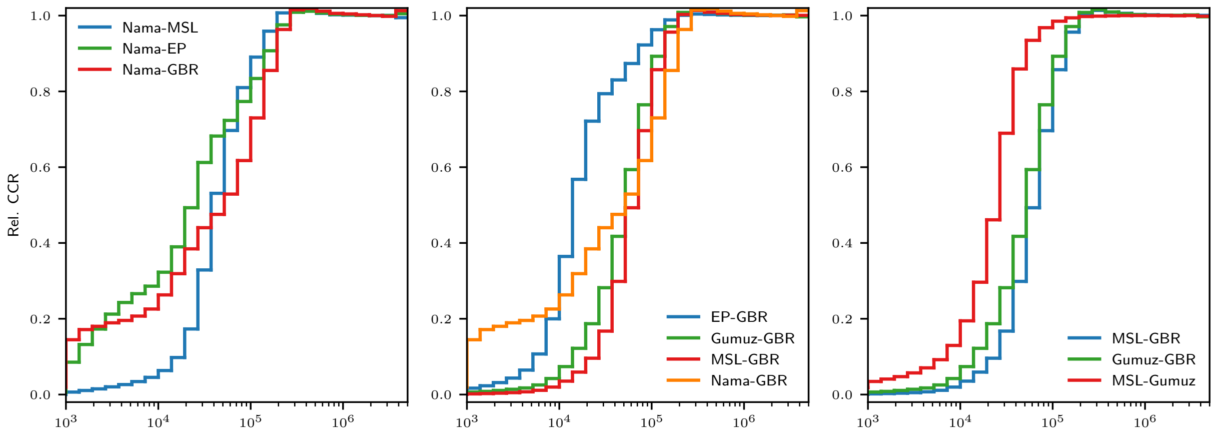

Figure S25: **Relative cross-coalescence rates reinferred from simulated data under the merger-without-stem-migration-model.** RCCR curves from genealogies reconstructed from simulated data using Relate match those inferred from the full dataset (S22, top) for distant time points. There are large discrepancies in the very recent history, where Relate and other genome-wide coalescence-based methods are known to be underpowered.

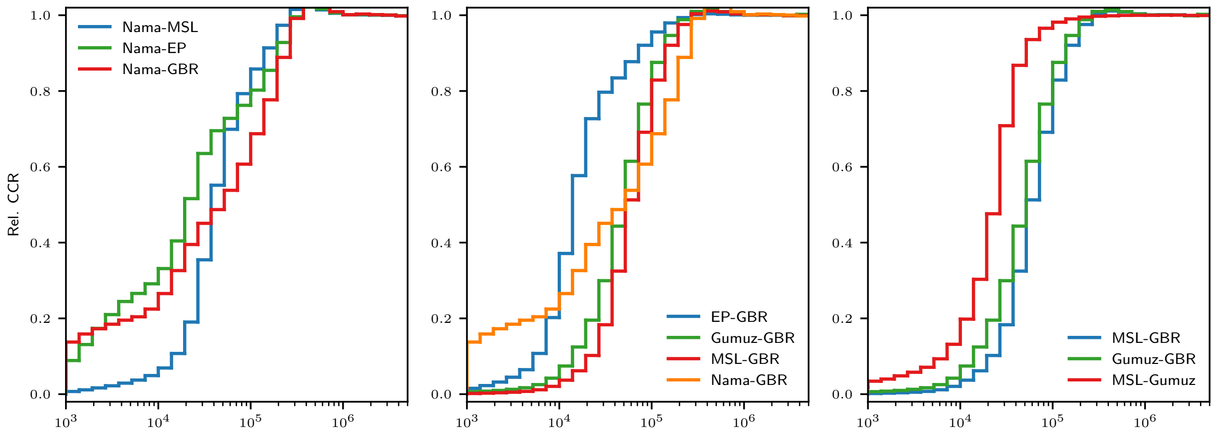

Figure S26: **Relative cross-coalescence rates reinferred from simulated data under the merger-with-stem-migration model.** RCCR curves from genealogies reconstructed from simulated data using Relate match those inferred from the full dataset (S22, top) for distant time points. There are large discrepancies in the very recent history, where Relate and other genome-wide coalescence-based methods are known to be underpowered.

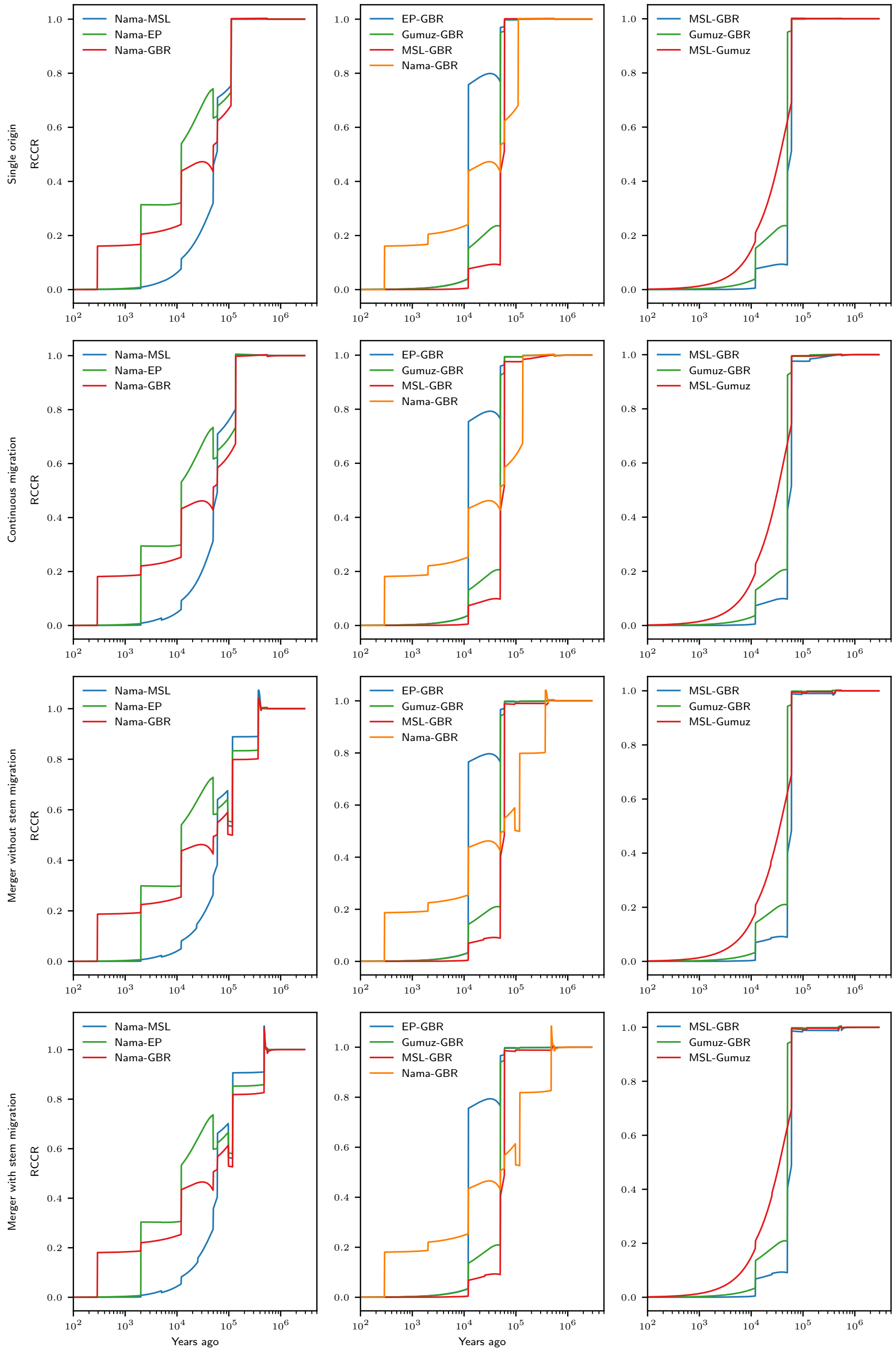

Figure S27: **Expected relative cross-coalescence rates from inferred models.** Our inferred demographic models allow for exact calculation of expected RCCR curves, showing that while the parameterizations of the four models are quite different, they each predict very similar RCCR profiles.

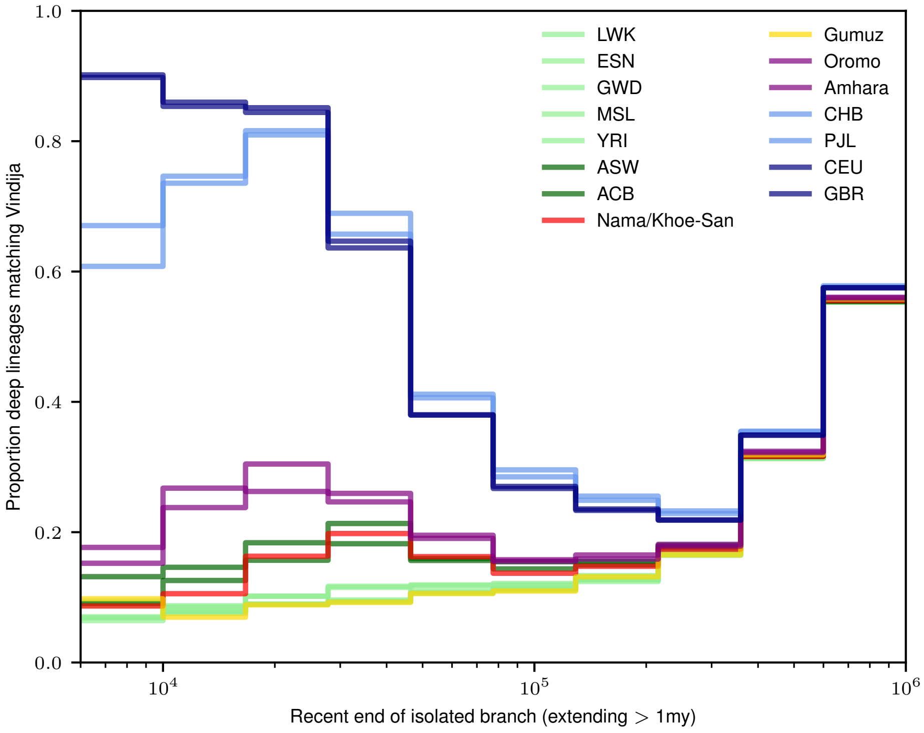

Figure S28: **Neanderthal-matching deep branch distribution from the data.** Following the approach of Speidel *et al.* (2019), we categorized Neanderthal-matching deep branches for each population from Relate-inferred gene genealogies. The proportion of deep branches with Neanderthal affinity is highest for Eurasian populations, as expected, and higher for African populations with recent Eurasian ancestry (Oromo, Amhara, ACB, ASW, and Nama) than the Gumuz and West African populations.

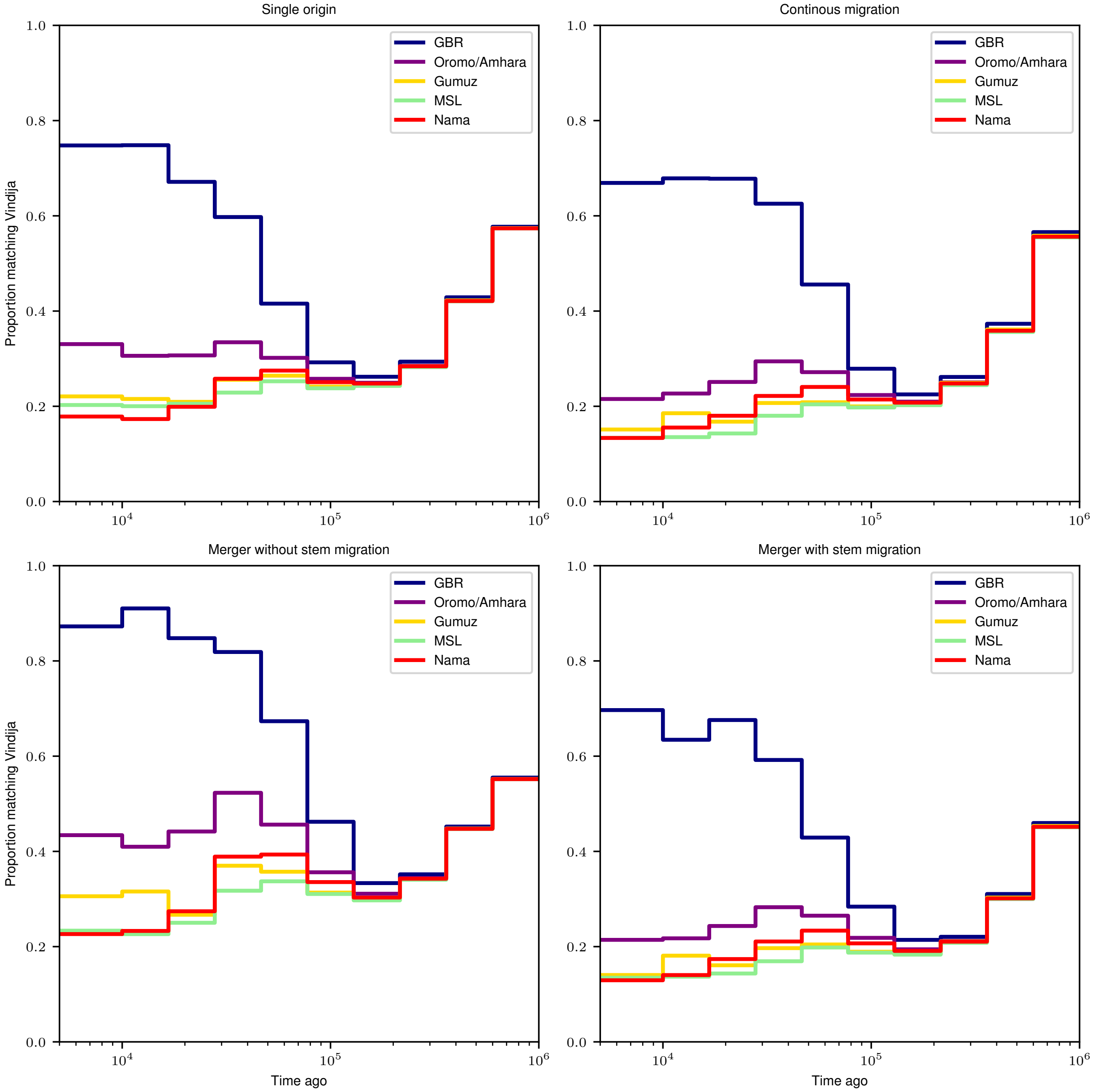

Figure S29: **Deep branch distribution reinferred from simulated data.** We used Relate to reinfer gene genealogies from simulated data under the four highlighted models and categorized Neanderthal-matching deep branches in the simulated datasets. Each model provided a qualitative match to the data (highest proportion of Neanderthal-matching deep branches in the British, followed by Amhara/Oromo and Nama). Similar to reinferred IIRC and RCCR, no model provided a perfect match to the reconstructions using Relate with the data, but the similarities between reconstructed genealogies in the simulated data suggest such coalescence-rate based curves are underpowered to differentiate between plausible models of history.

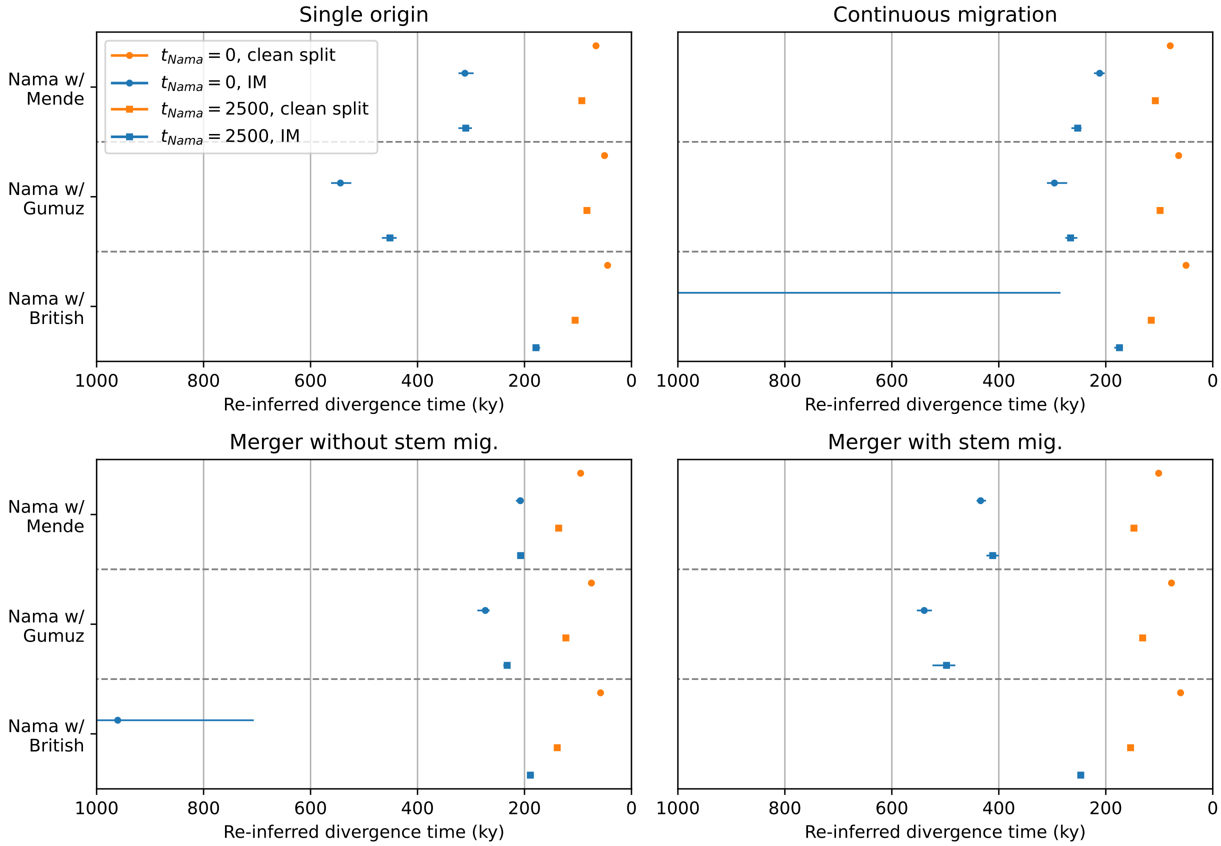

Figure S30: **IM model split times reinferred from simulated data of full model.** We simulated 10 diploid individuals under the four highlighted models (Sec. 7.3). Using the joint-SFS, we reinferred a simpler isolation-with-migration (IM) model for pairs of populations including the Nama to explore the effect of model misspecification on the earliest inferred divergence times between Khoe-San and other populations. We simulated data with Nama individuals sampled at present ($t = 0$) as well as Nama individuals sampled 2,500 years ago, prior to the recent gene flow from East Africans and Europeans. We fit two models for each model and population pair: one that allowed for migration between the populations (IM), and one that disallowed migration (clean split). Despite all original models having recent population divergences $\sim$ 120ka, the reinferred IM models all settled on divergence times greater than 200ka and sometimes much greater. If the true historical model includes reticulation or long-lasting structure, simpler IM models may be severely biased in favor of more ancient divergence times.
